# PRDM9 drives the location and rapid evolution of recombination hotspots in salmonids

**DOI:** 10.1101/2024.03.06.583651

**Authors:** Marie Raynaud, Paola Sanna, Julien Joseph, Julie Clément, Yukiko Imai, Jean-Jacques Lareyre, Audrey Laurent, Nicolas Galtier, Frédéric Baudat, Laurent Duret, Pierre-Alexandre Gagnaire, Bernard de Massy

## Abstract

In many eukaryotes, meiotic recombination occurs preferentially at discrete sites, called recombination hotspots. In various lineages, recombination hotspots are located in regions with promoter-like features and are evolutionarily stable. Conversely, in some mammals, hotspots are driven by PRDM9 that targets recombination away from promoters. Paradoxically, PRDM9 induces the self-destruction of its targets and this triggers an ultra-fast evolution of mammalian hotspots. PRDM9 is ancestral to all animals, suggesting a critical importance for the meiotic program, but has been lost in many lineages with surprisingly little effect on meiosis success. However, it is unclear whether the function of PRDM9 described in mammals is shared by other species. To investigate this, we analyzed the recombination landscape of several salmonids, the genome of which harbors one full-length PRDM9 and several truncated paralogs. We identified recombination initiation sites in *Oncorhynchus mykiss* by mapping meiotic DNA double-strand breaks (DSBs). We found that DNA DSBs clustered at hotspots positioned away from promoters, enriched for the H3K4me3 and H3K4me36 marks and the location of which depended on the genotype of full-length *Prdm9*. We observed a high level of polymorphism in the zinc finger domain of full-length *Prdm9*, but not of the truncated paralogs. Moreover, population-scaled recombination maps in *O. mykiss*, *Oncorhynchus kisutch* and *Salmo salar* revealed a rapid turnover of recombination hotspots caused by PRDM9 target motif erosion. Our results imply that PRDM9 function is conserved across vertebrates and that the peculiar evolutionary runaway caused by PRDM9 has been active for several hundred million years.

## Introduction

Meiotic recombination (*i.e.* the exchange of genetic material between homologous chromosomes during meiosis) is highly conserved in a wide range of sexually reproducing eukaryotes, including plants, fungi and animals (1). This process is initiated by the programmed formation of DNA double-strand breaks (DSBs), followed by their repair using the homologous chromosome as template. Recombination events can lead to the reciprocal exchange of flanking regions (crossovers, COs), or proceed without reciprocal exchange (non-crossovers, NCOs). COs are essential for the proper segregation of homologous chromosomes (2). Failure to form COs can lead to aneuploid reproductive cells or to defects in meiotic progression and sterility (3). Meiotic recombination also plays an important evolutionary role. It increases genetic diversity by creating novel allele combinations (4, 5) that in turns facilitate adaptation and the removal of deleterious mutations from natural populations (6–8).

Intriguingly, the CO rate varies not only among species, populations, sexes and individuals, but also along the genome (9–11). Broad-scale patterns of variation within chromosomes (megabase scale) have been observed in some species: low recombination rate near centromeres and high recombination rate in telomere-proximal regions (12). At a finer scale (kilobases), CO rate across the genome ranges from nearly uniform (*e.g.* flies, worms and honeybees) (13–15) to highly heterogeneous (*e.g.* yeast, plants and vertebrates). In such non-uniform recombination landscapes, most recombination events are typically concentrated within short intervals of about 2kb, called recombination hotspots (16, 17). Studies on the evolutionary dynamics of recombination hotspots have identified two alternative mechanisms for controlling hotspot localization. In many eukaryotes (*e.g. Arabidopsis*, budding yeast, swordtail fish, birds and canids), hotspots tend to be located near chromatin accessible regions enriched for H3K4me3, including promoters and transcription start sites (TSSs) (18–25). Elevated recombination rates are also observed at transcription end sites (TESs) in plants and birds (25, 26). In vertebrates, recombination hotspots are particularly associated with TSSs that are located within CpG islands (CGIs)(18, 25, 27). Hotspot location is conserved over large evolutionary timescales in birds and yeasts (22, 23, 25), likely because promoters are evolutionarily stable. However, the generality of this conclusion remains to be evaluated (28). On the other hand, mammalian species, including primates, mice and cattle, show a drastically different pattern. Their recombination hotspots tend to occur independently of open chromatin regions (29–32), and their positions evolve rapidly between closely related species and even populations (29, 30, 33-35). The genomic location of mammalian hotspots is controlled by the PRDM9 protein (32, 36, 37) that has four canonical domains (KRAB, SSXRD, PR/SET and zinc finger, ZF), among which the C2H2 ZF domain binds to a specific DNA motif. After PRDM9 binding to this motif, PRDM9 trimethylates H3K4 and H3K36 on adjacent nucleosomes through its SET domain. Then, the proteins required for DSB formation are recruited at PRDM9 binding sites. The formed DSBs are repaired by homologous recombination, leading to COs and NCOs (38). Two striking evolutionary properties of PRDM9 have been identified. First, PRDM9 triggers the erosion of its binding sites, due to biased gene conversion during DSB repair (32, 39, 40). Second, its ZF array presents a very high diversity (41–46) resulting from rapid evolution driven by a Red Queen dynamic in which positive selection favors the formation of new ZF arrays that recognize new binding motifs (39, 40, 47-50). This is the direct consequence of PRDM9 binding site erosion that increases the fraction of heterozygous binding sites where DSB repair efficiency is reduced, thus leading to lower fitness (47, 50-53). As a result, in *Mus musculus*, strains carrying different PRDM9 alleles generally share only 1-3% of DSB hotspots (34), and hotspot locations hardly overlap between humans and chimpanzees (29). Thus, PRDM9-dependent and -independent hotspots display different genomic locations and also evolutionary lifespan. This raises the question of why and how the genetically unstable mechanism of PRDM9-directed recombination has evolved (19, 54).

Understanding the function and evolutionary dynamics of PRDM9 in mammals has been a major breakthrough (29, 32, 36, 55). However, the function of PRDM9 homologs in non-mammal species remains poorly documented. Phylogenetic studies of the *Prdm9* gene have revealed the presence of a full-length copy in many metazoans, and also repeated partial or complete losses (19, 27, 56, 57). This is surprising for a gene that controls such a crucial mechanism. Among vertebrates, fine-scale recombination maps from species lacking *Prdm9* (*e.g.* birds and dogs) or harboring a truncated *Prdm9* (*e.g.* swordtail fish), revealed that their recombination hotspots are enriched at CGI-associated promoters (18, 19, 22, 25, 27), as observed in *Prdm9* knockout mice or rats (30, 58). This suggests that the partial or complete loss of *Prdm9* leads to a reversal of the default mechanism of hotspot location at gene promoters. Unfortunately, besides mammals, the currently available fine-scale recombination maps are essentially from species lacking a functional PRDM9 (*e.g.* fruit flies (13), birds (22, 25), three-spined stickleback (59), lizards (60) and honeybees (15)). This prevents any assessment of the link between PRDM9 presence and recombination landscape. Recent studies in snakes, which carry a full-length *Prdm9* copy, concluded that PRDM9 has a role in specifying recombination hotspots, and that a fraction of hotspots is at promoter-like features (61, 62). This complexity raises the question of PRDM9 function and how it evolved, particularly whether it was ancestrally involved in regulating recombination hotspots, or whether this function appeared relatively recently in mammals. To answer this question, we need to characterize the recombination landscapes in other non-mammalian taxa that harbor PRDM9 and determine whether their characteristics and dynamics are similar to those described in mammals.

To this aim, here, we investigated the putative function of PRDM9 in salmonids, a diverse family of teleost fishes in which a full-length *Prdm9* has been found (19, 56). The phylogenetic position of salmonids is ideal for testing the hypothesis of an ancestral PRDM9 role in regulating meiotic recombination in vertebrates. We used the large amount of genomic resources available in salmonids and also generated new data to test the role of PRDM9 in driving the location of recombination events in salmonids. Specifically, if PRDM9 role in salmonids were the same as in mammals, we would expect (i) the presence of recombination hotspots, (ii) located away from promoters, (iii) overlapping with enrichment for H3K4me3 and H3K36me3, (iv) showing rapidly evolving landscapes between closely related populations and species, and (v) associated with high diversity of the PRDM9 ZF domain. Importantly, salmonids have undergone two rounds of whole genome duplication (WGD) (63–65), offering the opportunity to investigate the impact of gene duplication on *Prdm9* evolutionary dynamics.

To test these hypotheses, we first analyzed the functional conservation of the many *Prdm9* duplicated copies across the phylogeny of salmonids. We then characterized the functional *Prdm9* allelic diversity in Atlantic salmon and rainbow trout to assess the evolutionary dynamics of the ZF array. We also determined the meiotic DSB landscape in rainbow trout using chromatin immunoprecipitation (ChIP) of the recombinase DMC1 followed by sequencing, and compared it with the genomic landscapes of the H3K4me3 and H3K4me36 marks. Lastly, we reconstructed linkage-disequilibrium (LD) based recombination landscapes in five populations from three different salmonid species to identify hotspots, test their association with genomic features, and measure their evolutionary stability. Our results provide a body of evidence supporting PRDM9 role as a determinant of recombination hotspots in salmonids.

## Results

### Duplication history and differential retention of *Prdm9* paralogs in salmonids

The analysis of the genomes of twelve salmonid species and of northern pike (used as outgroup) revealed multiple paralogous copies of the *Prdm9* gene. These paralogs partly resulted from two rounds of WGD: the teleost-specific WGD that occurred ∼320 Mya (referred to as Ts3R) (63, 65) and a more recent WGD in the common ancestor of salmonids at ∼90 Mya, after their speciation with pikes (referred to as Ss4R) (64). Taking advantage of the known pairs of ohnologous chromosomes resulting from WGD in salmonids (103–105), we reconstructed the duplication history of *Prdm9* paralogs by combining chromosome location information and phylogenetic inference. The number of *Prdm9* paralogs detected per genome ranged from 6 copies in rainbow trout, huchen and European grayling, to 14 in lake whitefish. Conversely, we found only 3 copies in northern pike. These paralogs clustered into two main groups that were previously identified as *Prdm9ɑ* and *Prdm9β* and originated from the Ts3R WGD (19). We found two additional subgroups among the *Prdm9β* copies (referred to as *β1* and *β2*) that were conserved in the twelve salmonid species, but only one *β* copy in the outgroup (S3 Fig). The *β* paralogs contained a complete SET domain and a conserved ZF domain, but all lacked the KRAB and SSXRD domains, as previously described (19). The *ɑ* sequences clustered into two well-supported groups of paralogs (named *ɑ1* and *ɑ2*) that could be subdivided in two groups of duplicated copies (designated as *ɑ1.1*/*ɑ1.2* and *ɑ2.1*/*ɑ2.2*; Fig 1A, S4 Fig). We found the sequence pairs *β1*/*β2*, *ɑ1.1*/*ɑ1.2* and *ɑ2.1*/*ɑ2.2* in three Ss4R ohnologous pairs, suggesting that they originated from the salmonid-specific WGD. We observed an additional subdivision within the *ɑ1* group, with pairs of copies duplicated in tandem present in each pair of ohnologs (*i.e. ɑ1.1 a* and *b* and *ɑ1.2 a* and *b*, Fig 1A, S7 Table). Although no phylogenetic signal was associated with the *a* and *b* copies, probably due to ectopic recombination and gene conversion, these copies are likely to represent a segmental duplication that preceded the Ss4R WGD. Thus, at least two *Prdm9* duplication events (*i.e.* one leading to *ɑ1/ɑ2* and the other to *ɑ1.a/ɑ1.b* copies) occurred in addition to the WGD-linked duplications. To summarize, our results indicate that *Prdm9 ɑ* and *β* copies originated from the Ts3R WGD. After the divergence of the Esociformes (pike) and Salmoniformes lineages ∼115 Myrs ago, the *ɑ* copy was duplicated on another chromosome, generating *ɑ1* and *ɑ2* copies. The *ɑ1* copy was subsequently duplicated in tandem, producing *ɑ1.a* and *ɑ1.b* copies on the same chromosome. Lastly, all these copies were duplicated on ohnologous chromosome pairs following the Ss4R WGD. This consensus evolutionary history was accompanied by gene conversion events and lineage-specific duplications and losses that were not fully identified in our analysis (Fig 1B). Most of these gene copies only contained a subset of the 10 expected exons and/or showed signatures of pseudogenization (stop codons, frameshifts). In each species, we identified exactly one complete *Prdm9* sequence with 9 to 11 exons, including the four canonical domains without evidence of pseudogenization The four genera *Coregonus*, *Thymallus*, *Oncorhynchus* and *Salvelinus* all shared the same *ɑ1.a.1* functional copy, while the complete copy found in the two *Salmo* species corresponded to the *ɑ1.a.2* paralog (Fig 1). Therefore, our results support the differential retention of functional *Prdm9ɑ* paralogs between salmonid lineages following the Ss4R WGD.

**Figure 1.**
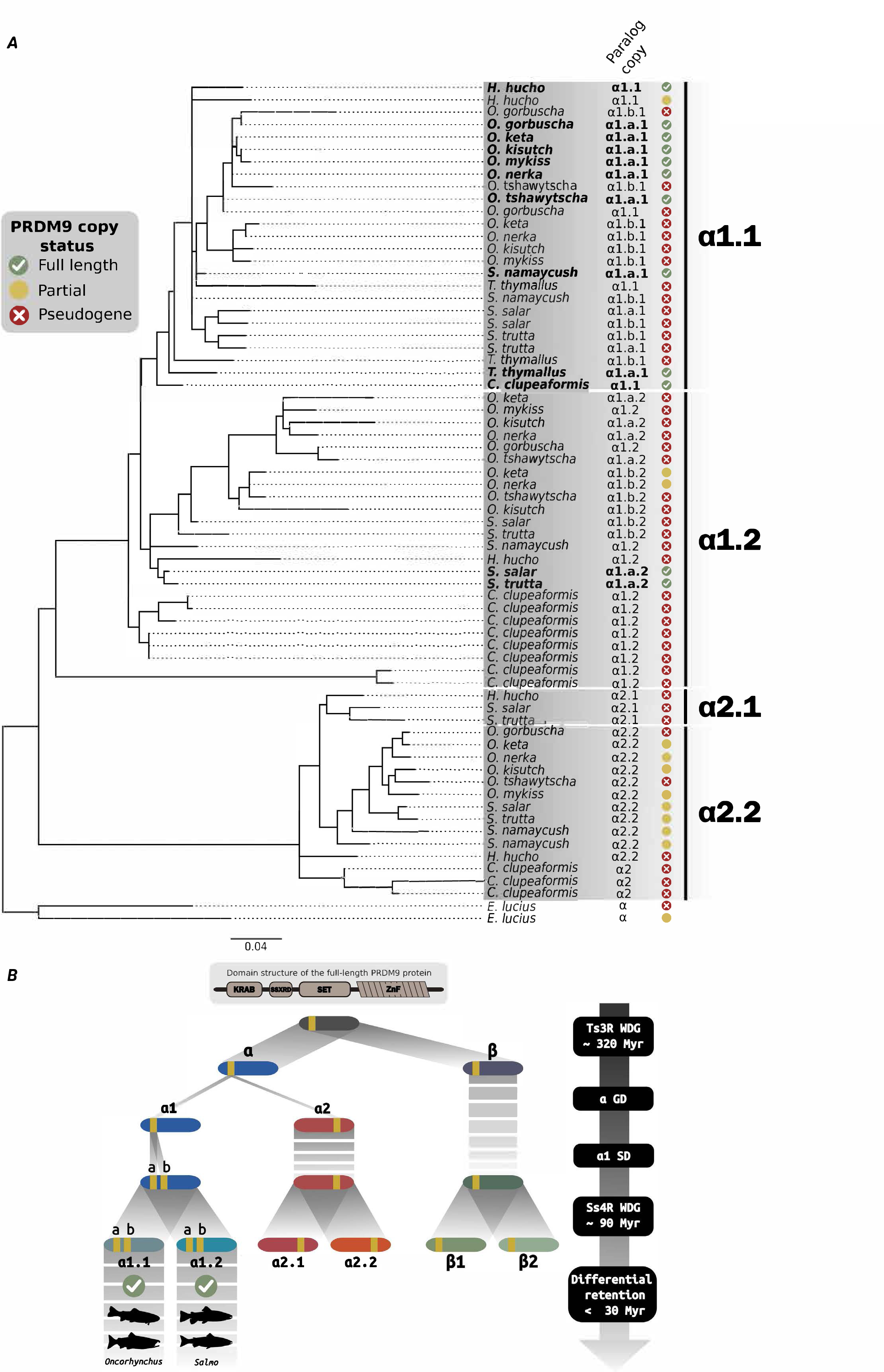
*Prdm9* duplication history in salmonids. **A)** Phylogenetic tree of *Prdm9α* paralogs in twelve salmonids (*i.e. Oncorhynchus kisutch*, *Oncorhynchus mykiss*, *Oncorhynchus nerka*, *Oncorhynchus keta*, *Oncorhynchus gorbuscha*, *Oncorhynchus tshawytscha*, *Salmo salar*, *Salmo trutta*, *Salvelinus namaycush*, *Hucho hucho*, *Coregonus clupeaformis*, *Thymallus thymallus*) and northern pike (*Esox lucius*) as outgroup species. *Prdm9*β is shown in S3 Fig. The phylogenetic tree was computed with IQTREE and the concatenated six exons of the three canonical PRDM9 domains KRAB, SSXRD and SET, with 1000 bootstrap replicates (values shown at nodes in S4 Fig). The columns, from left to right, indicate the i) species name; ii) annotated paralog copy (in bold: full-length copy without peusogenization), iii) *Prdm9* copy status. The four major α clusters. *Prdm9α* clusters into two main groups (*α1* and *α2*) that are divided in two subgroups (*α1.1/α1.2* and *α2.1/α2.2*). *α1* was duplicated in tandem before the salmonid-specific whole genome duplication Ts3R event (Ss4R), leading to the generation of a second copy: *α1.a* and *α1.b*. The scale bar is in unit of substitution per site. **B)** Consensus history of *Prdm9* duplication events in salmonids. After the teleost-specific whole-genome duplication (Ts3R WDG) ∼320 Mya, the chromosomes of the common ancestor of teleosts were duplicated. Two ohnolog chromosomes arose from the one carrying the ancestral *Prdm9* locus: one carrying the *Prdm9α* copy and the other the *Prdm9*β copy. Gene duplication (GD) of the *α* paralog (referred to as *α1*) led to the appearance of a new *α* copy (*α2*) on another chromosome. The *α1* copy (becoming *α1.a*) then underwent a segmental duplication (SD), generating a *α1.b* copy in tandem on the same chromosome. By this time, the β paralog had lost the KRAB and SSXRD domains. Lastly, the four copies were duplicated during the salmonids-specific Ss4R WGD ∼90 Mya, with the newly formed paralogs (annotated *α1.a.2*, *α1.b.2*, *α2.2*, β*2*) on ohnolog chromosomes. One full-length copy was retained in each species. The *Salmo* genus (*S. trutta* and *S. salar*) retained the *α1.2* copy, whereas all other salmonids retained the *α1.1* copy (only *O. kisutch* and *O. mykiss* are represented). This differential retention of paralogs must have happened after the speciation of the *Salmo* lineage ∼30 Mya. Ohnolog chromosomes are represented with similar color shades (*i.e.* blue, red, green), and *Prdm9* locus in yellow. This global picture of the duplication events in the salmonid history does not show other duplication events and losses that arose independently in some lineages.

### High PRDM9 ZF array diversity in *O. mykiss* and *S. salar*

We analyzed the allelic diversity of the ZF array of the complete PRDM9*α* copy found in *S. salar* (*α1.a.2*) and *O. mykiss* (*α1.a.1*) (Fig 1). We identified 11 PRDM9 ZF alleles in 26 *S. salar* individuals, and 7 alleles in 23 *O. mykiss* individuals (Fig 2A). The major allele had a frequency of 0.40 in *S. salar* and 0.35 in *O. mykiss*, and the four most frequent alleles had a cumulative frequency >0.8 in both species (Fig 2B). *S. salar* and *O. mykiss* alleles contained 5 to 10 and 7 to 15 ZFs, respectively. In both species, the last ZF of the arrays was probably not functional, because it lacked the conserved histidine involved in the interaction with a zinc ion required to stabilize the finger array (S5 Fig). As seen in other species, the four positions in contact with DNA (position −1, 2, 3 and 6 of the alpha helix) were highly variable among ZF units (Fig 2C). We characterized the proportion of total amino acid diversity at these DNA-binding residues among all different ZF units identified in each species following (19). This proportion, which is sensitive to the rapid evolution at DNA-binding sites and to the homogenization at other amino acid positions due to concerted evolution between repeats within the array, was 0.49 in *S. salar* and 0.55 in *O. mykiss* (Fig 2C). These values were within the range reported for full length PRDM9α in vertebrates (19). The observed high level of allelic diversity and the pattern of amino acid diversity within the ZFs were consistent with the rapid and concerted evolution of the ZF array of the full-length *Prdm9* gene that characterizes PRDM9 copies involved in specifying meiotic recombination sites (19, 54).

**Figure 2.**
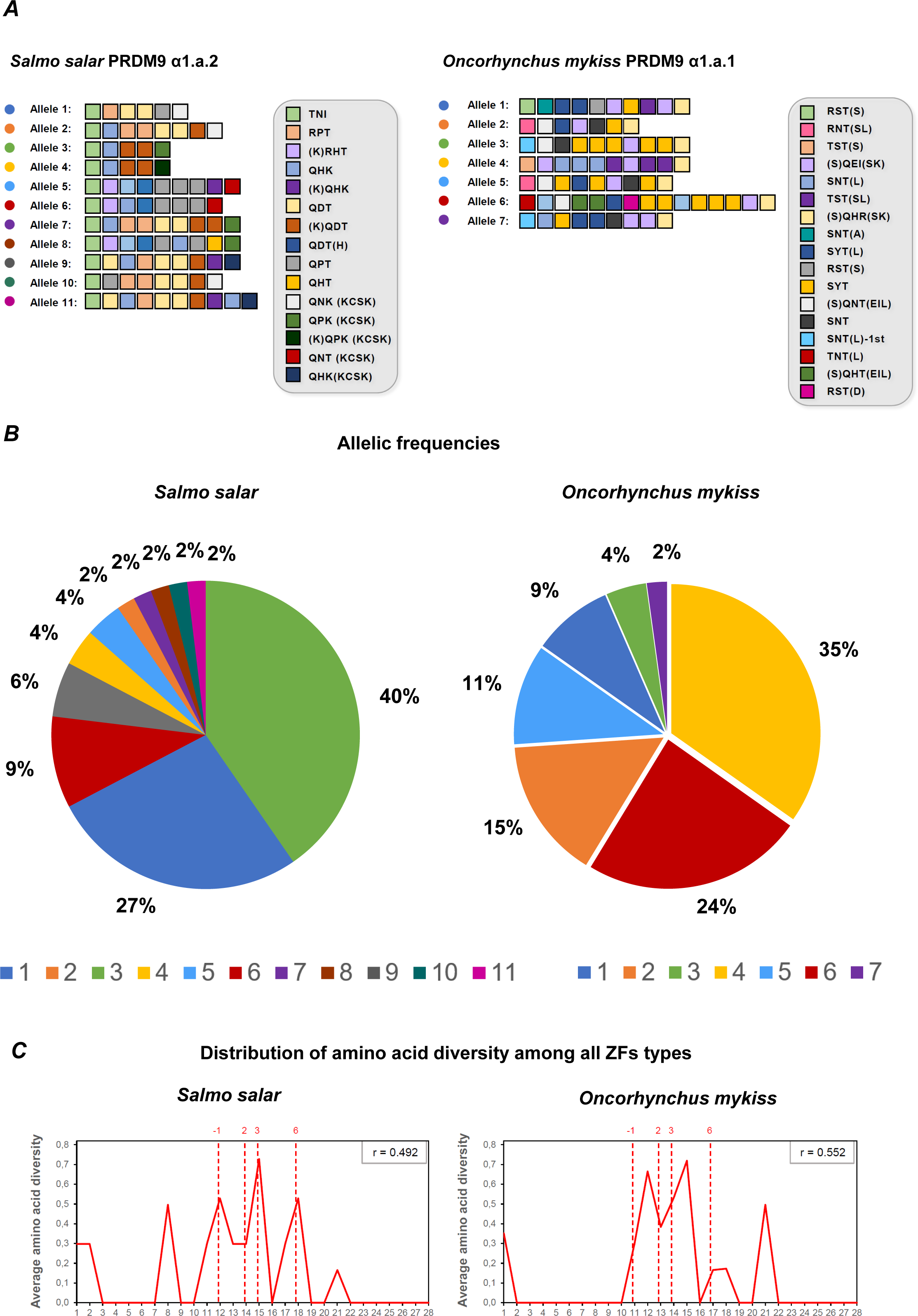
Zinc finger allelic diversity of full-length PRDM9 in *S. salar* and *O. mykiss*. **A)** Structure of the identified PRDM9 alleles in *S. salar* PRDM9 α1.a.2 and *O. mykiss* PRDM9 α1.a.1. Colored boxes represent unique ZFs, characterized by the three amino acids in contact with DNA (3-letter code). Additional variations relative to the reference sequence are indicated in between brackets. The complete zinc finger amino-acid sequences are shown in S5 Fig. **B)** Frequencies of the alleles displayed in panel A among the 26 *S. salar* and 23 *O. mykiss* individuals in which *Prdm9* was genotyped. **C)** Distribution of amino acid diversity among all unique ZFs found in the alleles shown in panel A, following a previously described methodology (19). The amino acid diversity is plotted in function of the amino acid position in the ZF array, from position 1 to position 28 (first and last residues) of a ZF. The ratio of amino acid diversity at the DNA-binding residues of the ZF array (−1, 2, 3 and 6), indicated as r, is shown in the upper box of each panel.

In addition to the full-length copy, one truncated copy of PRDM9*α* (*α2.2*) was strongly expressed in testes in both *Oncorhynchus* and *Salmo* genera. This truncated copy lacked a KRAB domain, but carried a SET domain that contained three conserved tyrosine residues important for methyltransferase catalytic activity (106) (S6 Fig). This suggested that its function, which remains unknown, involves the methyltransferase catalytic activity. In *S. salar*, the allelic diversity of the ZF array in the truncated *Prdm9 α2.2* was very low: in the 20 individuals analyzed, we only observed one single allele where the array had 5 ZF units (S5 and S7A Figs). In *O. mykiss*, we identified five PRDM9 *α2.2* alleles in 20 individuals, with 6 to 12 ZFs (S7A-B Fig). Some ZFs lost one amino acid, with unknown consequence on their DNA binding capacity (S5 Fig). In *O. mykiss*, the proportion of amino acid diversity at DNA-binding residues was lower in the truncated PRDM9 α2.2 (r=0.367, S7C Fig) than in the full-length PRDM9 α1.a.1 (r=0.552, Fig 2C). However, in *S. salar*, the proportion of amino acid diversity at DNA-binding residues was relatively high among the 5 ZFs of the unique PRDM9 α2.2 allele (r=0.471). PRDM9 α2.2 may have retained this signature of a past positive selection, suggesting that the functional shift associated with the loss of the KRAB domain may be relatively recent. Overall, in both species, the ZF diversity was strikingly reduced in the truncated PRDM9α (α2.2) compared with the full-length copy, consistent with the hypothesis that KRAB-less PRDM9 homologs lost the capacity to trigger recombination hotspots, and therefore are no longer subject to the Red Queen dynamics (19).

### PRDM9 specifies meiotic DSB hotspots in *O. mykiss*

To directly assess whether the full-length PRDM9α copy (hereafter PRDM9 unless otherwise specified) determines the localization of DSB hotspots, we investigated the genome-wide distribution of DMC1-bound ssDNA in *O. mykiss* testes by DMC1-SSDS (Fig 3A). DMC1 is a meiosis-specific recombinase that binds to ssDNA 3’ tails resulting from DSB resection. Therefore, meiotic DSB hotspots can be mapped by identifying fragment-enriched regions (*i.e.* peaks) in DMC1-SSDS data (30, 33, 75). We detected several hundred peaks in the three rainbow trout individuals analyzed by DMC1-SSDS (616 peaks in TAC-1, 209 in TAC-3, and 1924 in RT-52). Differences in peak number may result from inter-sample differences in cell composition related to the testis developmental stage (see S1 Methods). In all three individuals, the DMC1-SSDS signal at DSB hotspots displayed a characteristic asymmetric pattern in which forward and reverse strand reads were shifted toward the left and the right of the hotspot center, respectively. This confirmed that the DMC1-SSDS peaks detected in rainbow trout were genuine meiotic DSB hotspots (30)(S8A Fig). The average width of DMC1-SSDS peaks was 1.5-2.5 kb, which is similar to what described in mice and humans (30, 33). The DSB hotspot density increased towards the chromosome ends, indicating that the U-shaped distribution of COs classically observed in male salmonids (107) is the result, at least in part, of a mechanism controlling DSB formation (S9A Fig).

**Figure 3.**
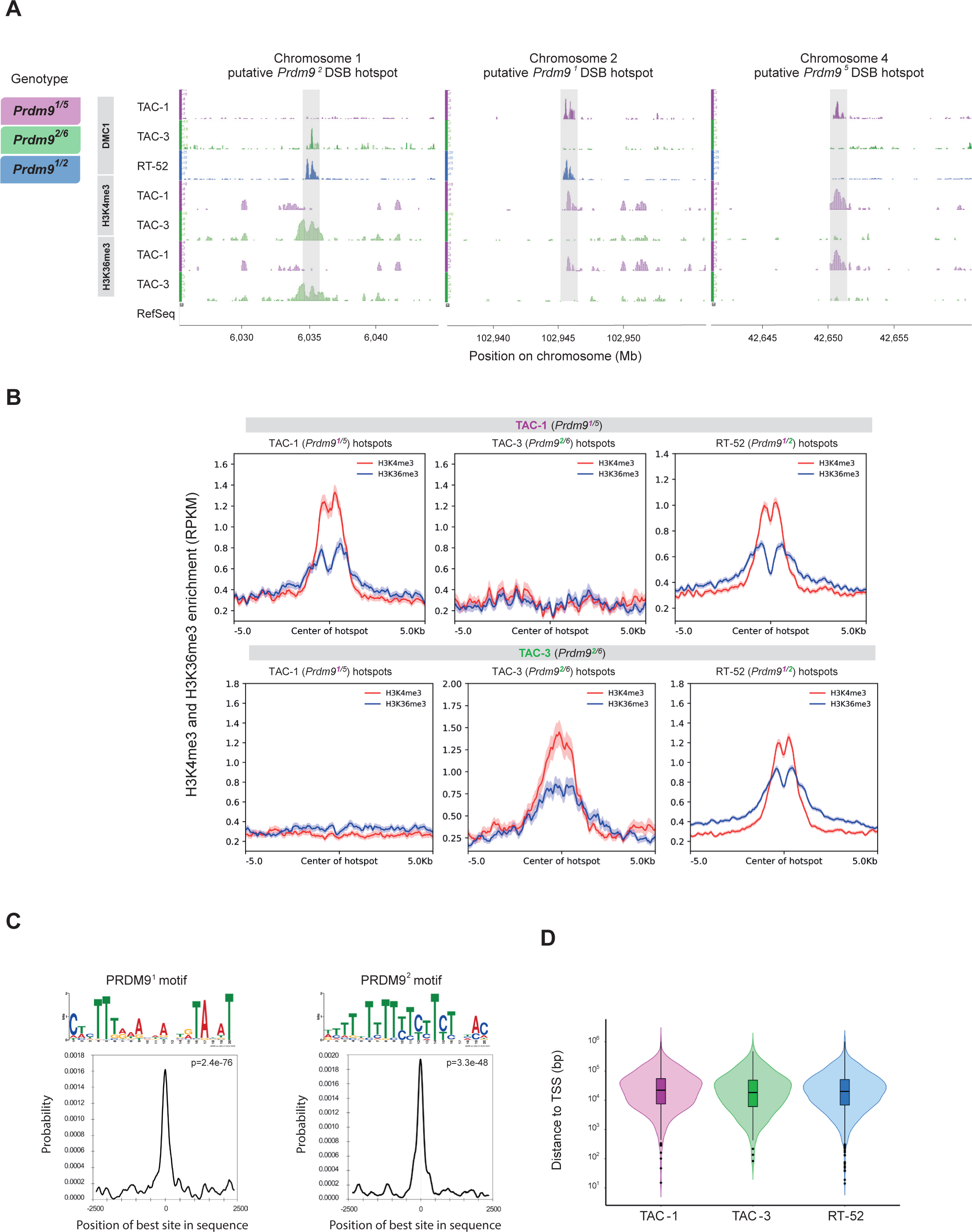
Meiotic DSB hotspots are specified by full length PRDM9 in *O. mykiss*. **A)** DSB hotspots detected by DMC1-SSDS (DMC1), H3K4me3 and H3K36me3 in selected regions of the *O. mykiss* genome in testes from two or three (DMC1) individuals. **B)** Average profile of H3K4me3 (red) and H3K36me3 (blue) ChIP-seq signal in TAC-1 (*Prdm9^1/5^*) and TAC-3 (*Prdm9^2/6^*) testes, at DSB hotspots detected in TAC-1 (*Prdm9^1/5^*), TAC-3 (*Prdm9^2/6^*) and RT-52 (*Prdm9^1/2^*). **C)** On top, the PRDM9 allele 1 (E=5.1e-37) and allele 2 motifs (E=1.2e-63) discovered at allele 1 (n=300) and allele 2 DSB sites (n=254) are shown. Below, the plots depict the distribution of hits for the PRDM9 allele 1 (left) and allele 2 (right) motifs at allele 1 and allele 2 DSB sites from the center of the sequence up to 2.5 kb of distance. The signal is the weighted smoothed moving average, and hits were calculated in a 250 bp sliding window. **D)** Violin plot showing the distribution of DSB hotspots in TAC-1 (magenta), TAC-3 (green) and RT-52 (blue) relative to the TSSs of RefSeq annotated genes.

Then, we tested whether the DSB hotspot formation was PRDM9-dependent by assessing the hotspot association with (*i*) specific *Prdm9* alleles and (*ii*) sites enriched for both H3K4me3 and H3K36me3 due to PRDM9 methyltransferase activity (38, 108). The three individuals analyzed (only TAC-1 and TAC-3 for histone marks) carried a functional *Prdm9* (*i.e. Prdm9 α1.a.1*) with different genotypes. TAC-1 (*Prdm9*^1/5^) and TAC-3 (*Prdm9*^2/6^) did not share any *Prdm9* allele, whereas RT-52 (*Prdm9*^1/2^) shared one allele with each of them. In line with the hypothesis that PRDM9 specifies DSB hotspots, some DMC1-SSDS peaks were common to RT-52 and either TAC-1 or TAC-3 (see Fig 3A for examples). Specifically, the overlap between TAC-1 and RT-52 DSB hotspots (167 of the 616 TAC-1 hotspots, 27%), and between TAC-3 and RT-52 DSB hotspots (42 of the 209 TAC-1 hotspots, 20%) was substantial, whereas only two hotspots were shared by all three individuals (S8B Fig). The 55 DMC1-SSDS peaks shared by TAC-1 and TAC-3 may be artifactual because the forward and reverse strand enrichment distribution did not follow the typical asymmetric pattern of DSB hotspots (compare S9B Fig and S8A Fig). The histone marks H3K4me3 and H3K36me3 usually do not colocalize at the same loci because H3K4me3 is enriched at promoters and other genomic functional elements and H3K36me3 within gene bodies. However, PRDM9-dependent DSB hotspots are specifically enriched for both H3K4me3 and H3K36me3 because PRDM9 catalyzes both modifications (109, 110). As expected, we did not detect H3K36me3 enrichment at PRDM9-independent H3K4me3 peaks identified in brain tissue (S10A Fig). Conversely, the DSB hotspots detected in TAC-1 and TAC-3 testes were enriched for both H3K4me3 and H3K36me3 (Fig 3A-B, S10B-C Fig). However, in testis chromatin from TAC-1 but not from TAC-3 H3K4me3 and H3K36me3 were enriched at TAC-1 DSB hotspots and reciprocally, they were enriched at TAC-3 DSB hotspots in TAC-3 but not in TAC-1. H3K4me3 and H3K36me3 were enriched at RT-52 DSB hotspots in testis chromatin from both TAC-1 and in TAC-3 (S10B-C Fig). H3K4me3 and H3K36me3 enrichment levels were correlated, as expected for histone marks catalyzed by PRDM9 (S11A Fig). Moreover, the majority of RT-52 DSB hotspots were enriched for H3K4me3 either in testis chromatin from TAC-1 or in TAC-3, but not in both (S11B Fig; Chi2 test of homogeneity, p=10^-26^). A similar trend for the H3K36me3 mark might have been undetectable due to the high level of PRDM9-independent H3K36me3 at a fraction of the sites (S11A-B Fig). This observation supports the hypothesis that a fraction of DSB hotspots in RT-52 is specified by PRDM9^1^ (shared with TAC-1) and the others by PRDM9^2^ (shared with TAC-3).

### Population genomic landscapes of recombination

The DMC1-SSDS approach we performed allowed analyzing DSB distribution in a given male individual, is thus restricted to one sex and does not provide information on the outcome of recombination events (CO or NCO). To get a more general picture of the genome-wide recombination landscapes and their evolution, we computed LD-based genetic maps in three salmonid species: coho salmon (*O. kisutch*), rainbow trout (*O. mykiss*), and Atlantic salmon (*S. salar*). In *S. salar*, we analyzed three populations: North Sea (NS), Barents Sea (BS) and Gaspesie Peninsula (GP). For comparison, we also reconstructed the LD-based recombination map of European sea bass (*D. labrax*) that carries the KRAB-less *Prdm9β* gene, but lacks a full-length *Prdm9ɑ*.

The population-scaled recombination landscapes showed consistent broad-scale characteristics between *O. kisutch*, *O. mykiss* and the three *S. salar* populations. The genome-wide population recombination rate ranged from 0.0032 (in units of *⍴*=4*N*_e_*r* per bp) in *O. kisutch* to 0.012 in *O. mykiss*, with intermediate values in *S. salar* populations (Table 1). At the intra-chromosomal level, 100-kb smoothed recombination landscapes showed a general increase towards the chromosome ends, up to a 6-fold increase in *S. salar* (S12 Fig). This U-shape pattern mirrored the chromosomal distribution of DSB hotspots in male rainbow trout (S9 Fig). The fine-scale analysis of the genomic landscapes also showed highly heterogeneous recombination rates within 2-kb windows (Table 1, S13 Fig). In each population, the local variation in recombination rate was of several orders of magnitude (S8 Table). On average, 90% of the total recombination appeared to be concentrated in 20% of the genome, a higher rate than what observed in humans and chimpanzees (29, 111) and slightly higher than what we observed in sea bass (Table 1, S14 Fig, S8 Table). This heterogeneity was largely driven by the presence of recombination hotspots. Based on the raw LD-maps reconstructed at each SNP interval, we confirmed that the size of most (>80% on average) salmonid hotspots was <2kb (S15 Fig, S8 Table). Therefore, we performed the rest of our analysis using the hotspots called within 2-kb windows. The total number of called hotspots per species ranged between 17,064 in *S. salar* and 22,948 in *O. kisutch*, with hotspot density values similar to those in sea bass and also humans, mice and snakes (62, 111, 112). The proportion of total recombination cumulated in hotspots ranged between 17% in *S. salar* and 36% in *O. kisutch*, while occupied less than 3% of the genome (Table 1). Hotspots were evenly distributed along chromosomes, but showed higher recombination rates towards the chromosome ends. This property was observed both for species with a full-length *Prdm9* (*O. kisutch*, *O. mykiss* and *S. salar*) and for the species with a truncated *Prdm9* (*D. labrax*)(S16 Fig).

**Table 1.** Summary of fine-scale recombination rate variations in 2-kb windows and hotspot detection for populations of *O. kisutch*, *O. mykiss* and *S. salar* (only the NS population is shown), and *D. labrax*.

|  | *O. kisutch* | *O. mykiss* | *S. salar* | *D. labrax* |
| --- | --- | --- | --- | --- |
| Genome wide recombination rate (rho/bp) | 0.0032 | 0.0012 | 0.0085 | 0.039 |
| Cumulative amount of recombination in the 20% most recombining regions | 90.1% | 89.1% | 98.1% | 84.6% |
| Number of hotspots | 22948 | 21145 | 17064 | 7897 |
| Fraction of recombination in hotspots | 36.7% | 19.3% | 18.3% | 26.5% |
| Fraction of the genome occupied by hotspots | 2.7% | 2.1% | 1.3% | 1.9% |
| Hotspot density (per Mb) | 13.6 | 10.8 | 6.8 | 9.6 |

Then, we compared the LD-based recombination landscape of *O. mykiss* and the location of DSB hotspots mapped by DMC1-SSDS (pooling peaks from the three samples). We found that 6.7% of DMC1-SSDS peaks overlapped with the LD-based hotspots, which is more than expected by chance (S17A-B Fig). This weak overlap was comparable with that observed in *Mus musculus castaneus* where 12% of DSB hotspots overlap with LD-based hotspots (113). We also found that in these shared peaks, population recombination rates were significantly higher than in non-shared LD-based or DSB hotspots and the rest of the background landscape (Kruskal-Wallis test p-value < 0.05, Wilcoxon post-hoc test < 0.05, S17C Fig).

### Recombination hotspots are located away from CGIs

In species that lack full-length PRDM9, recombination hotspots should be located in open-chromatin regions, such as unmethylated CGI-associated promoters and/or constitutive H3K4me3 sites (18, 19, 22, 25, 27), unlike in species like mice, where PRDM9 targets regions away from these genetic elements (30). To test whether PRDM9ɑ plays a similar role in salmonids, we first examined how DSB hotspots were distributed relative to TSSs in rainbow trout. We found that the percentage of DSB hotspots overlapping with TSSs was either not different or lower than expected by chance (4.5 and 5.3% versus 7.6% for TSSs of coding and non-coding genes; S9 Table, S8C Fig). Moreover, the vast majority of DSB hotspots mapped several kb or more away from the closest TSS (Fig 3D). Therefore, DSB hotspots, at least those strong enough to be detected by our DMC1-SSDS assay, did not localize at TSSs.

We then examined how population recombination rates were distributed relative to TSSs that overlapped or not with CGIs, by comparing the three salmonid species to sea bass that only has a truncated PRDM9β protein. Although the criteria classically used to predict CGIs in mammals and birds are not appropriate for teleost fish where CGIs are CpG-rich but have a low GC-content (94, 95), we could predict TSS-associated CGIs in fish genomes simply based on their CpG content (see S2 Analysis). Sea bass (truncated PRDM9β) showed a high level of recombination at promoter regions, with a strong 3-fold enrichment of recombination at TSSs associated with CGIs (Fig 4A), as reported in birds (25). Conversely, in salmonid species (full-length PRDM9), recombination rate varied little between TSSs and their flanking regions (at most 1.2-fold enrichment). Specifically, at CGI-associated TSSs, recombination rate tended to be lower than at other TSSs (Fig 4A-B). Moreover, hotspots overlapping with TSS represented <5% of all hotspots in the three salmonid populations and up to 21% in sea bass (Fig 4C). The analysis of other genomic features showed little variation in recombination rate and hotspot density, with similar levels in genes, introns, exons, TEs and CGIs compared with intergenic regions (Fig 4B-C). We observed only a very small increase in recombination rate at TSSs that did not overlap with CGIs and TESs in *O. kisutch* and *O. mykiss*. Therefore, our results indicated that salmonid recombination events do not concentrate at promoter-like features overlapping with CGIs, as already shown in primates and in the mouse.

**Figure 4.**
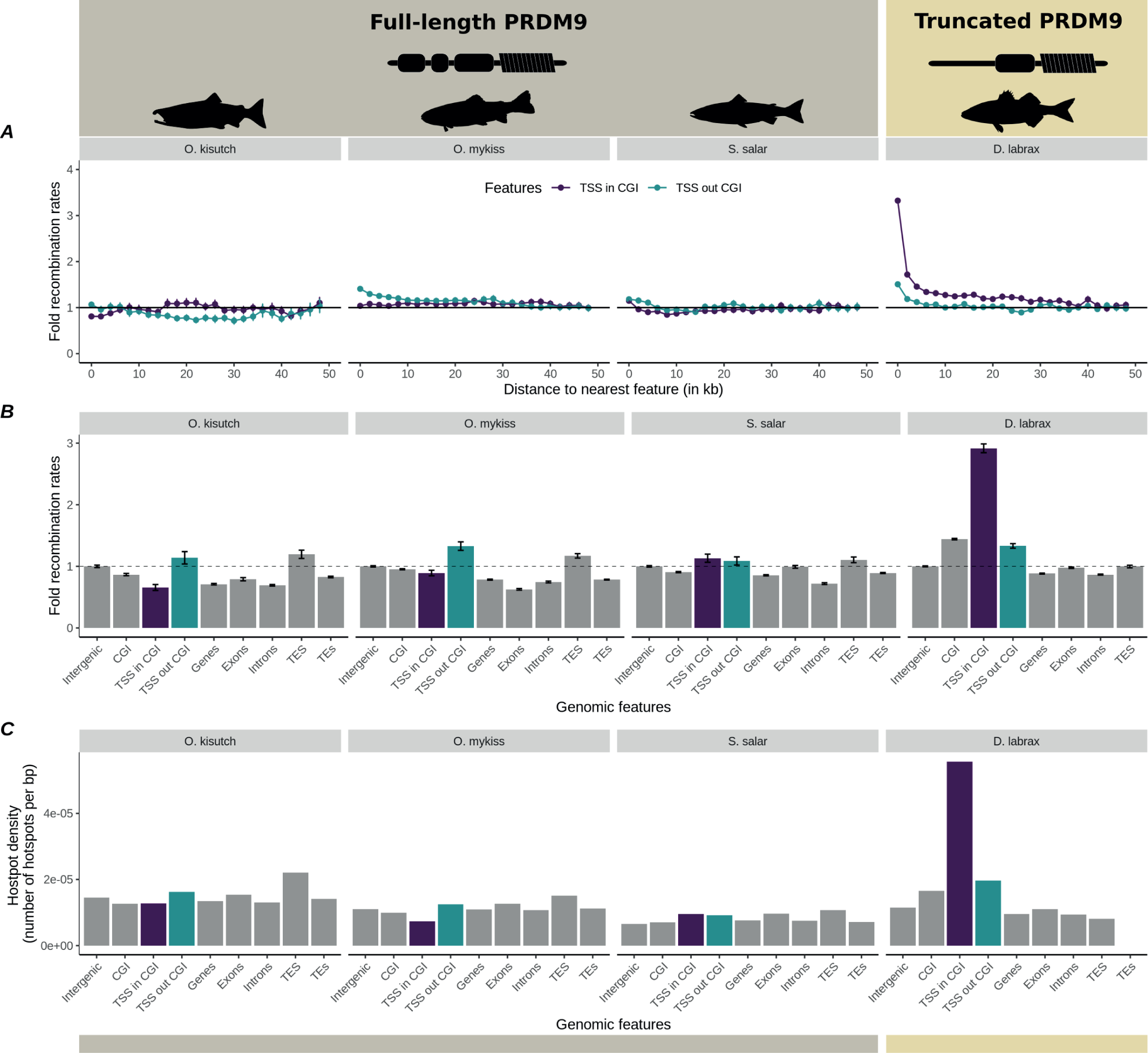
Recombination rates at genomic features. The recombination rates at different genomic features are shown for *O. kisutch*, *O. mykiss* and *S. salar* (NS population), and compared to those of sea bass (*Dicentrarchus labrax*) that lacks a full-length PRDM9 copy. **A)** Fold recombination rates (scaled to the average recombination rate at 50kb from the nearest feature) according to the distance to the nearest TSS (overlapping or not with a CGI). **B)** Fold recombination rates (scaled to the average recombination rates in intergenic regions) at the indicated genomic features. **C)** Hotspot density at the indicated genomic features. TSS and TES were defined as the first and last positions of each gene. CGIs were mapped with EMBOSS using CpGoe > 0.6 and GC > 0.

We also examined other genomic correlates and features that might influence population recombination rate variation at different levels of resolution. As expected from the joint effect of the local effective population size (*N*_e_) on both nucleotide diversity and population recombination rate, SNP density was positively correlated with the *⍴* averaged at the 100-kb scale, although this trend was not significant in *O. mykiss* (S18A Fig). More locally, we also observed an increase in SNP density in the 10kb surrounding recombination hotspots (S19A Fig). These positive relationships could be amplified by a direct mutagenic effect of recombination during DSB repair, and a more pronounced erosion of neutral diversity in low-recombining regions due to linked selection (29, 33, 114-116). However, we cannot exclude the possibility that the accuracy of the recombination rate estimate depends on SNP density (79), leading to possible confounding effects.

In mammals, GC-biased gene conversion causes an increase in GC-content at recombination hotspots (33, 40, 117, 118). Conversely, in four of the five salmonid populations, GC content tended to decrease close to hotspots (S19B Fig). At a larger scale (*i.e.* 100 kb), we observed significant positive correlations between GC content and recombination rates (S18B Fig). However, these correlations were very weak, suggesting that GC-biased gene conversion has a very small impact in salmonids compared with mammals and birds (118).

The salmonid genomes contain a large proportion (∼50%) of TEs among which Tc1/mariner is the most abundant superfamily (>10%)(103). It is not known whether Tc1/mariner transposons influence the estimation of recombination rates. Our TE analysis identified between 47.37% and 52.26% of interspersed repeats in *O. kisutch* and *S. salar*, respectively, and showed that 12.48% to 14.7% of the genome was occupied by Tc1-mariner elements (S6 Table). TEs and intergenic regions showed similar average recombination rates and hotspot density (Fig 4B-C). Recombination rates tended to slightly increase with TE density at the larger scale, except in *O. mykiss* for which we observed the opposite relation (S18C Fig), without any strong effect of the TE superfamilies (S20 Fig). As recombination rates and hotspot density at TEs were globally comparable to those at intergenic regions (Fig 4B-C), TEs and among them Tc1-mariner elements did not seem to be characterized by extreme recombination values that may have affected our recombination rate estimations.

Lastly, residual tetrasomy resulting from the salmonid WGD event at ∼90 Mya (Ss4R) (64) is observed at several chromosome regions characterized by increased genomic similarity between ohnologs. This could also affect the inference of LD-based recombination rates. Such regions have been identified in *O. kisutch*, *O. mykiss* and *S. salar* (103–105). We tried to filter non-diploid allelic variation from chromosomes showing residual tetrasomy, and we also controlled their effect by comparing their recombination patterns with those of fully re-diploidized chromosomes. Overall, we found <2-fold increase of the mean recombination rate in chromosomes containing tetraploid regions (S21A Fig). This was mostly explained by the local increase towards the end of chromosomes with residual tetraploidy compared with fully re-diploidized chromosomes, an effect that was especially pronounced in *O. mykiss* (S21B Fig). Nevertheless, recombination rates behaved similarly in function of the distance to the nearest promoter-like feature in the two chromosome sets, and rate variations were similar between genomic features (S22 Fig). Overall, chromosomes containing regions with residual tetraploidy and re-diploidized chromosomes showed similar recombination patterns.

### Rapid evolution of recombination landscapes

Another key feature of the mammalian system is the rapid evolution of PRDM9-directed recombination landscapes due to self-induced erosion of its binding DNA motif and rapid PRDM9 ZF evolution (29, 32, 34). To determine whether this feature was present also in salmonids, we compared the location of recombination hotspots in the two *Oncorhynchus* species and in two geographical lineages and two closely related populations of *S. salar*. We estimated that only 6.2% of hotspots were shared by *O. kisutch* and *O. mykiss*, which diverged from their common ancestor about 16 Myr ago (119). Although this value was significantly higher than expected by chance (S23A Fig), there was almost no increase in recombination rate at the orthologous positions of hotspots in the two species (Fig 5A). Similarly, the two genetically differentiated lineages of *S. salar* only shared 10.3% (GP vs. BS, *F*_ST_ = 0.26) and 11.2% (GP vs. NS, *F*_ST_ = 0.28) of their hotspots, with a weak recombination rate increase at the alternate lineage hotspots (Fig 5B, S23B-C Fig). Conversely, the two closely related BS and NS *S. salar* populations (*F*_ST_ = 0.02) shared 26.3% of their hotspots, which was much more than expected by chance (Fig 5C, S23D Fig). In addition, recombination rate at NS hotspots in the BS population showed a 5-fold increase (and reciprocally), reflecting high correlation between BS and NS recombination landscapes (Spearman’s rank coefficient > 0.7, p-value < 0.05; S24 Fig). Overall, these analyses revealed a rapid evolution of hotspot localization between species and also between geographical lineages of the same species. Only closely related populations shared a substantial fraction of their hotspots, possibly because they share similar sets of *Prdm9* alleles that recognize common binding DNA motifs.

**Figure 5.**
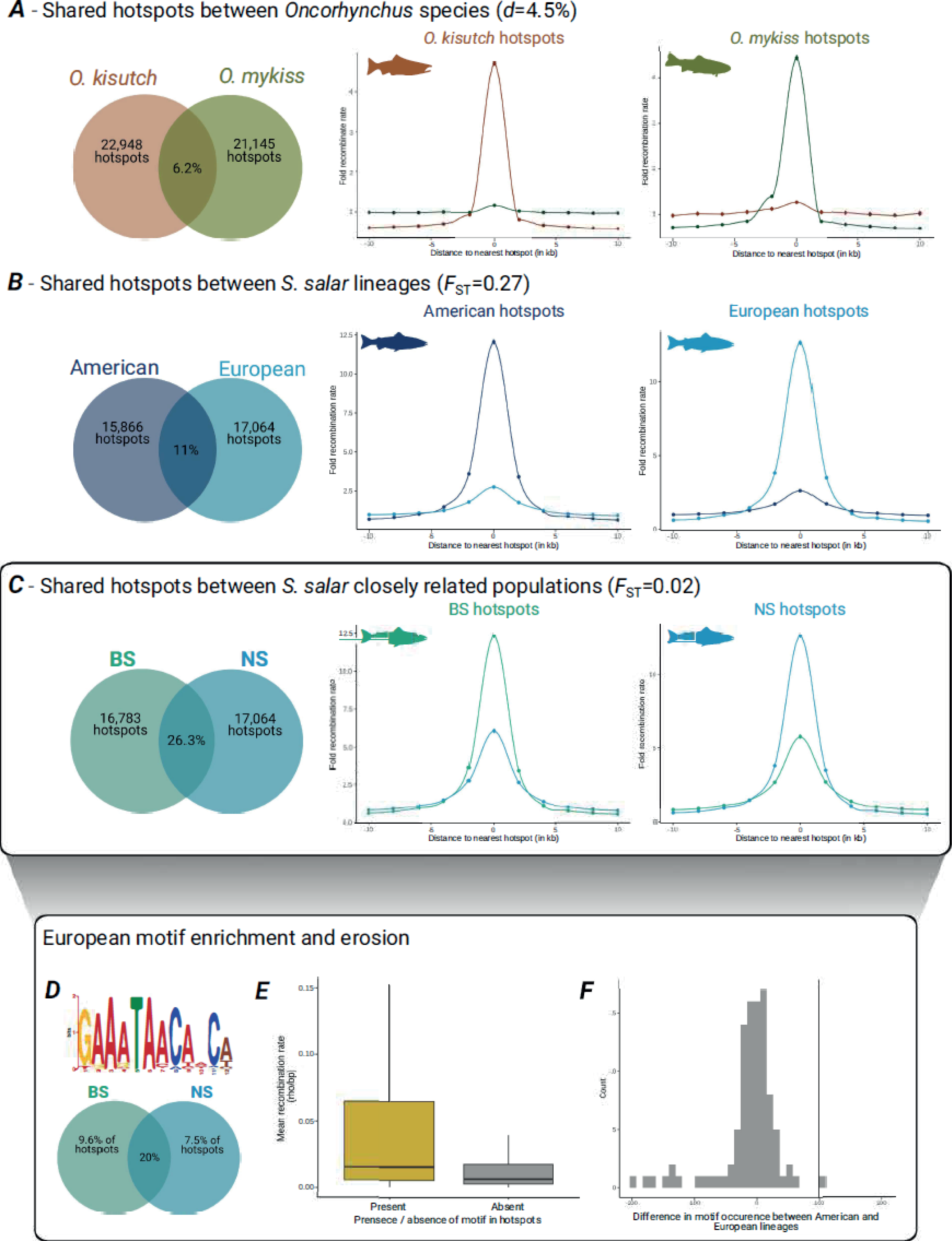
Recombination hotspots shared between populations and motif enrichment. In panels A, B and C, the Venn diagrams (left) show the percentages of recombination hotspots shared between pairs of taxa, and the graphs (middle and right) show the recombination rates around hotspots at orthologous loci in the two taxa, for the two *Oncorhynchus* species (**A**), the American (GP population) and European (BS and NS populations) *S. salar* lineages (**B**), and between the two closely related European *S. salar* populations (BS and NS) (**C**). The percentage of shared hotspots was calculated using the number of hotspots in the population with fewer hotspots as the denominator. **D)** Motif found enriched in the hotspots identified in the European populations of *S. salar* (BS and NS). The Venn diagram shows the percentages of population-specific and shared hotspots where the motif was found. **E)** Mean recombination rate at shared (between the BS and NS populations) hotspots that harbor (936 hotspots) or not (3485 hotspots) the detected motif. The recombination rate was significantly higher at hotspots with the motif (Student’s *t-*test p-value < 0.05). **F)** Motif erosion in the European *S. salar* populations. The vertical line represents the observed difference in the occurrence of the motif in panel D between the American and European lineages. The null distribution (in gray) shows the difference for 100 random permutations of the motif.

### Motifs enriched at hotspots show signs of erosion

A landmark of the PRDM9-dependent hotspots identified in mammals is the presence of DNA motifs, as a consequence of the sequence-specificity of the PRDM9 ZF domain. Therefore, we investigated the presence of PRDM9 allele-specific DNA motifs enriched at hotspots in salmonids. We first searched for potential PRDM9 binding motifs in rainbow trout, focusing on RT-52 (*Prdm9^1/2^*) DSB-based hotspots. As the *Prdm9^1^*allele is present in RT-52 and TAC-1 (*Prdm9^1/5^*), we defined a subset of RT-52 DSB hotspots presumably specified by PRDM9^1^, based on their overlap with H3K4me3/H3K36me3 peaks in TAC-1 (n=300). Similarly, we defined a subset of DSB hotspots enriched in putative targets of the PRDM9^2^ allele, which is present also in TAC-3 (*Prdm9^2/6^*) (n=254). We identified two consensus motifs: one strongly enriched in PRDM9^1^ DSB hotspots and the other in PRDM9^2^ DSB hotspots (Fig 3C). Consistent with the *Prdm9* genotypes of the three rainbow trout samples, both motifs were enriched at RT-52 DSB hotspots. The PRDM9^1^ motif was also enriched in TAC-1 DSB hotspots and the PRDM9^2^ motif in TAC-3 DSB hotspots (S25 Fig). Moreover, the PRDM9^1^ motif was co-centered with DSB hotspots only in RT-52 (Fisher’s test, p=8.5×10^−196^) and TAC-1 (p=7.7×10^−27^), while the PRDM9^2^ motif was co-centered with DSB hotspots in RT-52 (p=3.3×10^−97^) and TAC-3 (p=1.7×10^−5^) (S26 Fig). These two consensus motifs were also significantly enriched at LD-based hotspots (S25 Fig). Particularly, the motif targeted by PRDM9^1^ was strongly enriched at the center of LD-based hotspots (S26 Fig), suggesting that this allele (or closely related alleles that recognize similar DNA sequences) has been quite frequent during the recent history of the wild population under study.

As PRDM9-binding DNA motifs are allele-specific, the sharing of *Prdm9* alleles between populations should lead to shared motif enrichment at shared hotspots. Therefore, we looked for enrichment of potential 10-20-bp motifs in the population-specific and shared hotspots of the three *S. salar* populations, compared with a control set of random sequences of equal size and GC-content. Of note, as LD-based hotspots reflect the population-scaled recombination rate, they may result from the activity of multiple PRDM9 variants that can hinder the discovery of targeted motifs. Nevertheless, after filtering candidate motifs (S27A Fig), we found a motif that was enriched in the NS and BS populations: 7.5% and 9.6% of their hotspots, respectively, and 20% of their shared hotspots (Fig 5D). Overall, the recombination rates at hotspots overlapping with this 12-bp motif were significantly higher than those at other hotspots (Student’s *t*-test p-value <0.05; Fig 5E, S27B-C Fig). This suggests that the detected motif is targeted by a frequent PRDM9 variant shared by the two closely related NS and BS populations, possibly originating from their common ancestral variation.

PRDM9-associated hotspot motifs undergo erosion in mammals due to biased gene conversion (32, 39, 40). Therefore, we tested whether the identified 12-bp motif showed signs of erosion in European *S. salar* populations. By comparing the number of motifs present in the available long-read genome assemblies from seven European and five North American Atlantic salmon genomes (accession numbers in S1 Methods), we found a 2.97% reduction in the mean number of motifs in the European genomes (mean Europe = 3,230 vs. mean North America = 3,329). This level of erosion was significant and not explained by differences in assembly sizes, as revealed by count comparisons on collinear blocks, obtained following 100 random permutations of the motif (Figure 5F). Therefore, the enriched motif shared by the NS and BS populations was partially eroded in the European lineage, as predicted by the Red Queen model of PRDM9 evolution.

## Discussion

To determine whether the PRDM9 functions characterized in humans and mice are shared by other animal clades or whether they correspond to derived traits, we investigated the evolution and function of full-length *Prdm9* in salmonids using phylogenetic, molecular and population genomic approaches. These analyses allowed us to determine the evolutionary history of *Prdm9* gene duplication and loss, the diversity of the PRDM9 ZF array, the historical sex-averaged recombination map in several populations, the locations of meiotic DSB sites in spermatocytes, their chromatin environment, and the presence of conserved motifs and their erosion. Collectively, these analyses led us to conclude that PRDM9 triggers recombination hotspot activity in salmonids through a mechanism similar to that described in mammals.

### PRDM9 specifies recombination sites in salmonids

Our conclusion is based on several evidences. First, we showed in *O. mykiss* that DSB hotspots, detected by DMC1-SSDS, are enriched for both H3K4me3 and H3K36me3 marks. We provide evidence that hotspot localization is determined by PRDM9 ZFs because the location of DSB hotspots and the associated H3K4me3 and H3K36me3 marks varied in function of the *Prdm9* ZF alleles present in the tested individuals (Fig 3A-B). Consistent with this interpretation, we identified DNA motifs enriched at DSB sites. Thus, in salmonids, PRDM9 retained its DNA binding activity to modify histones and to attract the recombination machinery at its binding sites. Comparison of DSB hotspots detected by DMC1-SSDS with LD-based CO hotspots in *O. mykiss* showed a limited, but significant overlap (S17 Fig). We also identified a DNA motif enriched at DSB hotspots targeted by *Prdm9^1^* that was also enriched at the center of strong CO hotspots detected in the LD-based recombination map (S26 Fig). The overlap between hotspots is compatible with the presence of a common *Prdm9* allele(s) between the individuals tested and the prevalent *Prdm9* allele(s) during the history of the populations analyzed. However, as the population-scaled recombination landscape in *O. mykiss* has been shaped by a diversity of alleles, not necessarily represented in the three studied individuals, the overall hotspot overlap was low. Similar recombination landscapes driven by multiple PRDM9 alleles have been described in mouse populations (35, 113). In the mouse, PRDM9 can suppress the recombination activity at chromatin accessible regions (30). Here, we found that this function is conserved in salmonids. Both DSB maps and LD maps show that recombination events are not enriched in promoter regions in salmonids (Fig 3D and 4). Conversely, in *D. labrax*, recombination hotspots were strongly enriched at promoter regions overlapping with CGIs (Fig 4), as reported for other vertebrate species that do not have a full-length *Prdm9* (18, 19, 22, 25).

In addition to the localization of recombination shaped by PRDM9, we detected a higher recombination activity at telomere-proximal regions when measuring DSB activity and LD, consistent with the recombination activity measured in *S. salar* pedigree-based linkage maps (107). We infer that this effect is PRDM9-independent because the putative PRDM9 motifs in *O. mykiss* (derived from DMC1-SSDS) and *S. salar* (derived from LD-based hotspots) did not show such biased distribution (S28 Fig). We hypothesize that some additional factor(s) might modulate PRDM9 binding or any other step required for DSB activity along chromosomes. This telomere-proximal effect appears to be a conserved property, but of variable strength between sexes and among species with/without *Prdm9* (120).

### PRDM9 evolutionary instability

Similarly to the pattern reported in mammals (32, 57, 121), we found an outstanding diversity of PRDM9 ZF alleles in *O. mykiss* and *S. salar* and signatures of positive selection for ZF residues that interact with DNA, specifically in the full-length PRDM9 paralog (α1.a.1 and α1.a.2, respectively) (Fig 2). This suggests that full-length PRDM9 in salmonids could be involved in a Red Queen-like process, as documented in mammals, whereby the ZF sequence responds to a selective pressure arising from the erosion of PRDM9 binding motifs (39, 40, 47-50). Consistent with this hypothesis, we found almost no overlap of LD hotspots in the two *Oncorhynchus* species we compared. A similar comparison performed in three *S. salar* populations revealed that the percentage of shared hotspots decreased with the increasing genetic divergence (Fig 5). The 26.3% overlap in hotspot activity we detected in the two Norwegian populations could reflect the existence of shared *Prdm9* alleles. On the other hand, the European and Northern American salmon populations, which belong to two divergent lineages, may not share the same *Prdm9* alleles and as a possible consequence, only have 10.5% of common hotspots. Such patterns of population-specific hotspots and partial overlaps have been observed also in mouse populations (35), great apes (122) and humans (123). However, hotspot overlapping is always well below the 73% of shared hotspots between zebra finch and long-tailed finch that do not carry *Prdm9* (25). Further support for a *Prdm9* intra-genomic Red Queen process in Atlantic salmon came from the detection of an enriched motif in 20% of the hotspots shared by the NS and BS populations. As this motif is likely to be the target of an active *Prdm9* allele in European populations, the average 3% decrease in total copy number in European populations compared with North American populations is indicative of ongoing motif erosion.

Another intriguing pattern revealed by our study is the complex duplication history of the *Prdm9* gene in salmonids, shaped by WGD events and by segmental duplications. Particularly, the differential retention of paralogs between salmonid genera suggests that two functional *Prdm9ɑ* copies have co-existed in the common ancestor to *Salmo* and the (*Coregonus*, *Thymallus*, *Oncorhynchus*, *Salvelinus*) group (Fig 1). This might be true also in primates where the pair of paralogs formed by *Prdm7* and *Prdm9* shares orthology with one ancestral copy in rodents (124). There is also some evidence that multiple paralogs of *Prdm9* are involved in recombination hotspot regulation in cattle (125). It has been shown that changes in *Prdm9* gene dosage affect fertility in mice (126, 127), suggesting that PRDM9 may be limiting in some contexts. Theoretical models also predict that the loss of fitness induced by the erosion of PRDM9 targets could be compensated by increased gene dosage (47, 50). Thus, the duplication of a *Prdm9* allele might be temporarily advantageous when the amount of its target motifs starts to become too low in the genome. However, this benefit is expected to be only transient. This could explain why in the twelve salmonid species analyzed, each genome contained a single full length, non-pseudogenized copy of *Prdm9*. The succession of duplications and losses reported here in salmonids and previously described in mammals contributes to the apparent instability of *Prdm9* at the macro-evolutionary timescale.

### The reinforced PRDM9 paradox

This study uncovers a remarkable similarity in the recombination landscape regulation between salmonids and mammals. The main conclusion is that the PRDM9 system most likely existed in the common ancestor to vertebrates, and might be even older. Certainly, it is not a mammalian oddity. Impressively, the ultra-fast Red Queen-driven evolution of *Prdm9* and its binding motifs has been around for more than 400 My, in several vertebrate lineages (57). This implies many thousands of amino acid substitutions per site in the ZF array (57). Our results highlight the many open questions about this remarkable system, particularly the question of its long-term maintenance, which is now demonstrated. *Prdm9* can evidently be lost, for instance in birds and canids. Its continuous presence in most mammals, snakes, salmonids, and presumably many other taxa might be partly explained by the molecular mechanisms of PRDM9-dependent and PRDM9-independent recombination. The net output of these two processes is the same: CO formation. However, there may be differences in the kinetics or efficiency of DNA DSB formation and repair and thus in the robustness of CO control. This is suggested by the interaction between PRDM9 and ZCWPW1, a protein that facilitates DNA DSB repair (128–130), and by the co-evolution of *Prdm9* with genes involved in DNA DSB repair and CO formation, such as *Tex15* and *Fbxo47* (56). If PRDM9 activity is linked to other molecular processes, its loss without loss of fertility may require several mutational events. Interestingly, an intermediate context, suggesting a reduction of PRDM9 activity, has been observed in the corn snake *Pantherophis guttatus*. Specifically, Hoge et al (61) reported elevated recombination rates at PRDM9 binding sites and promoter-like features, introducing the idea of a “tug of war” between *Prdm9* and the default, *Prdm9*-independent, system. A recent study in mammals also showed that many species with *Prdm9* make substantial use of default sites, unlike humans and mice (131). The relative efficiency of the *Prdm9*-independent and *Prdm9*-dependent pathways presumably evolves and differs among species. When the *Prdm9*-independent pathway is sufficiently efficient, the conditions might be met for losing *Prdm9* irreversibly. The characterization of recombination patterns and mechanisms in species with and without *Prdm9* should help to understand the paradox of its peculiar evolution.

## Material and Methods

### Phylogenetic analysis of PRDM9 paralogs in salmonids

We investigated the presence of full-length PRDM9 in twelve species from the three salmonid subfamilies (*Coregoninae*, *Thymallinae*, *Salmoninae*). We searched for *Prdm9*-related genes by homology using the full-length copy of *O. kisutch* (coho salmon), focusing on the three PRDM9 canonical domains: KRAB (encoded by 2 exons), SSXRD (1 exon) and SET (3 exons). We obtained coho salmon PRDM9 from a nearly full-length coding sequence annotated in the RefSeq database (XP_020359152.1), complemented in its 3’ end using a cDNA identified in a brain RNA-seq dataset sequenced with PacBio long reads (SRR10185924.264665.1). We used this reference sequence to identify, with BLAST, *Prdm9* homologs in the whole genome assembly of lake whitefish (*Coregonus clupeaformis*), European grayling (*Thymallus thymallus*), huchen (*Hucho hucho*), coho salmon (*O. kisutch*), rainbow trout (*O. mykiss*), chinook salmon (*Oncorhynchus tschawytscha*), chum salmon (*Oncorhynchus keta*), red salmon (*Oncorhynchus nerka*), pink salmon (*Oncorhynchus gorbuscha*), Atlantic salmon (*Salmo salar*), brown trout (*Salmo trutta*), and lake trout (*Salvelinus namaycush*), and also of northern pike (*Esox lucius*, Esocidae), a closely related outgroup. As we obtained multiple hits, we filtered out copies containing only one of the six exons. We compared candidates to all *PRDM*-related genes annotated in the human and mouse genomes in Ensembl to exclude non-*Prdm9* homologs. We aligned the retained exons separately using Macse (v2.06) (66), to take into account potential frameshifts and stop codons. We manually examined and edited the alignments before concatenating exons of the same copy using AMAS concat (67). Several paralogous copies of *Prdm9* are expected to result from the two WGD events that occurred in the common ancestor of teleosts (Ts3R, *c.a.* 320 Mya) and salmonids (Ss4R, *c.a.* 90 Mya), respectively (63–65). We used the location of these paralogs on pairs of ohnologous chromosomes resulting from the most recent Ss4R duplication to trace the evolutionary history of *Prdm9* duplications, retention and losses. We built the maximum-likelihood phylogeny of the three canonical domains using IQ-TREE (68) based on amino acid alignments, using ultrafast bootstrap with 1000 replicates. Lastly, to identify functional *Prdm9* copies with sequence orthology to the 10 exons found in human and mouse *Prdm9* (69), we predicted the structure of each gene copy surrounded by 10 kb flanking regions using Genewise (v2.4.1) (70). We selected representative paralogous sequences across the obtained *Prdm9* phylogenetic tree to perform a sequence similarity-based annotation of the copies in each species. See details in Supporting information (S1 Methods).

### Analysis of PRDM9 ZF diversity in rainbow trout and Atlantic salmon

We characterized the allelic diversity of the ZF domain of *Prdm9α* copies in two species with different functional α-paralogs: Atlantic salmon (*S. salar*) and rainbow trout (*O. mykiss*). We focused on *Prdm9α* because a previous study showed that in teleost fish*, Prdm9β* copies lack the KRAB and SSXRD domains, have a slowly evolving ZF domain and carry a presumably inactive SET domain (19).

First, to validate the presence of expressed *Prdm9α* copies, we inferred the expression levels of multiple *Prdm9α* paralogs in immature testes from the *Salmo* and *Oncorhynchus* genera, using publicly available RNA-seq data from the Sequence Read Archive (SRA) repository. Specifically, we analyzed data from two *S. salar* samples (SRR1422872 and SRR9593306) and two *O. kisutch* samples (SRR8177981 and SRR2157188). Our analysis revealed high expression of two distinct *Prdm9α* paralogs in both genera that were previously identified in the phylogenetic analysis. We then sequenced the *Prdm9α* paralogs *α1.a.2* (full length, chromosome 5, n=26) and *α*2.2 (partial, chromosome 17, n=20) in *S. salar*, and the *Prdm9α* paralogs *α1.a.1* (full length, chromosome 31, n=23) and *α2.2* (partial, chromosome 7, n=20) in *O. mykiss*.

We used wild Atlantic salmon samples from Normandy (France) and rainbow trout samples from an INRAE selected strain (S1 Table). We extracted genomic DNA from fin clips stored in ethanol at −20°C, using the Qiagen DNAeasy Kit following the manufacturer’s instructions. We measured DNA concentration and purity with a Nanodrop-1000 Spectrophotometer (Thermo Fisher Scientific), and assessed DNA quality by agarose gel electrophoresis. We designed primers using NCBI Primer Blast, ensuring specificity against the reference assemblies. Primers targeted the ZF sequence encoded in the last exon of the gene, framed by the flanking arms of the array, avoiding any specificity of the paralogous loci (S2 Table). We carried out PCR reactions using 1X Phusion HF buffer, 200 µM dNTPs, 0.5 µM forward primer, 0.5 µM reverse primer, 3% DMSO, 2.5-10 ng template, and 0.5 units of Phusion Polymerase (NEB) (total volume: 25 μl). Cycling conditions were: initial denaturation at 98 °C for 2 min followed by 35 cycles of 98 °C for 10 s, 66-70 °C for 30 s, 72 °C for 90 s, and a final elongation step at 72 °C for 3 min, followed by hold at 10 °C, in a C1000 Cycler (Bio-Rad). We examined PCR products on agarose gels and purified them using the NucleoSpin Gel and PCR clean-up kit (Machery-Nagel). We performed Sanger sequencing of single-size amplicons. Conversely, we separated by electrophoresis heterozygous samples showing two different length alleles, followed by cloning using the TOPO Blunt Cloning Kit (Invitrogen) and sequencing. Sequencing was done by Azenta-GeneWiz (Leipzig, Germany).

We assembled and aligned forward and reverse reads to the reference ZF array from *S. salar* ICSASG_v2 and *O. mykiss* USDA_OmykA_1.1, using SnapGene (v5.1.4.1 - 5.2.3). We translated contigs into amino acid sequences used to categorize individual *Prdm9*α alleles. We annotated all ZF arrays to match the C2H2 ZF motif X7-CXXC-X12-HXXXH. We reported new alleles every time we found a single amino acid variation. We aligned the DNA sequences for each allele to create a consensus sequence. We then followed (56) and (19) to compare amino acid diversity at DNA-binding residues of the ZF array (positions −1, 2, 3 and 6 of the α-helix) with diversity values at each site of the ZF array. We calculated the proportion of the total amino acid diversity (r) at DNA-binding sites as the sum of diversity at DNA-binding residues over the sum of diversity at all 28 residues of the array (see details in S1 Methods).

### Identification of DSB hotspots in rainbow trout using ChIP-sequencing

We investigated the genome-wide distribution of DMC1-bound ssDNA in *O. mykiss* testes by ChIP followed by ssDNA enrichment (DMC1-Single Strand DNA Sequencing, DMC1-SSDS). We chose three rainbow trout individuals from the pool of samples previously used to characterize PRDM9 ZF diversity. Briefly, we excised male gonads, fixed them in Bouin’s solution, and stained 7 µm-thick tissue sections with hematoxylin-eosin (Diapath) followed by mounting with afcolene mounting medium. We determined the stage of gonadal maturation by macroscopic (whole gonads) and histological (gonad sections) analyses, according to (71). As DMC1 binds to chromatin during the early stages of the meiotic prophase I, we used testes at stages III and IV from three individuals with different *Prdm9* genotypes (TAC-1: *Prdm9^1/5^*, stage IIIb; TAC-3: *Prdm9^2/6^*, stage IIIa; and RT-52: *Prdm9^1/2^*, stage IV). This allowed us to compare DSB hotspots between individuals sharing or not a *Prdm9* allele.

For H3K4me3 and H3K36me3 ChIP experiments we used the protocols described in (72, 73) with some adjustments and rabbit anti-H3K4me3 (Abcam, ab8580) and anti-H3K36me3 (Diagenode, Premium, C15410192) antibodies. For DMC1 ChIP, we used previously described methods (74, 75) and antibodies against DMC1. These antibodies were raised by immunization of a rabbit and a guinea pig with a His-tagged recombinant zebrafish DMC1 (see details in S1 Methods). All ChIP experiments were performed in duplicate. A list of the samples and antibodies used for the ChIP-seq experiments and the number of mapped reads is in S3 Table.

For H3K4me3 and H3K36me3 ChIP-seq, we generated libraries using the NEBNext Ultra II protocol for Illumina (NEB, E7645S-E7103S), with minor adjustments. For all clean-up steps in the protocol we used MinElute columns (Qiagen, 28004). For size selection, we performed agarose gel electrophoresis and excised DNA fragments of 200 to 400 bp that were purified using MinElute columns (Qiagen, 28604). For DMC1-SSDS, we generated libraries following the Illumina TruSeq protocol (Illumina, IP-202-9001DOC), with the introduction of an additional step of kinetic enrichment, as previously described (74, 75). Libraries were sequenced on a NovaSeq6000 platform (Illumina) with S4 flow cells by Novogene Europe (Cambridge, United Kingdom).

We analyzed histone modifications with the nf-core/chipseq v1.2.1 pipeline developed by (76). Briefly, we aligned the sequencing reads of all ChIP-seq experiments to the USDA_OmykA_1.1 assembly with BWA (v0.7.17-r1188). For both H3K4me3 and H3K36me3 marks, we normalized the signal based on the read coverage and by subtracting the input. We performed peak calling with MACS2 (v2.2.7.1) for both replicates, and provided an input for each sample. We assessed the histone mark enrichment at DMC1 peaks, and the enrichment of H3K36me3 signal at H3K4me3 peaks in brain using the deepTools suite (77) and the bed files produced by the AQUA-FAANG project (https://www.aqua-faang.eu/). We analyzed the DMC1-SSDS data as described in (75), with some implementations described in (78), using the hotSSDS pipeline (version 1.0). We mapped reads with the modified BWA algorithm (BWA *Right Align*), developed to align and recover ssDNA fragments, as described by (74). We normalized the signal based on the library size and the type 1 ssDNA fragments. We performed peak calling with MACS2 (v2.2.7.1) and relaxed conditions for each of the two replicates, and provided an input control. We carried out an Irreproducible Discovery Rate (IDR) analysis to identify reproducible enriched regions. Then, we used these peaks as DSB hotspots (see details in S1 Methods). We used the final peaks to check the distribution of ssDNA type 1 signal at DSB hotspots.

We explored the relationship between H3K4me3 and H3K36me3 signal distribution by calculating the correlation between H3K4me3 and H3K36me3 read enrichment at DSB hotspots in the RT-52 sample, of the H3K4me3 read enrichment between the TAC-1 and TAC-3 samples at the RT-52 DSB hotspots, and of the H3K36me3 read enrichment between the TAC-1 and TAC-3 samples at RT-52 DSB hotspots. Lastly, we assessed the proportion of DSB hotspots overlapping with H3K4me3 and H3K36me3 peaks.

### Reconstruction of population recombination landscapes in three salmonid species

#### Whole-genome resequencing data

To reconstruct population-based recombination landscapes, we collected high coverage whole-genome resequencing data from five natural populations of three salmonid species from the SRA database: coho salmon (*O. kisutch*), rainbow trout (*O. mykiss*) and Atlantic salmon (*S. salar*). We used ∼20 individuals per population as recommended (79). We retrieved 20 genomes of the Southern British Columbia population of coho salmon (80), 22 genomes of rainbow trout from North West America (81), and 60 genomes of three populations of Atlantic salmon belonging to the two major lineages from North America and Europe (82): 20 from the Gaspesie Peninsula in Canada (GP population thereafter), 20 from the North Sea (NS population), and 20 from the Barents Sea in Norway (BS population). Sample accession numbers and locations are in S4 Table and S1 Fig.

#### Variant calling

Variants and genotypes called by (80) using GATK were used for *O. kisutch*. We followed the same methodology for variant calling and genotyping in *O. mykiss* and *S. salar*, using the GATK best-practice pipeline (> v3.8-0, see S5 Table for the detailed versions of the programs (83, 84)). First, we aligned paired-end reads to their reference genome (Okis_V1, GCF_002021735.1; Omyk_1.0, GCF_002163495.1; Ssal_v3.1, GCF_905237065.1) using BWA-MEM (v0.7.17, Li and Durbin, 2009; -*M* option), yielding an average read coverage depth per sample of 29.54x, 24.87x and 9.97x for *O. kisutch*, *O. mykiss* and *S. salar*, respectively (S6 Table). We used Picard (> v2.18.29) to mark PCR duplicates and add read groups. Then, we performed variant calling separately for each individual using HaplotypeCaller before joint genotyping with GenotypeGVCFs. In total, we analyzed 9,590,270, 39,601,311 and 27,061,466 single nucleotide polymorphisms (SNPs) for *O. kisutch*, *O. mykiss* and *S. salar*, respectively.

After genotyping, we removed variants within 5 bp of an indel with the Bcftools filter (v 1.9; Li, 2011; *-g 5*). We filtered low-quality SNPs with Vcftools (> v 0.1.16) (85), keeping only biallelic SNPs, and excluding genotypes with low quality scores (*--minGP 20*) and SNPs with >10% of missing genotypes (*--max-missing 0.9*). For the *S. salar* dataset, we set the missingness threshold at 50% to take into account the lower sequencing coverage depth in this species. To remove the effect of poorly sequenced and duplicated regions, we kept only sites with a mean coverage depth within the 5–95% quantiles of that species distribution. To further eliminate shared excesses of heterozygosity due to residual tetrasomy or contaminations, we applied a Hardy-Weinberg equilibrium filter with a *p*-value exclusion threshold of 0.01 (*--hwe 0.01*). We removed singletons by applying a Minor Allele Count (MAC) filter with Vcftools (*--mac 2*). For *S. salar,* we used the missingness, Hardy-Weinberg and MAC filters separately for each of the three populations. After these filtering steps, we retrieved a total of 7,205,269, 16,079,097 and 5,575,430 SNPs for *O. kisutch*, *O. mykiss* and *S. salar*, respectively (S6 Table).

#### Variant phasing and orientation

We used the read-based phasing approach in WhatsHap (> v0.18) (86) to identify phase blocks from paired-end reads that overlapped with neighboring individual heterozygous positions. This allowed us to locally resolve the physical phase of 73.45%, 76.98% and 7.32% of variants for *O. kisutch*, *O. mykiss* and *S. salar*, respectively. Then, we performed the statistical phasing of pre-phased blocks with SHAPEIT4 (> v4.2.1, (87), default settings) in each species, assuming a uniform recombination rate of 3 cM/Mb (representative of the average recombination rates in teleosts, (11)) and using the effective population size estimated from the mean nucleotide diversity of each chromosome calculated with Vcftools.

We inferred ancestral allelic state probabilities for the set of retained variants of each species with the maximum likelihood method implemented in est-sfs (v2.04, Kimura-2-parameter substitution model)(88), using three outgroups per species chosen among the available salmonid reference genomes (see details in S1 Methods). The method uses the ingroup allele frequencies and the allelic states of the outgroups to infer ancestral allelic state, taking into account the phylogenetic relationships between ingroup and outgroup species (89).

#### Estimation of linkage disequilibrium (LD)-based recombination rates

For each of the five population datasets (*O. kisutch*, *O. mykiss*, and *S. salar* GP, BS and NS populations, S1 Fig), we estimated the population-scaled recombination rate parameter *⍴* (*⍴* = 4*N_e_r*, where *N_e_* is the effective population size and *r* the recombination rate in M/bp) using LDhelmet (v1.19) (13). LDhelmet relies on a reversible-jump Markov Chain Monte Carlo algorithm to infer the *⍴* value between every pair of consecutive SNPs. Variant orientation was provided using the probabilities, estimated by est-sfs, that the major and minor alleles were ancestral, and a transition matrix was computed following (13). We run LDhelmet five times independently for each population. For each chromosome, we created the haplotype configuration files with the find_conf function using the recommended window size of 50 SNPs. We created the likelihood look-up tables once for the five runs with the table_gen function using the recommended grid for the population recombination rate (*ρ*/pb) (*i.e. ρ* from 0 to 10 by increments of 0.1, then from 10 to 100 by increments of 1), and with the Watterson *θ* = 4*N*_e_*μ* parameter of the corresponding chromosome obtained using *μ*=10^-8^. We created the Padé files using 11 Padé coefficients as recommended. We run the Monte Carlo Markov chain for 1 million iterations with a burn-in period of 100,000 and a window size of 50 SNPs, using a block penalty of 5. We checked the convergence of the five independent runs by comparing the estimated recombination values with the Spearman’s rank correlation test (Spearman’s rho > 0.96; S2 Fig). We averaged and smoothened the five runs within 2-kb, 100-kb and 1-Mb windows using custom python scripts.

We reconstructed the fine-scale recombination landscape of the European sea bass (*Dicentrarchus labrax*) to compare recombination features in salmonids with a species that lacks a complete *Prdm9* gene due to loss of the KRAB domain. We used whole-genome haplotype data obtained by phasing-by-transmission and statistical phasing (90) to infer recombination in the Atlantic sea bass population with a similar strategy, using the seabass_V1.0 genome assembly (GenBank accession number GCA_000689215.1) (S5 and S6 Tables).

#### Identification of LD-based recombination hotspots

We identified recombination hotspots from the raw recombination map inferred by LDhelmet (*i.e.* raw hotspots) and from the 2-kb smoothed recombination map (*i.e.* 2-kb hotspots) using a sliding window approach. We defined hotspots as intervals between consecutive SNPs or 2-kb windows with a relative recombination rate ≥5-fold higher than the mean recombination rate in the 50-kb flanking regions. When consecutive 2kb-windows exceeded the threshold, we retained only the window with the highest rate.

### Analysis of recombination landscapes

#### Comparison between DMC1 and LD-based recombination maps

We compared DSB hotspots mapped by DMC1-SSDS of the three pooled samples and the LD-based recombination hotspots retrieved from the recombination landscapes of *O. mykiss*. For this, we converted the genomic positions of the DSB hotspots mapped on the OmykA_1.1 assembly to the Omyk_1.0 coordinates on which we built the LD map using the Remap program from NCBI. We compared the locations of LD-hotspots and of DSB hotspots using Bedtools intersect (91).

#### Recombination at genomic features

We investigated how DSB-based (*O. mykiss*) and LD-based (three salmonid species and sea bass) recombination rates and hotspots were distributed relative to genomic features. We first retrieved the positions of genes, exons and introns from genome annotations in each species. We *de novo* identified transposable element (TE) families in each genome using RepeatModeler (v2.0.3; option -LTRStruct) (92), before mapping TEs and low complexity DNA sequences with RepeatMasker (version 4.1.3, http://www.repeatmasker.org/, options -xsmall, -nolow). We deduced intergenic regions from gene and TE locations using Bedtools *subtract*. We defined the TSS and TES as the first and last position of a gene, respectively. For each reference genome, we predicted CGIs with EMBOSS *cpgplot* (v6.6.0) (93), using the parameters *-window 500 -minlen 250 -minoe 0.6 -minpc 0*. It should be noted that the criteria that are classically used to predict CGIs in mammals and birds (CpG observed/expected ratio >0.6, GC content >50%) are not appropriate for teleost fish in which CGIs are CpG-rich but have a low GC-content (94, 95). Therefore, we predicted CGIs based only on their CpG content, without any constraint on their GC content. We confirmed that these criteria efficiently predicted TSS-associated CGIs, using whole genome DNA methylation and H3K4me3 data from rainbow trout and coho salmon (see S2 Analysis).

We investigated DSB hotspots overlap with genomic features and their distance to the nearest promoter-like feature (TSS) using Bedtools. For this, we analyzed DNA DSB distribution using DMC1-SSDS read enrichment as metric. As the coordinates of genomic features were mapped on Omyk_1.0, we converted them to the OmykA_1.1 assembly using the NCBI Remap tool.

We assessed population recombination rate (2-kb scale) variations in function of the distance to the nearest TSS (overlapping or not with a CGI) using the *distanceToNearest* function of the R package *GenomicRanges* (96). We retrieved the averaged recombination rates at each genomic feature (*i.e.* genes, exons, introns, intergenic regions, TSSs within and outside CGIs and TEs) using the *subsetByOverlaps* function of the same package. We compared recombination rates at genomic features in the five salmonid populations and in sea bass.

Lastly, we investigated the effect of SNP density, GC content and TEs on the population recombination rate variation and the presence of recombination hotspots. We calculated SNP density, TE density, and GC content in non-overlapping 100-kb windows, and compared them with the window-averaged recombination rates using the Spearman’s rank correlation test. We assessed the association of hotspots with SNP density and GC content at the 2-kb scale.

#### Comparison of LD-based landscapes between populations and species

We assessed the correlation between the 100-kb smoothed recombination maps of each of the three *S. salar* populations using the Spearman’s rank test. We identified shared hotspots between populations as overlapping 2-kb hotspots using *BedTools intersect*. To compare the recombination hotspots of the *O. kisutch* and *O. mykiss* populations, we used a reciprocal blast approach to identify homologous regions of the genome in these two species (see S1 Methods). We used random permutations to calculate the expected amount of hotspot overlap between pairs of the three *S. salar* populations and between the two *Oncorhynchus* populations. We drew random spots (same number as that of 2-kb hotspots) 100 times from the genome for each population using Bedtools *shuffle*, and we compared each of these random sets to those of the compared population to calculate the average overlap expected only by chance.

#### Identification of DNA motifs at hotspots and motif erosion

In rainbow trout, we performed motif detection analysis at DSB hotspots using the MEME Suite (97), focusing on the RT-52 dataset due to its high number of DSB hotspots (DMC1 peaks). We defined two subsets of allele-specific hotspots using Bedtools *intersect*: Allele 1 set [RT-52 DMC1 peaks (center ± 200 bp) overlapping with H3K4me3 and H3K36me3 peaks from TAC-1 (N = 300)] and Allele 2 set [RT-52 DMC1 peaks (center ± 200 bp) overlapping with H3K4me3 and H3K36me3 peaks from TAC-3 (N = 254)]. We used MEME-ChIP (98) to detect motifs. Then, we assessed motif enrichment in DSB hotspots relative to control sequences using FIMO (99) and evaluated central enrichment using CentriMo (100). We also quantified the fold-enrichment of the detected motifs at LD-based hotspots (see S1 Methods).

In Atlantic salmon, we used STREME from the MEME Suite (101) to find motifs between 10 and 20 bp in length that were enriched at LD-based hotspot positions, compared with control random sequences with a similar GC-content distribution (see S1 Methods). We retained the detected motifs as potential PRDM9-binding motifs if they were enriched ≥2-fold at the hotspots compared with the control sequences, and were found in ≥5% of hotspots. We searched for motifs associated with the hotspots of each *S. salar* population and with the shared hotspots between pairs of populations.

We then tested whether the candidate motifs showed signs of erosion between lineages by comparing the number of motifs present in the available long-read genome assemblies from five North American and seven European Atlantic salmon genomes. To take into account potential differences in assembly size, we aligned these twelve genomes with SibeliaZ (102) and retrieved collinear blocks that represented 89.5% of the whole genome alignment. We then used FIMO (99) to count motif occurrence in the aligned fraction of each genome, using a p-value cut-off of 1.0E-7. To assess the statistical significance of motif erosion in a given lineage, we obtained a null distribution of the between-lineage difference in motif occurrence by running FIMO on 100 random permutations of the candidate motif matrix.

## Data Accessibility

Sequencing data from ChIPSeq will be archived at the Gene Expression Omnibus (GEO) under accession GSExxx. Sequencing data will be deposited in the GenBank Sequence Read Archive under the accession code BioProject ID PRJNAXXXXXX, and a file containing scripts and commands used for all bioinformatic analyses will be provided following acceptance.

## Acknowledgments

We thank the European project AQUA-FAANG for sharing unpublished epigenetic data of *O. mykiss* and *S. salar*, and the Aqua Genome project for providing published Wild Atlantic salmon genome sequencing data. We particularly thank Sigbjørn Lien, Marie-Odile Baudement and Yann Guiguen for helpful discussions, and Gareth Gillard for bioinformatic analysis of AQUA-FAANG data. We also thank Guillaume Evanno for providing wild Atlantic salmon samples, and Rajalekshmi Navaryana Sarna for help with *Prdm9* genotyping. We are grateful to Ben Coop and Eric Rondeau for providing us with the re-sequenced genomes and genotype dataset of coho salmon. We thank Louis Bernatchez, Eric Normandeau and Maeva Leitwein for providing the whole genome bisulfite sequencing methylation data of coho salmon. We also thank Thomas Brazier, Nicolas Lartillot and Carina Mugal for helpful discussions and feedback.

## References

1. Hunter N. Meiotic Recombination: The Essence of Heredity. Cold Spring Harb Perspect Biol. 2015;7(12).

2. Zickler D, Kleckner N. Meiosis: Dances Between Homologs. Annu Rev Genet. 2023.

3. Nagaoka SI, Hassold TJ, Hunt PA. Human aneuploidy: mechanisms and new insights into an age-old problem. Nat Rev Genet. 2012;13(7):493–504.

4. Coop G, Przeworski M. An evolutionary view of human recombination. Nat Rev Genet. 2007;8(1):23–34.

5. Otto SP, Lenormand T. Resolving the paradox of sex and recombination. Nat Rev Genet. 2002;3(4):252–61.

6. Charlesworth B, Morgan MT, Charlesworth D. The effect of deleterious mutations on neutral molecular variation. Genetics. 1993;134(4):1289–303.

7. Ritz KR, Noor MAF, Singh ND. Variation in Recombination Rate: Adaptive or Not? Trends Genet. 2017;33(5):364–74.

8. Smith JM, Haigh J. The hitch-hiking effect of a favourable gene. Genetical research. 1974;23(1):23–35.

9. Pazhayam NM, Turcotte CA, Sekelsky J. Meiotic Crossover Patterning. Frontiers in cell and developmental biology. 2021;9:681123.

10. Penalba JV, Wolf JBW. From molecules to populations: appreciating and estimating recombination rate variation. Nat Rev Genet. 2020;21(8):476–92.

11. Stapley J, Feulner PGD, Johnston SE, Santure AW, Smadja CM. Variation in recombination frequency and distribution across eukaryotes: patterns and processes. Philos Trans R Soc Lond B Biol Sci. 2017;372(1736).

12. Haenel Q, Laurentino TG, Roesti M, Berner D. Meta-analysis of chromosome-scale crossover rate variation in eukaryotes and its significance to evolutionary genomics. Mol Ecol. 2018;27(11):2477–97.

13. Chan AH, Jenkins PA, Song YS. Genome-Wide Fine-Scale Recombination Rate Variation in Drosophila melanogaster. PLoS Genet. 2012;8(12):e1003090.

14. Kaur T, Rockman MV. Crossover heterogeneity in the absence of hotspots in Caenorhabditis elegans. Genetics. 2014;196(1):137–48.

15. Wallberg A, Glemin S, Webster MT. Extreme Recombination Frequencies Shape Genome Variation and Evolution in the Honeybee, Apis mellifera. PLoS Genet. 2015;11(4):e1005189.

16. Jeffreys AJ, Holloway JK, Kauppi L, May CA, Neumann R, Slingsby MT, et al. Meiotic recombination hot spots and human DNA diversity. Philos Trans R Soc Lond B Biol Sci. 2004;359(1441):141-52.

17. McVean GA, Myers SR, Hunt S, Deloukas P, Bentley DR, Donnelly P. The fine-scale structure of recombination rate variation in the human genome. Science. 2004;304(5670):581-4.

18. Auton A, Rui Li Y, Kidd J, Oliveira K, Nadel J, Holloway JK, et al. Genetic Recombination Is Targeted towards Gene Promoter Regions in Dogs. PLoS Genet. 2013;9(12):e1003984.

19. Baker Z, Schumer M, Haba Y, Bashkirova L, Holland C, Rosenthal GG, et al. Repeated losses of PRDM9-directed recombination despite the conservation of PRDM9 across vertebrates. eLife. 2017;6.

20. Choi K, Zhao X, Tock AJ, Lambing C, Underwood CJ, Hardcastle TJ, et al. Nucleosomes and DNA methylation shape meiotic DSB frequency in Arabidopsis thaliana transposons and gene regulatory regions. Genome Res. 2018;28(4):532–46.

21. Fowler KR, Sasaki M, Milman N, Keeney S, Smith GR. Evolutionarily diverse determinants of meiotic DNA break and recombination landscapes across the genome. Genome research. 2014;24(10):1650–64.

22. Kawakami T, Mugal CF, Suh A, Nater A, Burri R, Smeds L, et al. Whole-genome patterns of linkage disequilibrium across flycatcher populations clarify the causes and consequences of fine-scale recombination rate variation in birds. Mol Ecol. 2017;26(16):4158–72.

23. Lam I, Keeney S. Nonparadoxical evolutionary stability of the recombination initiation landscape in yeast. Science. 2015;350(6263):932-7.

24. Pan J, Sasaki M, Kniewel R, Murakami H, Blitzblau HG, Tischfield SE, et al. A Hierarchical Combination of Factors Shapes the Genome-wide Topography of Yeast Meiotic Recombination Initiation. Cell. 2011;144(5):719–31.

25. Singhal S, Leffler EM, Sannareddy K, Turner I, Venn O, Hooper DM, et al. Stable recombination hotspots in birds. Science. 2015;350(6263):928-32.

26. Choi K, Henderson IR. Meiotic recombination hotspots - a comparative view. Plant J. 2015;83(1):52–61.

27. Axelsson E, Webster MT, Ratnakumar A, Ponting CP, Lindblad-Toh K. Death of PRDM9 coincides with stabilization of the recombination landscape in the dog genome. Genome Res. 2012;22(1):51–63.

28. Dutreux F, Dutta A, Peltier E, Bibi-Triki S, Friedrich A, Llorente B, et al. Lessons from the meiotic recombination landscape of the ZMM deficient budding yeast Lachancea waltii. PLoS Genet. 2023;19(1):e1010592.

29. Auton A, Fledel-Alon A, Pfeifer S, Venn O, Segurel L, Street T, et al. A Fine-Scale Chimpanzee Genetic Map from Population Sequencing. Science. 2012;336(6078):193-8.

30. Brick K, Smagulova F, Khil P, Camerini-Otero RD, Petukhova GV. Genetic recombination is directed away from functional genomic elements in mice. Nature. 2012;485(7400):642-5.

31. Coop G, Wen X, Ober C, Pritchard JK, Przeworski M. High-Resolution Mapping of Crossovers Reveals Extensive Variation in Fine-Scale Recombination Patterns Among Humans. Science. 2008;319:1395–8.

32. Myers S, Bowden R, Tumian A, Bontrop RE, Freeman C, MacFie TS, et al. Drive against hotspot motifs in primates implicates the PRDM9 gene in meiotic recombination. Science. 2010;327(5967):876-9.

33. Pratto F, Brick K, Khil P, Smagulova F, Petukhova GV, Camerini-Otero RD. DNA recombination. Recombination initiation maps of individual human genomes. Science. 2014;346(6211):1256442.

34. Smagulova F, Brick K, Pu Y, Camerini-Otero RD, Petukhova GV. The evolutionary turnover of recombination hot spots contributes to speciation in mice. Genes Dev. 2016;30(3):266–80.

35. Wooldridge LK, Dumont BL. Rapid Evolution of the Fine-scale Recombination Landscape in Wild House Mouse (Mus musculus) Populations. Mol Biol Evol. 2023;40(1).

36. Baudat F, Buard J, Grey C, Fledel-Alon A, Ober C, Przeworski M, et al. PRDM9 is a major determinant of meiotic recombination hotspots in humans and mice. Science. 2010;327(5967):836-40.

37. Parvanov ED, Petkov PM, Paigen K. Prdm9 controls activation of mammalian recombination hotspots. Science. 2010;327(5967):835.

38. Grey C, Baudat F, de Massy B. PRDM9, a driver of the genetic map. PLoS Genet. 2018;14(8):e1007479.

39. Baker CL, Kajita S, Walker M, Saxl RL, Raghupathy N, Choi K, et al. PRDM9 Drives Evolutionary Erosion of Hotspots in Mus musculus through Haplotype-Specific Initiation of Meiotic Recombination. PLoS Genet. 2015;11(1):e1004916.

40. Lesecque Y, Glemin S, Lartillot N, Mouchiroud D, Duret L. The red queen model of recombination hotspots evolution in the light of archaic and modern human genomes. PLoS Genet. 2014;10(11):e1004790.

41. Alleva B, Brick K, Pratto F, Huang M, Camerini-Otero RD. Cataloging Human PRDM9 Allelic Variation Using Long-Read Sequencing Reveals PRDM9 Population Specificity and Two Distinct Groupings of Related Alleles. Frontiers in cell and developmental biology. 2021;9:675286.

42. Berg IL, Neumann R, Lam KW, Sarbajna S, Odenthal-Hesse L, May CA, et al. PRDM9 variation strongly influences recombination hot-spot activity and meiotic instability in humans. Nat Genet. 2010;42(10):859–63.

43. Buard J, Rivals E, Dunoyer de Segonzac D, Garres C, Caminade P, de Massy B, et al. Diversity of Prdm9 Zinc Finger Array in Wild Mice Unravels New Facets of the Evolutionary Turnover of this Coding Minisatellite. PLoS One. 2014;9(1):e85021.

44. Damm E, Ullrich KK, Amos WB, Odenthal-Hesse L. Evolution of the recombination regulator PRDM9 in minke whales. BMC Genomics. 2022;23(1):212.

45. Kono H, Tamura M, Osada N, Suzuki H, Abe K, Moriwaki K, et al. Prdm9 polymorphism unveils mouse evolutionary tracks. DNA Res. 2014;21(3):315–26.

46. Schwartz JJ, Roach DJ, Thomas JH, Shendure J. Primate evolution of the recombination regulator PRDM9. Nat Commun. 2014;5:4370.

47. Baker Z, Przeworski M, Sella G. Down the Penrose stairs, or how selection for fewer recombination hotspots maintains their existence. eLife. 2023;12.

48. Latrille T, Duret L, Lartillot N. The Red Queen model of recombination hot-spot evolution: a theoretical investigation. Philos Trans R Soc Lond B Biol Sci. 2017;372(1736).

49. Ubeda F, Wilkins JF. The Red Queen theory of recombination hotspots. J Evol Biol. 2011;24(3):541–53.

50. Genestier A, Duret L, Lartillot N. Bridging the gap between the evolutionary dynamics and the molecular mechanisms of meiosis : a model based exploration of the <em>PRDM9</em> intra-genomic Red Queen. bioRxiv. 2023:2023.03.08.531712.

51. Davies B, Hatton E, Altemose N, Hussin JG, Pratto F, Zhang G, et al. Re-engineering the zinc fingers of PRDM9 reverses hybrid sterility in mice. Nature. 2016;530(7589):171-6.

52. Forejt J, Jansa P, Parvanov E. Hybrid sterility genes in mice (Mus musculus): a peculiar case of PRDM9 incompatibility. Trends Genet. 2021;37(12):1095–108.

53. Gregorova S, Gergelits V, Chvatalova I, Bhattacharyya T, Valiskova B, Fotopulosova V, et al. Modulation of Prdm9-controlled meiotic chromosome asynapsis overrides hybrid sterility in mice. eLife. 2018;7.

54. Ponting CP. What are the genomic drivers of the rapid evolution of PRDM9? Trends Genet. 2011;27(5):165–71.

55. Sandor C, Li W, Coppieters W, Druet T, Charlier C, Georges M. Genetic Variants in REC8, RNF212, and PRDM9 Influence Male Recombination in Cattle. PLoS Genet. 2012;8(7):e1002854.

56. Cavassim MIA, Baker Z, Hoge C, Schierup MH, Schumer M, Przeworski M. PRDM9 losses in vertebrates are coupled to those of paralogs ZCWPW1 and ZCWPW2. Proc Natl Acad Sci U S A. 2022;119(9).

57. Oliver PL, Goodstadt L, Bayes JJ, Birtle Z, Roach KC, Phadnis N, et al. Accelerated Evolution of the Prdm9 Speciation Gene across Diverse Metazoan Taxa. PLoS Genet. 2009;5(12):e1000753.

58. Mihola O, Landa V, Pratto F, Brick K, Kobets T, Kusari F, et al. Rat PRDM9 shapes recombination landscapes, duration of meiosis, gametogenesis, and age of fertility. BMC Biol. 2021;19(1):86.

59. Shanfelter AF, Archambeault SL, White MA. Divergent Fine-Scale Recombination Landscapes between a Freshwater and Marine Population of Threespine Stickleback Fish. Genome Biol Evol. 2019;11(6):1573–85.

60. Versoza CJ, Rivera JA, Rosenblum EB, Vital-Garcia C, Hews DK, Pfeifer SP. The recombination landscapes of spiny lizards (genus Sceloporus). G3 (Bethesda). 2022;12(2).

61. Hoge C, de Manuel M, Mahgoub M, Okami N, Fuller Z, Banerjee S, et al. Patterns of recombination in snakes reveal a tug of war between PRDM9 and promoter-like features. bioRxiv. 2023.

62. Schield DR, Pasquesi GIM, Perry BW, Adams RH, Nikolakis ZL, Westfall AK, et al. Snake Recombination Landscapes Are Concentrated in Functional Regions despite PRDM9. Mol Biol Evol. 2020;37(5):1272–94.

63. Christoffels A, Koh EG, Chia JM, Brenner S, Aparicio S, Venkatesh B. Fugu genome analysis provides evidence for a whole-genome duplication early during the evolution of ray-finned fishes. Mol Biol Evol. 2004;21(6):1146–51.

64. Macqueen DJ, Johnston IA. A well-constrained estimate for the timing of the salmonid whole genome duplication reveals major decoupling from species diversification. Proc Biol Sci. 2014;281(1778):20132881.

65. Vandepoele K, De Vos W, Taylor JS, Meyer A, Van de Peer Y. Major events in the genome evolution of vertebrates: paranome age and size differ considerably between ray-finned fishes and land vertebrates. Proc Natl Acad Sci U S A. 2004;101(6):1638–43.

66. Ranwez V, Douzery EJP, Cambon C, Chantret N, Delsuc F. MACSE v2: Toolkit for the Alignment of Coding Sequences Accounting for Frameshifts and Stop Codons. Mol Biol Evol. 2018;35(10):2582–4.

67. Borowiec ML. AMAS: a fast tool for alignment manipulation and computing of summary statistics. PeerJ. 2016;4:e1660.

68. Nguyen LT, Schmidt HA, von Haeseler A, Minh BQ. IQ-TREE: a fast and effective stochastic algorithm for estimating maximum-likelihood phylogenies. Mol Biol Evol. 2015;32(1):268–74.

69. Hayashi K, Yoshida K, Matsui Y. A histone H3 methyltransferase controls epigenetic events required for meiotic prophase. Nature. 2005;438(7066):374-8.

70. Birney E, Clamp M, Durbin R. GeneWise and Genomewise. Genome Res. 2004;14(5):988–95.

71. Billard R, Solari A, Escaffre AM. [Method for the quantitative analysis of spermatogenesis in teleost fish]. Ann Biol Anim Biochim Biophys. 1974;14(1):87–104.

72. Diagouraga B, Clement JAJ, Duret L, Kadlec J, de Massy B, Baudat F. PRDM9 Methyltransferase Activity Is Essential for Meiotic DNA Double-Strand Break Formation at Its Binding Sites. Mol Cell. 2018;69(5):853–65 e6.

73. Tardat M, Brustel J, Kirsh O, Lefevbre C, Callanan M, Sardet C, et al. The histone H4 Lys 20 methyltransferase PR-Set7 regulates replication origins in mammalian cells. Nat Cell Biol. 2010;12(11):1086–93.

74. Brick K, Pratto F, Sun CY, Camerini-Otero RD, Petukhova G. Analysis of Meiotic Double-Strand Break Initiation in Mammals. Methods Enzymol. 2018;601:391–418.

75. Khil PP, Smagulova F, Brick KM, Camerini-Otero RD, Petukhova GV. Sensitive mapping of recombination hotspots using sequencing-based detection of ssDNA. Genome Res. 2012;22(5):957–65.

76. Ewels PA, Peltzer A, Fillinger S, Patel H, Alneberg J, Wilm A, et al. The nf-core framework for community-curated bioinformatics pipelines. Nat Biotechnol. 2020;38(3):276–8.

77. Ramírez F, Ryan DP, Grüning B, Bhardwaj V, Kilpert F, Richter AS, et al. deepTools2: a next generation web server for deep-sequencing data analysis. Nucleic Acids Res. 2016;44(W1):W160–5.

78. Auffret P, de Massy B, Clement JAJ. Mapping Meiotic DNA Breaks: Two Fully-Automated Pipelines to Analyze Single-Strand DNA Sequencing Data, hotSSDS and hotSSDS-extra. Methods Mol Biol. 2024;2770:227–61.

79. Raynaud M, Gagnaire P-A, Galtier N. Performance and limitations of linkage-disequilibrium-based methods for inferring the genomic landscape of recombination and detecting hotspots: a simulation study. Peer Community Journal. 2023;3.

80. Rondeau EB, Christensen KA, Minkley DR, Leong JS, Chan MTT, Despins CA, et al. Population-size history inferences from the coho salmon (Oncorhynchus kisutch) genome. G3 (Bethesda). 2023;13(4).

81. Gao G, Nome T, Pearse DE, Moen T, Naish KA, Thorgaard GH, et al. A New Single Nucleotide Polymorphism Database for Rainbow Trout Generated Through Whole Genome Resequencing. Front Genet. 2018;9:147.

82. Bertolotti AC, Layer RM, Gundappa MK, Gallagher MD, Pehlivanoglu E, Nome T, et al. The structural variation landscape in 492 Atlantic salmon genomes. Nat Commun. 2020;11(1):5176.

83. McKenna A, Hanna M, Banks E, Sivachenko A, Cibulskis K, Kernytsky A, et al. The Genome Analysis Toolkit: a MapReduce framework for analyzing next-generation DNA sequencing data. Genome Res. 2010;20(9):1297–303.

84. Van der Auwera GA, Carneiro MO, Hartl C, Poplin R, Del Angel G, Levy-Moonshine A, et al. From FastQ data to high confidence variant calls: the Genome Analysis Toolkit best practices pipeline. Current protocols in bioinformatics. 2013;43(1110):11.0.1-.0.33.

85. Danecek P, Auton A, Abecasis G, Albers CA, Banks E, DePristo MA, et al. The variant call format and VCFtools. Bioinformatics. 2011;27(15):2156–8.

86. Martin M, Patterson M, Garg S, Fischer SO, Pisanti N, Klau GW, et al. WhatsHap: fast and accurate read-based phasing. bioRxiv. 2016:085050.

87. Delaneau O, Zagury JF, Robinson MR, Marchini JL, Dermitzakis ET. Accurate, scalable and integrative haplotype estimation. Nat Commun. 2019;10(1):5436.

88. Keightley PD, Jackson BC. Inferring the Probability of the Derived vs. the Ancestral Allelic State at a Polymorphic Site. Genetics. 2018;209(3):897–906.

89. Crespi BJ, Teo R. Comparative phylogenetic analysis of the evolution of semelparity and life history in salmonid fishes. Evolution. 2002;56(5):1008–20.

90. Duranton M, Allal F, Valière S, Bouchez O, Bonhomme F, Gagnaire PA. The contribution of ancient admixture to reproductive isolation between European sea bass lineages. Evol Lett. 2020;4(3):226–42.

91. Quinlan AR, Hall IM. BEDTools: a flexible suite of utilities for comparing genomic features. Bioinformatics. 2010;26(6):841–2.

92. Flynn JM, Hubley R, Goubert C, Rosen J, Clark AG, Feschotte C, et al. RepeatModeler2 for automated genomic discovery of transposable element families. Proc Natl Acad Sci U S A. 2020;117(17):9451–7.

93. Larsen F, Gundersen G, Lopez R, Prydz H. CpG islands as gene markers in the human genome. Genomics. 1992;13(4):1095–107.

94. Cross S, Kovarik P, Schmidtke J, Bird A. Non-methylated islands in fish genomes are GC-poor. Nucleic Acids Res. 1991;19(7):1469–74.

95. Long HK, Sims D, Heger A, Blackledge NP, Kutter C, Wright ML, et al. Epigenetic conservation at gene regulatory elements revealed by non-methylated DNA profiling in seven vertebrates. eLife. 2013;2:e00348.

96. Lawrence M, Huber W, Pagès H, Aboyoun P, Carlson M, Gentleman R, et al. Software for computing and annotating genomic ranges. PLoS Comput Biol. 2013;9(8):e1003118.

97. Bailey TL, Johnson J, Grant CE, Noble WS. The MEME Suite. Nucleic Acids Res. 2015;43(W1):W39–49.

98. Machanick P, Bailey TL. MEME-ChIP: motif analysis of large DNA datasets. Bioinformatics. 2011;27(12):1696–7.

99. Grant CE, Bailey TL, Noble WS. FIMO: scanning for occurrences of a given motif. Bioinformatics. 2011;27(7):1017–8.

100. Bailey TL, Machanick P. Inferring direct DNA binding from ChIP-seq. Nucleic Acids Res. 2012;40(17):e128.

101. Bailey TL. STREME: accurate and versatile sequence motif discovery. Bioinformatics. 2021;37(18):2834–40.

102. Minkin I, Medvedev P. Scalable multiple whole-genome alignment and locally collinear block construction with SibeliaZ. Nat Commun. 2020;11(1):6327.

103. Lien S, Koop BF, Sandve SR, Miller JR, Kent MP, Nome T, et al. The Atlantic salmon genome provides insights into rediploidization. Nature. 2016;533(7602):200-5.

104. Pearse DE, Barson NJ, Nome T, Gao G, Campbell MA, Abadía-Cardoso A, et al. Sex-dependent dominance maintains migration supergene in rainbow trout. Nature ecology & evolution. 2019;3(12):1731–42.

105. Sutherland BJG, Gosselin T, Normandeau E, Lamothe M, Isabel N, Audet C, et al. Salmonid Chromosome Evolution as Revealed by a Novel Method for Comparing RADseq Linkage Maps. Genome Biol Evol. 2016;8(12):3600–17.

106. Wu H, Mathioudakis N, Diagouraga B, Dong A, Dombrovski L, Baudat F, et al. Molecular Basis for the Regulation of the H3K4 Methyltransferase Activity of PRDM9. Cell reports. 2013;5(1):13–20.

107. Brekke C, Johnston SE, Knutsen TM, Berg P. Genetic architecture of individual meiotic crossover rate and distribution in a large Atlantic Salmon (<em>Salmo salar)</em> breeding population. bioRxiv. 2023:2023.06.07.543993.

108. Paigen K, Petkov PM. PRDM9 and Its Role in Genetic Recombination. Trends Genet. 2018;34(4):291–300.

109. Grey C, Clement JA, Buard J, Leblanc B, Gut I, Gut M, et al. In vivo binding of PRDM9 reveals interactions with noncanonical genomic sites. Genome Res. 2017;27(4):580–90.

110. Powers NR, Parvanov ED, Baker CL, Walker M, Petkov PM, Paigen K. The Meiotic Recombination Activator PRDM9 Trimethylates Both H3K36 and H3K4 at Recombination Hotspots In Vivo. PLoS Genet. 2016;12(6):e1006146.

111. Myers S, Bottolo L, Freeman C, McVean G, Donnelly P. A fine-scale map of recombination rates and hotspots across the human genome. Science. 2005;310(5746):321-4.

112. Brunschwig H, Levi L, Ben-David E, Williams RW, Yakir B, Shifman S. Fine-scale maps of recombination rates and hotspots in the mouse genome. Genetics. 2012;191(3):757–64.

113. Booker TR, Ness RW, Keightley PD. The Recombination Landscape in Wild House Mice Inferred Using Population Genomic Data. Genetics. 2017;207(1):297–309.

114. Corbett-Detig RB, Hartl DL, Sackton TB. Natural selection constrains neutral diversity across a wide range of species. PLoS Biol. 2015;13(4):e1002112.

115. Hinch R, Donnelly P, Hinch AG. Meiotic DNA breaks drive multifaceted mutagenesis in the human germ line. Science. 2023;382(6674):eadh2531.

116. Spencer CC. Human polymorphism around recombination hotspots. Biochem Soc Trans. 2006;34(Pt 4):535–6.

117. Clement Y, Arndt PF. Meiotic recombination strongly influences GC-content evolution in short regions in the mouse genome. Mol Biol Evol. 2013;30(12):2612–8.

118. Duret L, Galtier N. Biased gene conversion and the evolution of mammalian genomic landscapes. Annu Rev Genomics Hum Genet. 2009;10:285–311.

119. Crête-Lafrenière A, Weir LK, Bernatchez L. Framing the Salmonidae family phylogenetic portrait: a more complete picture from increased taxon sampling. PLoS One. 2012;7(10):e46662.

120. Zelkowski M, Olson MA, Wang M, Pawlowski W. Diversity and Determinants of Meiotic Recombination Landscapes. Trends Genet. 2019;35(5):359–70.

121. Ahlawat S, De S, Sharma P, Sharma R, Arora R, Kataria RS, et al. Evolutionary dynamics of meiotic recombination hotspots regulator PRDM9 in bovids. Mol Genet Genomics. 2017;292(1):117–31.

122. Stevison LS, Woerner AE, Kidd JM, Kelley JL, Veeramah KR, McManus KF, et al. The Time Scale of Recombination Rate Evolution in Great Apes. Mol Biol Evol. 2016;33(4):928–45.

123. Spence JP, Song YS. Inference and analysis of population-specific fine-scale recombination maps across 26 diverse human populations. Sci Adv. 2019;5(10):eaaw9206.

124. Fumasoni I, Meani N, Rambaldi D, Scafetta G, Alcalay M, Ciccarelli FD. Family expansion and gene rearrangements contributed to the functional specialization of PRDM genes in vertebrates. BMC Evol Biol. 2007;7:187.

125. Ma L, O’Connell JR, VanRaden PM, Shen B, Padhi A, Sun C, et al. Cattle Sex-Specific Recombination and Genetic Control from a Large Pedigree Analysis. PLoS Genet. 2015;11(11):e1005387.

126. Baker CL, Petkova P, Walker M, Flachs P, Mihola O, Trachtulec Z, et al. Multimer Formation Explains Allelic Suppression of PRDM9 Recombination Hotspots. PLoS Genet. 2015;11(9):e1005512.

127. Flachs P, Mihola O, Simecek P, Gregorova S, Schimenti JC, Matsui Y, et al. Interallelic and intergenic incompatibilities of the prdm9 (hst1) gene in mouse hybrid sterility. PLoS Genet. 2012;8(11):e1003044.

128. Huang T, Yuan S, Gao L, Li M, Yu X, Zhang J, et al. The histone modification reader ZCWPW1 links histone methylation to PRDM9-induced double strand break repair. eLife. 2020;9.

129. Mahgoub M, Paiano J, Bruno M, Wu W, Pathuri S, Zhang X, et al. Dual histone methyl reader ZCWPW1 facilitates repair of meiotic double strand breaks in male mice. eLife. 2020;9.

130. Wells D, Bitoun E, Moralli D, Zhang G, Hinch A, Jankowska J, et al. ZCWPW1 is recruited to recombination hotspots by PRDM9, and is essential for meiotic double strand break repair. eLife. 2020;9.

131. Joseph J, Prentout D, Laverré A, Tricou T, Duret L. High prevalence of Prdm9-independent recombination hotspots in placental mammals. bioRxiv. 2023:2023.11.17.567540.

