## Supporting information for "PRDM9 drives the location and rapid evolution of recombination hotspots in salmonids"

Raynaud et al. (26/02/2024)


1. **Supplementary Methods** 1
2. **Supplementary Analysis:** Prediction of CGI-associated TSSs in salmonids 16
3. **Supplementary Figures** 23
4. **Supplementary Tables** 44
5. **Supplementary references** 62


#### 1. S1 Methods

##### Phylogenetic analysis of PRDM9 paralogs across Salmonids

We investigated the presence of a complete PRDM9 protein in twelve species from the three salmonid subfamilies (Coregoninae, Thymallinae, Salmoninae). We deduced a reference protein sequence for the three canonical domains of PRDM9 in the coho salmon: KRAB (encoded by 2 exons), SSXRD (1 exon) and SET (3 exons). This was obtained from a nearly full-length CDS annotated in the RefSeq database (XP_020359152.1), complemented in its 3' end using a cDNA identified in a brain RNAseq dataset sequenced with PacBio long reads (SRR10185924.264665.1). The nucleotide sequence of the 6 exons was retrieved as the best tblastn (1) hit of the protein sequence against the *O. kisutch* reference genome. This reference sequence was subsequently used to identify PRDM9 homologs in the whole genome assembly of the lake whitefish (*Coregonus clupeaformis*, assembly ASM2061545v1, RefSeq accession number GCF_020615455.1), the European grayling (*Thymallus thymallus*, fThyThy.pri.20220222, GenBank accession number GCA_023634145.1), the huchen (*Hucho hucho*, ASM331708v1, GCA_003317085.1), the coho salmon (*O. kisutch*, Okis_V2, GCF_002021735.2), the rainbow trout (*O. mykiss*, Omyk_1.0, GCF_002163495.1), the chinook salmon (*O. tschawytscha*, Otsh_v2.0, GCF_018296145.1), the chum salmon (*O. keta*, Oket_V2, GCF_023373465.1), the red salmon (*O. nerka*, Oner_1.1, GCF_006149115.2), the pink salmon (*O. gorbuscha*, OgorEven_v1.0, GCF_021184085.1), the Atlantic salmon (*S. salar*, Ssal_v3.1, GCF_905237065.1), the brown trout (*S. trutta*, fSalTru1.2, GCA_901001165.2), the lake trout (*Salvelinus namaycush*, GCF_016432855.1), as well as a closely related outgroup, the northern pike (*Esox lucius*, fEsoLuc1.pri, GCF_011004845, Esocidae). The six exons were independently blasted to the 13 assemblies with both blastn and tblastn allowing multiple hits. We filtered out copies containing only one of the six exons. We used annotated *Prdm* family genes from the human (GRCh38.p14, GCF_000001405.40) and mouse (GRCm39, GCF_000001635.27) Ensembl genomes to remove non-*Prdm9* copies. Retained exons were aligned separately using Macse (v2.06)(2), a program of coding sequence alignment simultaneously accounting for the nucleotide and amino-acid levels, thus potentially allowing for the inclusion of frameshifts and stop codons. The alignments were manually examined and edited before concatenating exons of the same copy using Amas concat (3). Salmonids have undergone two recent whole duplication (WGD) events that occurred in the common ancestor of teleosts (Ts3R, c.a. 320 Mya) and salmonids (Ss4R, c.a.

90 Mya) respectively (4-6). Consequently, their chromosomes appeared as duplicate pairs referred to as ohnolog chromosomes, derived from the same ancestral chromosome. We therefore retrieved the chromosomal locations of the retained copies in order to trace the evolutionary history of Prdm9 duplications. A maximum-likelihood phylogeny of the three canonical domains was built using IQ-TREE (7) based on amino-acid alignments, using ultrafast bootstrap with 1000 replicates. In order to identify functional *Prdm9* copies with sequence orthology to the 10 exons found in human and mouse (8), we finally predicted the gene structure of each copy surrounded by its 10kb flanking regions using Genewise (v2.4.1)(9). We selected representative paralogous sequences across the obtained PRDM9 phylogenetic tree (accession number: XP_036826454.1, XP_055786392.1, XP_014035317.2, XP_055791730.1, XP_036838644.1) to perform a sequence similarity-based annotation of the copies in each species.

#### Analysis of PRDM9 ZF diversity in rainbow trout and Atlantic salmon

##### *Samples used for Prdm9α Zinc Finger Array genotyping*

We characterized the allelic diversity of the ZF array domain of *Prdm9*α in two species: *S. salar* and *O. mykiss*. We focused on *Prdm9α*, as *Prdm9β* orthologs in teleost fish were previously shown to lack KRAB and SSXRD domains, to have a slowly evolving ZF array and to carry a presumably inactive SET domain (10). We used wild Atlantic salmon (*S. salar*) samples from Normandy (France) that were biopsied with caudal fin clips during routine monitoring and stored in ethanol at -20°C. Samples were kindly provided by Guillaume Evanno (INRAE). We also analysed rainbow trout (*O. mykiss*) samples of unknown genetic origin that came from a population acquired by the INRAE in the 1990s and selectively bred for fall spawning (ID: INRA-AUT) at an INRAE experimental fish farm (PEIMA). Trout samples were generously supplied by Jean-Jacques Lareyre (INRAE). Genomic DNA was extracted from fin clips using the Qiagen DNeasy Blood & Tissue kit following the manufacturer's instructions. DNA concentration and purity were measured with a Nanodrop-1000 Spectrophotometer (Thermo Fisher Scientific), and quality was assessed via agarose gel electrophoresis. 10 ng/µl working dilutions were prepared and stored at -20 °C.

##### *Amplification and sequencing of Prdm9α Zinc Finger Array*

We inferred the expression levels of multiple *Prdm9α* paralogs in immature testes from both genera, *Salmo* and *Oncorhynchus*, using publicly available RNA-seq data from SRA. Specifically, we analyzed data from two samples in *S. salar* (SRR1422872 and SRR9593306) and two samples in *O. kisutch* (SRR8177981 and SRR2157188). Our analysis revealed high expression of two distinct *Prdm9α* paralogs in both genera, which were previously identified in the phylogenetic analysis. Namely, we then sequenced *Prdm9α* paralog *α1.a.2* (full length, chromosome 5, n=26) and *α2.2* (partial, chromosome 17, n=20) in *S. salar*, and *Prdm9α* paralog *α1.a.1* (full length, chromosome 31, n=23) and *α2.2* (partial, chromosome 7, n=20) in *O. mykiss*. The complete list of samples used in this study is available in **S1 Table**.

Primers were designed using NCBI Primer Blast, ensuring specificity against the reference assemblies (*S.salar* ICSASG_v2 and *O. mykiss* USDA_OmykA_1.1). They targeted the ZF sequence encoded in the last exon of the gene, framed by the flanking arms of the array, avoiding any specificity of the paralogous loci. The primers were synthetized by Eurofins Genomics (Ebersberg, Germany). For primers sequences refer to **S2 Table**.

PCR reactions were carried out in 25 µl volume containing 1X Phusion HF buffer, 200 µM dNTPs, 0.5 µM forward primer, 0.5 µM reverse primer, 3% DMSO, 2.5-10 ng of template and 0.5 units of Phusion Polymerase (NEB). Cycling conditions were: initial denaturation at 98 °C for 2 min followed by 35 cycles of 98 °C 10 s, 66-70 °C 30 s, 72 °C 1:30 min, a final elongation at 72 °C 3 min, and hold at 10 °C in a C1000 Cycler (Bio-Rad). PCR products were inspected on a 1% agarose gel stained with 0.5 µg/ml ethidium bromide. Samples were re-amplified if the amplification was not efficient or in case smearing was observed. Samples showing a single size amplicon, were purified using the NucleoSpin Gel and PCR clean-up kit (Machery-Nagel) and Sanger sequenced in 5' and 3' direction. Samples showing two different length alleles were separated by electrophoresis, each single band was purified and cloned using the TOPO Blunt cloning kit (Invitrogen). The ligated vector was then transformed into OneShotTop10 chemically competent cells (Invitrogen), following the manufacturer's instructions. At least 4 clones per amplicon were purified using the QIAprep Spin Miniprep Kit (Qiagen) and they were Sanger sequenced either using the same primers used to sequence the PCR products or, plasmid specific primers M13 forward and reverse. When Sanger sequencing revealed heterozygosity in the chromatograms from single-sized PCR product, the remaining amplicon was cloned and sequenced as just described for heterozygous samples. All the sequencing was conducted by Azenta-GeneWiz (Leipzig, Germany).

##### *Allelic diversity analysis*

The sequencing results were processed as follows. Forward and reverse reads were *de-novo* assembled to generate contigs and each contig was aligned to the reference ZF array using SnapGene software (versions 5.1.4.1 - 5.2.3). Each contig was translated either using SnapGene, or ExPaSy Translate tool (https://web.expasy.org/translate/) and the single letter amino acid sequence corresponding to each ZF array was recorded, this allowed us to categorize individual PRDM9α alleles. All ZF arrays were annotated such that they matched a C2H2 ZF motif: X7-CXXC-X12-HXXXH, where X is any amino acid, C indicates the cysteines and H the histidine residues. We reported a new allele whenever a single variation in the array at the level of the amino acid sequence was found. DNA sequences for each allele were aligned to create a consensus sequence.

We then determined the proportion of amino acid diversity at DNA-binding residues of the ZF array (positions -1, 2, 3, and 6 of the α-helix) as in (10, 11). The ZF arrays were adapted to match the following motif for the alignment: X2-CXXC-X12-HXXXH-X5. Briefly, we calculated the amino acid diversity as a function of amino acid position in the ZFs array. A single ZF array was analyzed for each paralog by pooling all ZF types identified, except for the partial ZFs (of type X2-CXXC-X12-H-X9). The diversity plots were generated by plotting the heterozygosity values reported at each site of the ZF array. We calculated the proportion of the total amino acid diversity (r) at DNA-binding sites as the sum of heterozygosity at DNA-binding residues over the sum of heterozygosity at all 28 residues of the array.

##### *Prediction of DNA-binding motifs of different Prdm9α alleles*

The DNA-binding sequences of each *Prdm9α* allele identified were predicted *in-silico* using the polynomial SVM method from Princeton C2H2 ZF binding site predictor (http://zf.princeton.edu/) as described (12, 13). Notably, for alleles with a partial last ZF in the array (where the last histidine was absent), predictions were performed excluding this incomplete ZF.

#### Identification of DSB hotspots using ChIP-Seq in the rainbow trout

##### *Samples used for ChIP-seq experiments*

We investigated the genome-wide distribution of DMC1-bound ssDNA in *O. mykiss* testes by chromatin-immunoprecipitation (ChIP) followed by ssDNA enrichment (DMC1-Single Strand DNA Sequencing, DMC1-SSDS). We chose three rainbow trout individuals from the pool of samples previously used for characterizing PRDM9 ZF diversity. Experiments were performed in accordance with the CNRS guidelines for animal welfare and ethical authorization n° APAFIS#13616-2018021315504139 v5 issued by the local committee for ethical animal experimentation and the French ministries of research and agriculture. The rainbow trout samples belong to the cohort previously described in section "*Samples used for PRDM9 Zinc Finger Array genotyping".* We assessed the gonadal stage of the samples, selecting those at an optimal meiotic stage for further analysis. Male gonads were excised, with the majority immediately snap-frozen in liquid nitrogen and stored at -80°C for subsequent processing. A smaller portion was fixed in Bouin's solution, dehydrated through increasingly concentrated ethanol solutions and finally imbedded in paraplast. 7 µm-thick tissue sections were stained with hematoxylin/eosin staining solutions (Diapath) and mounted using the afcolene mounting medium. The stage of the gonadal maturation was determined from macroscopic and histological observations of the gonads and gonadal sections, respectively, according to (14). In testes at stage I, only undifferentiated A spermatogonia are observed. In stage II testes, testicular lobules become organized and many B spermatogonia are present in addition to A spermatogonia. Testes at stages III and IV increase in size and contain an increasing number of primary spermatocytes including leptotene, zygotene, and pachytene spermatocytes. Few spermatids appear concomitantly. Testes at stages V to VI continue to grow and become whitish due the rapid increasing accumulation of spermatids and spermatozoa, respectively. At stage VII, the collector tube becomes dilated and fills will sperm. At stage VIII, testes reach their maximum size and only spermatozoa are observed in the lumen of the testicular lobules (spermiation). Since DMC1 binds to chromatin during early stages of the meiotic prophase I, we used testes at stages III and IV to perform the chromatin immunoprecipitation experiments. We thus selected three individuals with different *Prdm9* genotypes (TAC-1: *Prdm9*$^{1/5}$, TAC-3: *Prdm9*$^{2/6}$ and RT-52: *Prdm9*$^{1/2}$), enabling the comparison of DSB hotspots between individuals sharing or not sharing a *Prdm9* allele.

##### *Antibodies*

Antibodies against DMC1 were from Yukiko Imai (NIG). Briefly, the antibodies were raised by immunizing two rabbits and one guinea pig with a His-tagged recombinant protein of zebrafish Dmc1, corresponding to amino-acid residues 7-220 (UniProt ID: B3DIR0). Dmc1 polyclonal serum was then affinity-purified by using CNBr Activated Sepharose™ 4B (Cytiva) conjugated with the Dmc1 recombinant protein. The obtained rabbit1, rabbit2 and guinea pig anti-zbDmc1, were tested by western blot on protein extracts from trout testes (result not shown) to select the most specific one. For DMC1 ChIP, we used either rabbit1 or guinea pig anti-zbDmc1, for H3K4me3 ChIP we used rabbit anti-H3K4me3 (Abcam, ab8580) and for H3K36me3 ChIP we used rabbit anti-H3K36me3 (Diagenode, Premium, C15410192).

##### *Chromatin immunoprecipitation*

We conducted crosslink ChIP experiments for H3K4me3 and H3K36me3 using protocols described (15, 16), with some adjustments. Briefly, a portion of frozen tissue (20-25 mg) was immediately immersed in PBS, 1% formaldehyde for 10 min at room temperature. After quenching the unbound PFA by addition of glycine to a final concentration of 250mM, the tissue was transferred in PBS for homogenization with a 2 ml glass dounce and the cell suspension was filtered with a 40µm cell strainer (Falcon). Cells were washed twice in buffer A (10mM Tris-HCl pH 8.0, 10mM KCl, 0.25% Triton X-100, 1mM EDTA, 0.5mM EGTA, 1x cOmplete protease inhibitor cocktail EDTA-free (Roche)) for 5 min on ice. After spinning, cells were washed once in buffer B (10mM Tris pH 8.0, 200mM NaCl, 1mM EDTA, 0.5mM EGTA, 1x cOmplete) for 10 min on ice. After incubation and spinning, cells were lysed in 1% SDS, 10mM EDTA, 50mM Tris-HCl pH 8.0, 1x cOmplete for 30 min at 4°C on a rotating wheel. Sonication was performed on a Bioruptor UCD-300 sonicator (Diagenode) with the following conditions: 30 sec ON, 30 sec OFF, High power, 4x10 Cycles. Chromatin was diluted ten folds in IP dilution buffer: 5mM Tris-HCl pH 8.0, 140mM NaCl, 0.5% Triton X-100, 0.05% sodium deoxycholate, 0.5mM EGTA and one ml of chromatin was pre-cleared with 20 µl of Dynabeads Protein A (Invitrogen) for 4 h at 4°C on a rotating wheel. In parallel, 20 µl of Dynabeads Protein A were incubated with 3 µg of the appropriate antibody for 5 h at 4°C on a rotating wheel. After incubation with antibodies, the beads were washed twice in PBS, 0.05% Tween 20, 0.1mM DTT for 5 min at 4°C on a rotating wheel, and then once in IP dilution buffer for 10 min at 4°C on a rotating wheel. After the pre-clear, one ml of chromatin was incubated with the beads-antibody complex overnight at 4°C on a rotating wheel. IPs were washed once on each of the following buffers: Wash 1 (10mM Tris-HCl pH 8, 150mM KCl, 0.50% NP40, 1 mM EDTA), Wash 2 (10mM Tris-HCl pH 8, 100mM NaCl, 0.10% sodium deoxycholate, 0.50% Triton X-100), Wash 3a (10mM Tris-HCl pH 8, 400mM NaCl, 0,10% sodium deoxycholate, 0.50% Triton X-100), Wash 3b (10mM Tris-HCl pH 8, 500mM NaCl, 0,10% sodium deoxycholate, 0.50% Triton X-100), Wash 4 (10mM Tris-HCl pH 8, 250mM LiCl, 0,50% sodium deoxycholate, 0.50% NP40, 1 mM EDTA), TE (10mM Tris-HCl, 1mM EDTA). DNA was eluted (and cross-linking reversed) by incubation at 65°C overnight in 50mM Tris-HCl pH 8, 1% SDS, 1mM EDTA. After treatments with RNAse A and proteinase K, DNA was purified with MinElute columns (Qiagen, 28004).

DMC1 ChIP experiments were executed following the methods previously outlined in (17) and (18). The chromatin was prepared from 150-200 mg of testes. Frozen tissue was fixated in PBS, 1% formaldehyde for 10 min at room temperature. Glycine was added to a final concentration of 125mM to quench unreacted PFA and the tissue was passed through a 7 ml dounce for homogenization. Single cell suspension was filtered passing it through a 70µm cell strainer (Falcon). Cells were washed once in PBS and after centrifugation, cells were resuspended in buffer L1 (10mM Tris-HCl pH 8.0, 10mM EDTA, 0.5mM EGTA, 0.25% Triton X-100, 1x cOmplete protease inhibitor cocktail EDTA-free (Roche)). After spinning, cells were resuspended in buffer L2 (10mM Tris pH 8.0, 200mM NaCl, 1mM EDTA, 0.5mM EGTA, 1x cOmplete). After centrifugation cells were resuspended in buffer L3 (1% SDS, 10mM EDTA, 50mM Tris-HCl pH 8.0, 1x cOmplete) and immediately sonicated on a Bioruptor UCD-300 sonicator with the following conditions: 15 sec ON, 45 sec OFF, High power, 15 Cycles. Sheared chromatin was diluted by adding one volume of ChIP dilution buffer: 16.7mM Tris-HCl pH 8.0, 167mM NaCl, 1.1% Triton X-100, 0.01% SDS, 1.2mM EDTA. The chromatin was dialyzed for 5 hours at 4°C using constant rotation prior incubation with antibodies. The chromatin was incubated overnight with 24µg of antibody at 4°C on a rotating wheel. 150 µl of Dynabeads Protein A were incubated with 3 ml of ChIP for 2 h at 4°C on a rotating wheel. IPs were washed once on each of the following buffers: Wash 1 (20mM Tris-HCl pH 8, 150mM NaCl, 1% Triton X-100, 0.1% SDS, 2mM EDTA), Wash 2 (20mM Tris-HCl pH 8, 500mM NaCl, 1% Triton X-100, 0.1% SDS, 2mM EDTA), Wash 3 (10mM Tris-HCl pH 8, 250mM LiCl, 1%

Deoxycholic acid, 1% IGEPAL, 1mM EDTA). Two final washes in TE (10mM Tris-HCl, 1mM EDTA) and DNA elution by incubation at 65°C for 30 min in 100mM $NaHCO_3$, 1% SDS. Beads were discarded and cross-linking was reversed by incubating at 65°C overnight in 200mM NaCl,100mM $NaHCO_3$, 1% SDS. After treatment with proteinase K, DNA was purified with MinElute columns (Qiagen, 28004).

##### *Library preparation and sequencing*

For histone modifications ChIP, standard library construction was performed according to the NEBNext Ultra II protocol for Illumina (NEB, E7645S-E7103S), with minor adjustments. All clean-up steps in the protocol were achieved though MinElute columns (Qiagen, 28004). For size selection, we performed agarose gel electrophoresis and DNA fragments within the range of 200 to 400 bp were excised from the gel and the DNA purified using MinElute columns (Qiagen, 28604). For DMC1 ChIP, library construction was performed following the Illumina TruSeq protocol (Illumina, IP-202-9001DOC), with the introduction of an extra step of kinetic enrichment as previously described (17, 18). The NovaSeq6000 platform (Illumina) with S4 flow cells was employed for sequencing the libraries, and all the sequencing procedures were carried out at Novogene Europe (Cambridge, United Kingdom).

##### *Computational Data Analysis*

For all ChIP-seq experiments, paired-end reads were mapped to the USDA_OmykA_1.1 assembly. We analyzed histone modifications through the nf-core/chipseq v1.2.1 pipeline (19), accessible at https://github.com/nf-core/chipseq. The pipeline was executed using *Nextflow v20.10.0*. The sequencing reads were aligned to the reference genome with *BWA v0.7.17-r1188*. The pipeline filtered by default the mapped reads to retain only the non-duplicated and high-quality uniquely mapping reads. For both histone marks, we normalized the signal based on the total read coverage and by subtracting the input using *bamCompare* function (*normalizeUsingRPKM*) of *deepTools* (20). Peak calling was performed using *MACS2 v2.2.7.1* (parameters: *--pvalue=1e-5 --narrow_peak --read_length=150 --macs_gsize 2.07E9*) on both replicates, and an input for each sample was provided. The peaks found within 1000 bp of one another were combined using *merge* function of *Bedtools* (21), and the mean of their scores was reported (parameters: -d 1000 -c 5 -o mean). We tested the enrichment of the two histone marks at RefSeq annotated genes to check the quality of the experiments (*i.e.* we validated H3K4me3 enrichment at TSS and H3K36me3 at gene body, data not shown). We did that through the *computeMatrix* function of *deepTools,* using *scale-regions* mode. We assessed the enrichment of the histone marks at DMC1 peaks by using *deepTools computeMatrix*, *plotProfile* and *plotHeatmap* to compute and display average profiles and heatmaps (**S9 Fig**) (parameters*: reference-point --referencePoint center -b 5000 -a 5000 --skipZeros --missingDataAsZero --scale 1*). Using the same parameters, we assessed the enrichment of H3K36me3 signal at H3K4me3 peaks in brain using the bed files produced in the context of the Aqua-FAANG project (https://www.aqua-faang.eu/). The peaks were processed by *Bedtools merge* and *Bedtools interstect* to allow meaningful comparison with our H3K4me3 data. Details regarding the overlapping strategies applied can be found in the "*Determination of Overlapping Peaks*" section. Subsequently, the bed files were subjected to analysis using *deeptools computeMatrix* and *deeptools plotHeatmap* to compute and visualize heatmaps.

The analysis of DMC1 ChIP-seq was performed as described (18), with some implementations (22). We used the *hotSSDS* pipeline (version 1.0), which is an adaptation of the original *SSDS* pipeline (https://github.com/kevbrick/SSDSnextflowPipeline) and the *SSDS call peaks* pipeline (https://github.com/kevbrick/callSSDSpeaks) created by Kevin Brick (NIH). The *hotSSDS* pipeline was customized and modified by Pauline Auffret (IFREMER) and Julie Clément (IHPE), and their version can be found at https://github.com/jajclement/hotSSDS. The pipeline was executed using *Nextflow 21.10.0*. Reads were mapped using the modified BWA algorithm (*BWA Right Align*), initially developed by (18) to align and recover ssDNA fragments, then improved (17). Aligned reads were filtered in order to retain only the non-duplicated and high-quality uniquely mapped reads. The signal was normalized by the library size, and then by the total type 1 ssDNA fragments. The peak calling was performed using *MACS2 v2.2.7.1* with relaxed conditions (parameters: *--pvalue=1e-2 --bw=1000 --nomodel --slocal=5000 --extsize=800*) on each of the two replicates, while an input control was provided. Peak calling was additionally performed on the pooled dataset, and on pseudo-replicates that were artificially generated by randomly subsampling half the reads twice from each replicate (self pseudo-reps), or from the pooled dataset (pooled pseudo-reps). Then Irreproducible Discovery Rate (*IDR*) analysis was performed as described (23) to allow the identification of highly reproducible peaks between the true replicates (with a threshold of 0.05), between the pooled pseudo-replicates (with a threshold of 0.01), or between self pseudo-replicates (with a threshold of 0.05), to assess the self-reproducibility. Final peak sets were created by picking up the top peaks from the dataset of origin, meaning all peaks having an *IDR* threshold inferior to 0.01 for pooled dataset, as recommended by the authors. The IDR peaks from the pooled dataset were processed using the script *normalizeStrengthByAdjacentRegions.pl* developed by Brick. The script considers the distribution of ssDNA fragments to recenter the peaks by the median of the forward/reverse fragments distribution and outputs recentered and normalized peaks.

The final peaks were used to check the distribution of ssDNA type 1 signal at the DSB hotspots using *deepTools computeMatrix* and *deepTools plotHeatmap* (parameters: *reference-point --referencePoint center -b 2500 -a 2500 --skipZeros --missingDataAsZero --scale 1 --maxThreshold 15* for TAC-1 and *--maxThreshold 10* for TAC-3). Additionally, we computed the relative position of each DSB along its respective chromosome by utilizing the R package *dplyr* (**S9 Fig**). As replicates were validated to be reproducible, we merged the normalized read distribution from each replicate (*i.e.* 2 bigwig files per sample) for further analyses by the *UCSC Genome Browser*'s utility.

##### *Correlation between read enrichments of histone marks*

To assess the correlation in signal distribution from histone ChIP between samples, we used the *multiBigwigSummary* function from *deepTools*. This tool computes summary statistics for multiple bigwig files, more specifically on signal intensity measured across specified genomic intervals. We thus calculated the correlation between H3K4me3 signal and H3K36me3 signal at RT-52 DMC1 peaks, the H3K4me3 signal in TAC-1 and TAC-3 samples at the RT-52 DSB hotspots and the correlation between H3K36me3 signal in TAC-1 and TAC-3 samples at the same regions (using read distribution bigwig files obtained for merged replicates). The statistical significance of the results was evaluated using the chi-square test, and scatterplots were generated using GraphPad Prism software.

##### *Analysis of DMC1 ChIP-seq signal at genomic features*

To investigate the DSB activity at various genomic features, we used the distribution of DMC1-SSDS read enrichment as a metric. The genomic features of interest were initially identified on the assembly Omyk_1.0, as described in section "*Recombination at genomic features*" below. We used *NCBI remap* tool to perform the liftover of the genomic regions on the assembly USDA_OmykA_1.1 (parameters: *--mode asm-asm --from GCF_002163495.1 --dest GCF_013265735.2 --annotation* <*bed file*> *--annot_out* <*bed file*>). Subsequently, we applied the functions *distanceToNearest* and *subsetByOverlaps* from the R package *GenomicRanges* (24), as detailed in the aforementioned section. The bedgraph files obtained from the DMC1-SSDS corresponding to read distribution for merged replicates were used to calculate the averaged read distribution within each genomic feature.

##### *Determination of overlapping peaks*

We used the *intersect* function of Bedtools to assess the overlaps between different features by the default options. For overlaps purposes, the central 400 bp of each DSB hotspot peak was used, as previously described (25). When comparing DMC1 peaks with H3K4me3 marks and H3K36me3 marks, we used the central 400 bp of the hotspots and the full-length positions of the histone marks. To determine the intersection with the transcription start/end sites, we defined the TSS and TES regions as, respectively the 5' and the 3' end of the transcripts from the RefSeq annotated genes ±1 kb. To identify the overlapping intervals between replicates and between samples when the IDR methodology was not applied (*i.e.* for H3K4me3 and H3K36me3 peaks), we used the tool with more stringent criteria (parameters: *-f 0.25 -r -u*) to retrieve reproducible peaks.

##### *Analysis of DSB hotspot annotation*

To annotate the DSBs in relation to the RefSeq genome annotation, we employed the *annotatePeaks.pl* function from the *HOMER* suite, utilizing the *-gtf* option. For visualizing the results, we utilized a script adapted from the *hotSSDS-extra* pipeline (22) for further processing the results generated by the *hotSSDS* pipeline. This pipeline is designed to automate the computation and visualization of general statistical data related to the SSDS signal, and can be found at (https://github.com/jajclement/hotSSDS-extra). Specifically, we used the script named *plot_homer_annotatepeaks.r*, which was integrated in the *nf-core chipseq v1.2.1* pipeline, as described (19). This script takes the results of peak annotation to generate visual representations (**Fig 3B**).

#### Reconstruction of population-based recombination landscapes in three salmonids (indirect estimation of recombination rates)

##### *Whole genome data*

We collected high coverage whole-genome resequencing data from five natural populations of three salmon species from the SRA database to reconstruct population-based recombination landscapes, with approximately 20 samples per population as recommended (26). We retrieved 20 samples from the Southern British Columbia population of the coho salmon, *O. kisutch* (27), 22 rainbow trout (*O. mykiss)* samples from North East America (28), and 60 genomes from three populations of Atlantic salmon (*S. salar)* belonging the two major lineages in North America and Europe (29): 20 from the Gaspesie Peninsula in Canada (referred to as the GP population), 20 from the North Sea (NS population) and 20 from the Barents Sea in Norway (BS population). Sample accession number and location is detailed in **S4 Table** and **S1 Fig**.

##### *Variant Calling*

Variant calling for *O. kisutch* was performed by (27). We followed the same methodology for variant calling and genotyping in *O. mykiss* and *S. salar*, using the GATK best-practice pipeline (> v3.8-0, see detailed versions of the programmes used in **S5 Table**) (30, 31). First, we aligned individual paired-end reads against their reference genome (assembly and RefSeq accession number: *O. kisutch*: Okis_v1, GCF_002021735.1; *O. mykiss*: Omyk_1.0, GCF_002163495.1; *S. salar*: Ssal_v3.1, GCF_905237065.1) using bwa-mem (v0.7.17) (32), yielding an average depth coverage per sample of 29.54x, 24.87x and 9.97x for *O. kisutch*, *O. mykiss* and *S. salar*, respectively (**S6 Table**). Among the multiple primary alignments generated per query sequences, we flagged shorter split hits as secondary with the *-M* option for Picard compatibility. Possible PCR duplicates were then marked with the Picard MarkDuplicates program (> v2.18.29, validation stringency parameter was set to lenient). All reads were assigned to a new read-group ID using Picard AddOrReplaceReadGroups. We called variants for each individual of *O. mykiss* and *S. salar* using HaplotypeCaller with allele-specific annotations, generating GVCF files (options *-G StandardAnnotation, -G AS_StandardAnnotation, -G StandardHCAnnotation*). After creating a GenomicsDB workspace (GenomicsDBImport), joint genotyping (GenotypeGVCFs, default settings) was completed by pooling all individuals for each species. A total of 9,590,270, 39,601,311 and 27,061,466 SNPs were called for *O. kisutch* (27), *O. mykiss* and *S. salar*, respectively.

Following genotyping, we removed variants within 5 bp of an indel with Bcftools filter (v 1.9; *-g 5*). We filtered low-quality SNPs with Vcftools (> v 0.1.16) (33). We excluded indels (*--remove-indel*), variants with more than 2 alleles (*--max-alleles 2, --min-alleles 2*), genotypes with quality scores < 20 (--minGP 20), and SNPs with >10% of missing values (*--max-missing 0.9*). For *S. salar*, the missingness threshold was set to 50% because of a higher rate of missing genotypes in this dataset. To control for poorly sequenced regions or duplicated loci, we kept only sites with a mean depth coverage falling within the 5–95% quantiles of the species' distribution (*O. kisutch*: 13x-33x; *O. mykiss*: 7x-23x; *S. salar*: 2x-9x). We applied a Hardy-Weinberg test with a *p*-value threshold of 0.01 (*--hwe 0.01*), expecting to filter excess of heterozygosity due to hidden paralogy in particular within residual tetrasomic regions. We removed singletons by applying a MAC (minor allele count) filter with Vcftools (*--mac 2*). For *S. salar*, the Hardy-Weinberg and MAC filters were applied separately for each of the three populations. The filtering steps retrieved 7,205,269, 16,079,097 and 5,575,430 SNPs for *O. kisutch*, *O. mykiss* and *S. salar*, respectively (**S6 Table**).

##### *Variant phasing and orientation*

We used the read-based phasing approach in WhatsHap (> v0.18) (34) to identify phase blocks from paired-end reads overlapping neighboring individual heterozygous positions. Prephasing statistics calculated using whatshap *stats* showed that 73.45% and 76.98% of the variants were physically phased for *O. kisutch* and *O. mykiss*. Due to the lower SNPs density and higher genotype missingness, the *S. salar* dataset was prephased for only 7.32% of its variants. Prephased blocks were then phased chromosome-wide using the statistical phasing approach in SHAPEIT4 (> v4.2.1) (35), default settings: phase-set error rate *--use-PS 0.0001* and MCMC iteration scheme *5b,1p,1b,1p,1b,1p,5m*), assuming a constant recombination rate of 3 cM/Mb (representative of average recombination rates in teleosts (36)) and using the effective population size estimated from the nucleotide diversity of each chromosome, which was calculated in 100-kb windows using Vcftools.

Ancestral allelic states were inferred for the set of variants of each species. Three outgroups were used for each species: *O. kisutch* variants were orientated using the reference genome of *O. mykiss*, the sockeye salmon *O. nerka* (assembly Oner_1.1, RefSeq accession number GCF_006149115.1) and the chinook salmon *O. tshawytscha* (Otsh_v1.0, GCF_002872995.1), *O. mykiss* with *O. kisutch*, *O. nerka* and *O. tschawytscha*, and *S. salar* with *O. mykiss*, the brown trout *S. trutta* (fSalTru1.2, GCA_901001165.2), and the arctic char *Salvelinus alpinus* (ASM291031v2, GCF_002910315.1). We retrieved 100bp-flanking sequences of each SNPs in the ingroup species with the *getfasta* function of Bedtools and used blastn to find their ortholog positions on the reference genome of each outgroup species, retaining only the best hits (*-outfmt 6, -max_target_seqs 1, -max_hsps 1*). The corresponding SNPs positions in the query sequences were retrieved using a custom rust script. Ancestral state probabilities were then inferred with the maximum likelihood method implemented in est-sfs (v2.04) (37), using the ingroup allele frequencies and the allelic states of the outgroups. Phylogenetic relationship between ingroup and outgroup species were taken into account (38) and the mean sequence identity scores computed with blastn: (((*O. kisutch, O. tshawytscha*), *O. nerka*), *O. mykiss*); (((*O. mykiss, O. tshawytscha*), *O. nerka*), *O. kisutch*); (((*S. salar, S. trutta*), *S. alpinus*), *O. mykiss*) (see **S10 Table**). Est-sfs was run using the Kimura-2-parameter substitution model.

##### *Estimation of linkage disequilibrium-based recombination rates*

We estimated population recombination rates $\rho$ ($\rho=4N_e r$, with $N_e$ the effective population size and *r* the recombination rate in M/bp) with LDhelmet (v1.19) (39) for the five populations of *O. kisutch*, *O. mykiss*, and S. salar (*i.e.* GP, BS and NS). LDhemet infers the $\rho$ parameter between pairs of SNPs with a reversible-jump Markov Chain Monte Carlo algorithm. SNPs data were converted into fasta sequences for each individual haplotype using the *vcf2fasta* function of vcflib (https://github.com/vcflib/vcflib), and into the position and SNPs input format with the *--ldhelmet* option of VCFtools. Ancestral allelic states were provided using the probability (P) of the major allele being ancestral computed by est-sfs, and 1-P to the minor allele. LDhelmet was run five times independently for each population. For each chromosome, the haplotype configuration files were created with the find_conf function using the recommended window size of 50 SNPs. The likelihood look-up tables were created once for the five runs with the table_gen function using the recommended grid for the population recombination rate (*ρ*/pb) (*i.e. ρ* from 0 to 10 by increments of 0.1, then from 10 to 100 by increments of 1), and with the Watterson $\theta = 4N_e\mu$ parameter of the corresponding chromosome computed in 100kb windows with Vcftools and using $\mu=10^{-8}$. The Padé files were created using 11 Padé coefficients as recommended. The Monte Carlo Markov chain was run for 1 million iterations with a burn-in period of 100,000 and a window size of 50 SNPs, using a block penalty (BP) of 5. A transition matrix, computed following (39) was used:

$$transition\,matrix = \begin{pmatrix} \frac{fAA}{nA\times M} & \frac{fAC}{nA\times M} & \frac{fAG}{nA\times M} & \frac{fAT}{nA\times M} \\ \frac{fCA}{nC\times M} & \frac{fCC}{nC\times M} & \frac{fCG}{nC\times M} & \frac{fCT}{nC\times M} \\ \frac{fGA}{nG\times M} & \frac{fGC}{nG\times M} & \frac{fGG}{nG\times M} & \frac{fGT}{nG\times M} \\ \frac{fTA}{nT\times M} & \frac{fTC}{nT\times M} & \frac{fTG}{nT\times M} & \frac{fTT}{nT\times M} \end{pmatrix}$$

with $fAA$, …, the number of substitution from A to A, …, computed from the polarized SNPs data; $nA$, …, the number of nucleotide A in the genome, $M$ a standardization factor corresponding to the maximum value between $(fAC + fAG + fAT)/nA$, $(fCA + fCG + fCT)/nC$, $(fGA + fGC + fGT)/nG$, and $(fTA + fTC + fTG)/nT$.

The convergence of the five independent runs of LDhelmet was estimated with Spearman's rank correlation test. The five runs were averaged together and smoothed within 2 kb, 100 kb and 1 Mb windows using custom python scripts.

We reconstructed the fine-scale recombination landscape of the European sea bass (*Dicentrarchus labrax*) to compare recombination properties in salmonids with those of a species lacking a complete *Prdm9* gene due to loss of the KRAB domain. Whole-genome haplotype data obtained via phasing-by-transmission and statistical phasing (40) was used to infer recombination in the Atlantic sea bass population with LDhelmet with a similar strategy, using the seabass_V1.0 genome assembly (GenBank accession number GCA_000689215.1) (**S5 and S6 Table**).

##### *Identification of LD-based recombination hotspots*

We identified recombination hotspots from both the raw recombination map inferred by LDhelmet (referred to as raw hotspots) and the 2-kb smoothed recombination map (referred to as 2-kb hotspots) using a sliding window approach. Hotspots were defined as intervals between two consecutive SNPs or windows of 2kb with a relative recombination rate 5-fold or higher than the mean recombination rate in the 50-kb flanking regions. When consecutive 2kb-windows exceeded the threshold, only the one with the highest rate was retained.

##### *Comparison between DSB sites and LD-based hotspots*

DSBs sites mapped with DMC1-SSDS for the pooled samples RT-52, TAC-1 and TAC-3 were compared to the LD-based recombination hotspots retrieved from the recombination landscapes of *O. mykiss*. To allow the comparison, we converted the genomic positions of the DSB hotspots mapped on the OmykA_1.1 assembly to Omyk_1.0 coordinates on which we built the LD map using the Remap program from NCBI. We compared the locations of LD-hotspots with DSBs hotspots using Bedtools intersect.

##### *Population recombination rate variation at genomic features*

We investigated how LD-based recombination rates and hotspots distribute with respect to genomic features. We first retrieved the positions of genes, exons and introns from genome annotations in each species. De novo identification of TEs families in the *O. kisutch*, *O. mykiss* and *S. salar* reference genomes was performed using RepeatModeler (v2.0.3) (41). We first used the BuildDatabase command without options to build a database from the reference genome. We ran RepeatModeler on the database generated with the LTR discovery pipeline (option -LTRStruct). We then soft-masked TEs with RepeatMasker (version 4.1.3, http://www.repeatmasker.org/, options -xsmall, -nolow) using the library of consensus sequences generated by RepeatModeler. An annotation file of the TEs and low complexity DNA sequences was generated. We retrieved the genomic location of introns from gene and exon locations, and intergenic regions from gene and TEs locations using Bedtools *subtract*. Transcription start and end sites (TSSs and TESs) were defined as the first and last positions of the genes, respectively. We predicted CpG islands (CGIs) of each reference genome with EMBOSS *cpgplot* (v6.6.0) (42), using a window size of 500 bp to calculate the percentage of GC content and the observed frequency of GCs (*-window 500*), with the minimum length of a CGI set to 250 bp (*-minlen 250*). It should be noted that the criteria that are classically used to predict CGIs in mammals or birds (CpG observed/expected ratio > 0.6, GC-content > 50%) are not appropriate for teleost fish, whose CGIs are CpG-rich but have a low GC-content (43, 44). We therefore predicted CGIs solely based on their CpG content (minimum average CpGoe < 0.6, -minoe 0.6), without any constraint on their GC-content (-minpc 0). We checked that these criteria efficiently predict TSS-associated CGIs, using whole genome DNA methylation and H3K4me3 data from rainbow trout and coho salmon (see **S2 Analysis**). TSS overlapping or not CGIs were then determined using the subsetByOverlaps function of the R package GenomicRanges (24).

We assessed population recombination rate (2-kb scale) variation according to the distance to the nearest TSS by calculating the distances of the genomic windows of the 2-kb smoothed map to the nearest TSS using the distanceToNearest. The averaged population recombination rates in genes, exons, introns, TEs, TSS overlapping or not a CGI, TES, CGIs were estimated using *subsetByOverlaps*. We compared levels of recombination rates at genomic features of the five salmonid populations to the sea bass.

We finally investigated the effect of SNP density, GC content and TEs density on population recombination rate variation and the presence of recombination hotspots. We retrieved SNP count in 100-kb and 2-kb sliding windows on the filtered SNP data containing singletons. We determined the number TEs in each window of the 100-kb smoothed maps using Bedtools *intersect* to calculate TE density. GC-content was also calculated in the windows of the 100-kb and 2-kb smoothed maps as the ratio of the sum of C and G nucleotides to the sum of the four nucleotides A, C, G, T in each window. We calculated Spearman's rank correlation between SNP density, GC content and TE density with recombination rates at the 100-kb scale. The distance of the 2-kb genomic windows to the nearest hotspots was calculated using Bedtools *closest* to assess SNP density and GC-content around recombination hotspots.

##### ***Comparison of LD-based landscapes between populations and species***

We assessed the correlation between the 100-kb smoothed recombination maps of each of the three *S. salar* populations using a Spearman's rank test. We identified shared hotspots as overlapping 2-kb hotspots using *Betdtools intersect*. We used random permutations to calculate the expected amount of hotspot overlap between the three pairs of populations. Random spots totalling the number of 2-kb hotspots were drawn one hundred times from the genome for each population using Bedtools *shuffle*, and each of these random spot sets was compared to those of the other two populations to calculate the expected average overlap between the populations.

To compare the recombination hotspots of *O. kisutch* and *O. mykiss*, whose recombination landscapes were built using their own reference genome, we used a reciprocal blast approach to retrieve the corresponding coordinates in the genome of the other species. We retrieved the fasta sequences of the *O. kisutch* 2-kb hotspots from the reference genome using Bedtools *getfasta,* and blasted with blastn on the *O. mykiss* reference genome, keeping only the best matches with *-max_targets_seq 1* and *-max_hsps 1* to retrieve the corresponding position in the *O. mykiss* genome. We retrieved the fasta sequences of the resulting blast hits in the *O. mykiss* reference genome with Bedtools *getfasta* and reciprocally blasted it on the *O. kisutch* reference genome with blastn, again retaining only the best match. This approach ensured that only reciprocal blasted positions were retained and gave the coordinates of the hotspots of *O. kisutch* on the genome of *O. mykiss*. We determined the common hotspots between *O. kisutch* (with the coordinates in the *O. mykiss* genome) and *O. mykiss* with Bedtools *intersect*. We also performed a similar reciprocal blast of *O. mykiss* 2-kb hotspots to obtain their coordinates in the *O. kisutch* genome, and determined their common hotspots. We then used the number of overlapping hotspots obtained with the *O. kisutch* coordinates, the results being similar with the coordinates in the *O. mykiss* reference genome. We also performed random permutation to obtain the distribution of shared hotspots between the two species expected by chance.

#### Identification of DNA motifs at hotspots and motif erosion

DNA motifs enriched in rainbow trout DSB hotspotsIn the rainbow trout, we used the MEME Suite (45) to detect motifs associated with DSB hotspots, focusing on the RT-52 dataset due to its high number of DSB hotspots (DMC1 peaks). Two distinct subsets of allele-specific hotspots were defined by the intersection of the DMC1 peaks with histone modification marks via BEDtools *intersect*. We then retrieved the fasta sequences using BEDtools *getfasta*.

- Allele 1 set: RT-52 DMC1 peaks (center ± 200 bp) overlapping H3K4me3 and H3K36me3 peaks from TAC-1 (N = 300)
- Allele 2 set: RT-52 DMC1 peaks (center ± 200 bp) overlapping H3K4me3 and H3K36me3 peaks from TAC-3 (N = 254).

DSB hotspots from allele 1 set showed no overlap with hotspots from TAC-3, and similarly, we did not report any overlap between allele 3 set and TAC-1 hotspots.

We used MEME-ChIP (v5.5.4) (Machanick and Bailey 2011) to screen for motifs the two allele-specific set of peaks, with the following settings:

- Discovery Mode: Classic
- Background: 2-order model from the input sequences
- Motif Width: 8-20 nt for MEME and STREME (inclusive)
- MEME Site Distribution: zero or one occurrence per sequence
- MEME Motif Count: 5 motifs

- STREME p-value Threshold: p-value ≤ 0.05
- STREME Site Positional Distribution Plots: Sequences are aligned on their centers
- CentriMo Match Score: match score ≥ 5
- CentriMo E-value Threshold: E-value ≤ 10
- CentriMo Local: central enriched regions.

Two motifs, one per allele-specific set, having high significance as well as central enrichment were further processed (**Fig 3C**). To investigate specific enrichment of the motifs at DSB and LD-based hotspots (center ± 1kb), we run FIMO (46) with a p-value cutoff of 1.0E-5 and the background model specified in the motif input. For proper comparison, fasta sequences for both DSB and LD hotspots have been collected into the assembly USDA_OmykA_1.1 using Bedtools *getFasta*, with a prior liftOver step for LD-based hotspot regions (see section "*Analysis of DMC1 ChIP-seq signal at genomic features"*).

The two sets of regions used to perform the motif discovery analysis were included as positive controls, one additional set of control sequences (*i.e.* non-hotspot regions) was generated to match the GC-content and the distance to telomeres of the positive control sets. All sets of sequences screened for the two motifs discovered by MEME-ChIP were 2kb long. We compared the frequency of sequences having at least one match to the motif for each set, to the frequency observed in the control sequences. The enrichment p-value relatively to the control was assessed by Fisher Exact test (**S25 Fig**). We used CentriMo (47) to further evaluate the central enrichment of the two motifs at the control sequences (**Fig 3C**), at the DSBs hotspots and at LD-based hotspots (**S26 Fig**), with default settings. The significance of the enrichment was automatically assessed by the tool and refers to the likelihood that the best match to the motif in a sequence occurs within the reported region.

##### *DNA motifs enriched in Atlantic salmon LD-based hotspots*

In the Atlantic salmon, we used STREME (47) to find motifs that are relatively enriched at hotspots in the three populations of *S. salar* as compared to control random spots. To generate the set of control sequences, we randomly drew windows totalling the number of hotspots from the reference genome ten times using *bedtools shuffle*, the ten datasets then being concatenated. The GC-content of the drawn sequences was calculated using Bedtools *nuc*. Windows with more than ¾ of missing data in the reference genome were removed. We retrieved a number of random windows equal to the number of hotspots with a similar GC-content distribution. The fasta sequences of the hotspots and random spots were retrieved using Bedtools *getfasta*. We ran STREME which compares the hotspot sequences to the control random sequences to find motifs between 10 and 20 bp in length. Motifs with a 2-fold enrichment or more in the hotspots compared to the control sequences, and found in more than 5% of the hotspots, were retained as potential PRDM9-binding motifs. We searched motifs in the 2-kb hotspots population-specific hotspots and in their shared hotspots.

##### *Motif erosion in European lineage of Atlantic salmon*

We then tested whether the candidate motifs showed signs of erosion in the American compared to the European lineage and conversely by comparing the number of motifs present in available long-read genome assemblies from 5 North American Atlantic salmon genomes (GCA_021399835.1, GCA_931345555.1, GCA_931345325.1, GCA_931347555.1, GCA_923944775.2) and 7 European genomes (GCA_905237065.2, GCA_931346935.2, GCA_931345835.1, GCA_931347365.1, GCA_931345645.1, GCA_931345955.1, GCA_931345925.1). To take into account potential differences in the various assemblies, we aligned these 12 genomes with SibeliaZ (48), and retrieved the motif occurrences from the collinear blocks. We ran FIMO to count motif occurrence in the aligned fraction of each genomes, using a p-value cut-off of 1.0E-7. To assess the statistical significance of motif enrichment in a lineage, a null distribution was obtained by running FIMO on 100 random permutations of the candidate motif matrix.

### 2. S2 Analysis: Prediction of CGI-associated TSSs in salmonids

#### Definition of CpG islands: constitutively hypomethylated genomic regions

Vertebrate genomes are heavily methylated, with generally more than 80% of CpG dinucleotides containing 5-methylcytosines (44, 49, 50). However, they also contain some short regions that escape DNA methylation and that are associated with H3K4me3 chromatin marks (trimethylation at lysine 4 of histone H3) (44, 51, 52). A large fraction of these non-methylated islands (NMIs) are associated to gene promoters (44). Typically, in human, mouse, chicken and zebrafish, 39% to 52% of NMIs overlap transcription start sites (TSS) of protein-coding genes, and reciprocally, 55% to 72% of protein-coding genes contain an NMI on their TSS (44). NMIs that are located in intergenic regions often show substantial variation in DNA methylation levels across tissues or during development (44). Conversely, NMIs that are associated to TSSs are maintained in the non-methylated state in most tissues, even in tissues where the corresponding gene shows no substantial transcription (44). Furthermore, the hypomethylation status of TSS-associated NMIs is often conserved across vertebrate species, which indicates that they are epigenetically stable not only across tissues but also through evolutionary time (44).

Methylated cytosines are hypermutable, which causes an overall depletion of CpG dinucleotides in vertebrate genomes, except in regions that escape DNA methylation (51, 53). This explains why NMIs generally display a much higher frequency of CpG dinucleotides compared to the rest of the genome (44). The term ‘CpG island’ (CGI) was coined to refer to these unmethylated CpG-rich loci (51). It should be noted that loci that are unmethylated in some tissues but methylated in the germline are not protected from CpG losses over evolutionary time. Thus, CGIs are expected to correspond to the subset of NMIs that are unmethylated in the germline, and this, stably over time. This explains why TSS-associated NMIs, which generally show a conserved and constitutive pattern of hypomethylation (44), are particularly CpG-rich (51).

#### Bioinformatic prediction of CpG islands

In mammals and birds, CGIs are also characterized by a relatively high G+C content (51). There is now clear evidence that this feature results from the process of GC-biased gene conversion (gBGC), which is induced by an increased level of meiotic recombination in germline NMIs (54-61).

Grounded on these observations, originally made in mammals and birds, algorithms have been developed to predict CGIs based solely on their CpG and G+C content. We will hereafter use the term ‘*pCGI*’ to refer to these putative CGIs, predicted bioinformatically, without any direct measurement of their DNA methylation level. Classically, sequences of more than 200 bp, with a G+C content above 50%, and a ratio of observed over expected CpG content ($CpG_{oe}$) above 0.6 are classified as *pCGI*s (62). Notably, the UCSC genome browser uses these criteria to annotate *pCGI*s in vertebrate genomes. These *pCGI*s are commonly used to investigate the genomic features associated to CGIs, in genomes for which DNA methylation or H3K4me3 data are not readily available.

It should be noted however that not all *pCGI*s correspond to *bona fide* CGIs. The overlap between experimentally characterized NMIs and *pCGI*s was analyzed in seven vertebrate species (44). In human, mouse and chicken, most *pCGI*s encompassed NMIs.

Thus, in these species, *pCGI*s are indeed good predictors of CGIs. However, in other species (platypus, green anole lizard, xenopus and zebrafish) a majority of *pCGI*s do not correspond to NMIs (44). Furthermore, in zebrafish, NMIs have a high $CpG_{oe}$, but a low G+C content (44). Interestingly, an early study, based on a limited number of genes, already had suggested that CGIs from carp and trout were not G+C-rich (43). These observations imply that the criteria that are classically used to annotate *pCGI*s in mammals or birds ($CpG_{oe}$ >0.6 and G+C>50%) are not appropriate to predict CGIs in teleost fish.

#### Prediction of CGI-associated TSSs in salmonids

In our study, we wanted to identify the subset of TSSs that are associated to a CGI in salmonid genomes, to investigate whether they show an elevated recombination rate. Since DNA methylation and H3K4me3 data were available only for a fraction of the species that we wanted to analyze, we sought to predict CGI-associated TSSs based on their base composition. However, given the observations mentioned above, we first explored the criteria to be used to annotate *pCGI*s in salmonids.

For this, we used DNA methylation data available in the coho salmon (*Oncorhynchus kisutch*) (63) to investigate the relationship between the base composition of promoter regions and their DNA methylation level. Overall, the coho salmon genome is heavily methylated. For instance, in the liver sample that we analyzed, 65.2% of CpG sites show a high methylation level (>0.6), 8.1% have an intermediate methylation level ([0.2,0.6]) and 26.7% are hypomethylated (<0.2). We measured methylation levels in promoter regions (defined as the 500 bp upstream of the TSS) of protein-coding genes (N=27,832). Similar to other vertebrates, a large fraction of promoter regions (59.3%) are hypomethylated (**S1A Fig in S2 Analysis)**. As expected, hypomethylated promoters have a higher $CpG_{oe}$ than highly methylated promoters (p-value Student's t test < 1e-10) (**S1C Fig in S2 Analysis**). However, contrarily to amniotes, hypomethylated promoters are not G+C rich: they rather tend to have a lower G+C content than highly methylated promoters (p-value Student's t test < 1e-10) (**S1D Fig in S2 Analysis**). Overall, only 1.1% of promoters (and 1.4% of hypomethylated promoters) match the criteria classically used to identify *pCGI*s ($CpG_{oe}$>0.6 and G+C>50%, **S1B Fig in S2 Analysis**).

Thus, as previously reported for zebrafish (44), a high G+C content is not an appropriate criterion to predict CGIs in salmonids. We therefore tried to predict CGIs solely based on their $CpG_{oe}$. We used the *cpgplot* software to identify DNA segments matching the following criteria:

- length > 250 bp
- $CpG_{oe}$ > 0.6

These DNA segments will hereafter be referred to as 'fish putative CGIs' (*fpCGIs*).

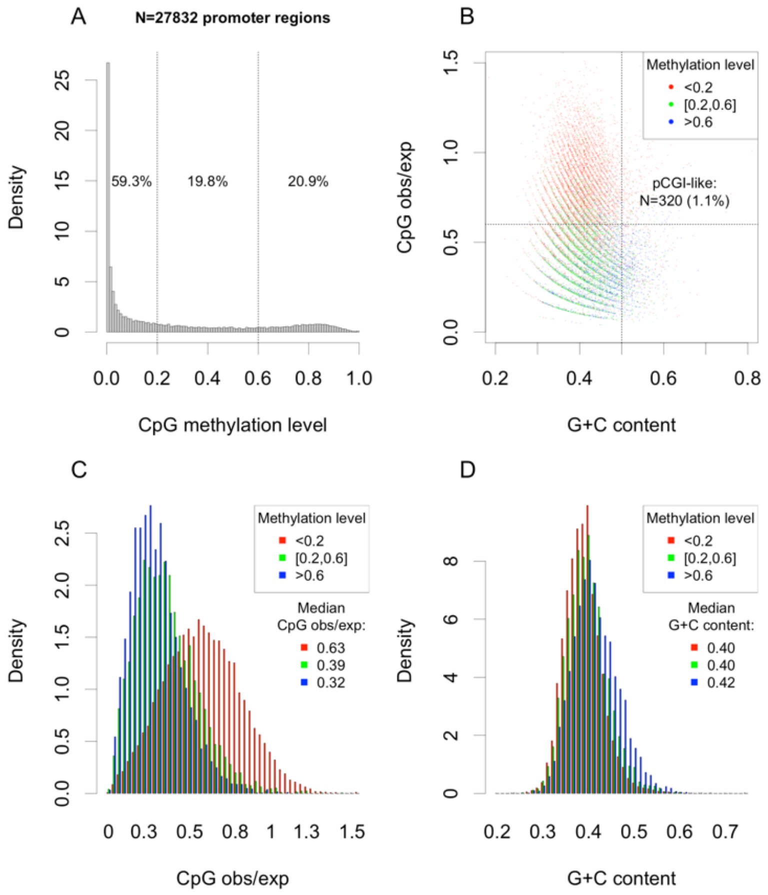


**Figure S1: Relationship between base composition and DNA methylation level in promoter regions of coho salmon (*Oncorhynchus kisutch*).** We selected protein-coding genes (N=27,832) annotated in the coho salmon reference genome assembly and extracted their promoter region, defined as the 500 bp upstream of the TSS. For each promoter region, we quantified the average methylation level at CpG sites, using DNA methylation data from a liver sample (63). **A)** Distribution of CpG methylation level within promoter regions. **B)** CpG observed/expected ratio vs. G+C content of promoter regions. Promoters were classified according to their CpG methylation level (red: hypomethylated; blue: highly methylated; green: intermediate methylation level). The number and percentage of promoter regions matching the classical criteria for CGI annotation (CpG obs/exp>0.6 and G+C content >0.5) are indicated. **C)** Distribution of CpG observed/expected ratio of promoter regions, for different classes of CpG methylation level. **D)** Distribution of G+C content of promoter regions, for different classes of CpG methylation level.

We identified 667,422 *fpCGIs* in the coho salmon genome, and among the 27,832 annotated TSSs, 13,723 (49.3%) are located close to a *fpCGI* (<250 bp). The total number of *fpCGIs* largely exceeds the number of NMIs reported in vertebrate genomes (~11,000 to 41,000 NMIs) (44), which suggests that a large fraction of *fpCGIs* do not correspond to *bona fide* CGIs. However, we observed that the presence of a *fpCGI* is informative regarding the epigenetic status of promoter regions: indeed, among TSSs located close to a *fpCGI*, 81.9% are hypomethylated, whereas only 13.8% of TSSs located far from a *fpCGI* (>1000 bp) are hypomethylated (**S2 Fig in S2 Analysis**). This 5.9-fold enrichment indicates that the presence of a *fpCGI* close to the TSS is a very good predictor of the methylation state of the promoter region.

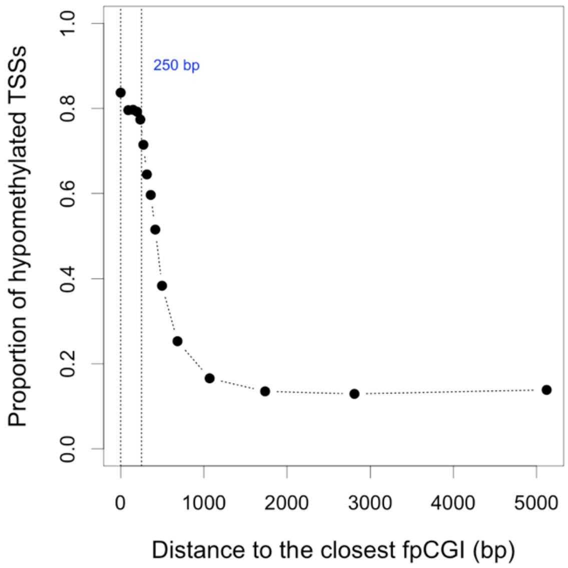


**Figure S2: Relationship between DNA methylation level in the promoter regions of coho salmon and the presence of a nearby *fpCGI*.** We classified TSSs according to their distance to the nearest *fpCGI*, and computed the proportion of hypomethylated TSSs in each bin.

To determine whether the presence of a *fpCGI* also predicts histone epigenetic marks of promoter regions, we analyzed H3K4me3 ChIPseq data from a rainbow trout (*Oncorhynchus mykiss*) brain sample (unpublished data, produced in the context of the Aqua-FAANG project, kindly provided by Lien Sigbjorn). We analyzed the distribution of H3K4me3 ChIPseq signal in promoter regions of protein-coding genes (N=40,786). Overall, 52.1% of promoter regions show a strong H3K4me3 ChIPseq signal (**S3A Fig in S2 Analysis**). These promoter regions with strong H3K4me3 marks display a high $CpG_{oe}$ and a relatively low G+C content (**S3B, S3C and S3D Fig in S2 Analysis**).

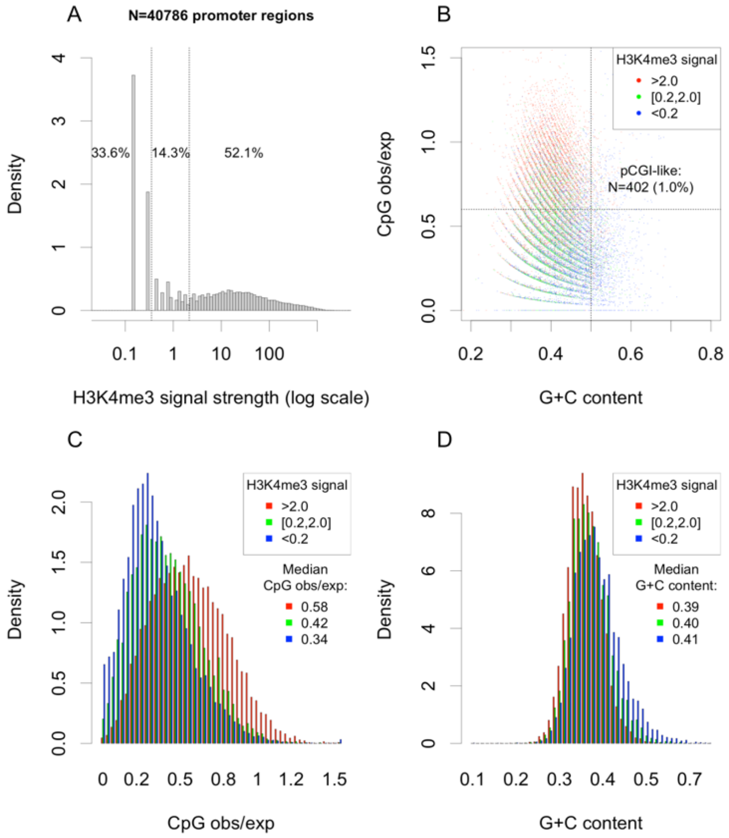


**Figure S3: Relationship between base composition and H3K4me3 marks in promoter regions of the rainbow trout (*Oncorhynchus mykiss*).** We selected protein-coding genes (N=40,786) annotated in the rainbow trout genome and extracted their promoter region, defined as the 500 bp upstream of the TSS. For each promoter region, we quantified the H3K4me3 level (ChIPseq data from a brain sample). **A)** Distribution of H3K4me3 signal within promoter regions. **B)** CpG observed/expected ratio vs. G+C content of promoter regions. Promoters were classified according to their CpG methylation level (red: hypomethylated; blue: highly methylated; green: intermediate methylation level). The number and percentage of promoter regions matching the classical criteria for CGI annotation (CpG obs/exp>0.6 and G+C content >0.5) are indicated. **C)** Distribution of CpG observed/expected ratio of promoter regions, for different classes of H3K4me3 signal. **D)** Distribution of G+C content of promoter regions, for different classes of H3K4me3 signal.

We identified 560,469 *fpCGIs* in the rainbow trout genome, and among the 40,786 annotated TSSs, 18,356 (45.0%) are located close to a *fpCGI* (<250 bp). Again, we observed that the presence of a *fpCGI* is informative regarding the epigenetic status of the promoter region: among *fpCGI*-associated TSSs, 69.4% display a strong H3K4me3 signal, compared to only 7.8% for TSSs located far from a *fpCGI* (>1000 bp) (**S4 Fig in S2 Analysis**). This 8.9-fold enrichment indicates that the presence of a nearby *fpCGI* is a very good predictor of the chromatin state of the promoter region.

CpG-island annotations were available from UCSC for the same rainbow trout genome assembly (N=18,220 *pCGIs*), allowing us to evaluate the capacity of this commonly used resource to predict salmonid CGIs. Only 0.7% of rainbow trout TSSs are located at less than 250bp from a UCSC *pCGI*. Among these *pCGI*-associated TSSs, 53.9% display a strong H3K4me3 signal (compared to 69.4% for *fpCGI*-associated TSSs). These results indicate i) that a majority of TSS-associated CGIs are missed by UCSC *pCGI* annotations, and ii) that the *fpCGIs* are better predictors of the epigenetic state of promoters than UCSC *pCGIs*.

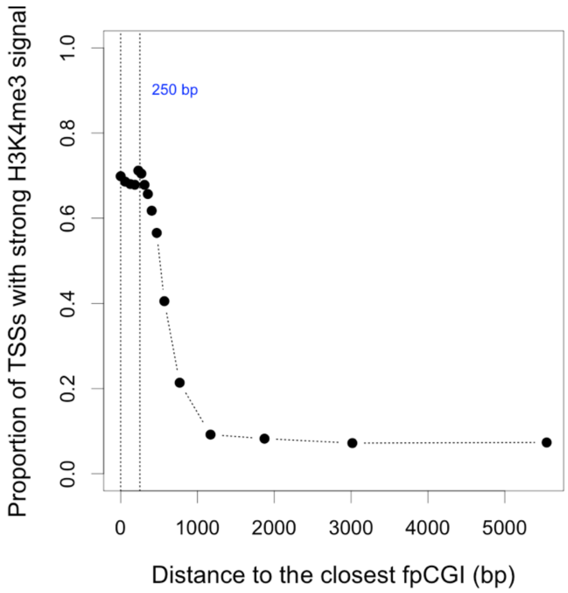


**Figure S4: Relationship between H3K4me3 marks in the promoter regions of the rainbow trout and the presence of a nearby *fpCGI*.** We classified TSSs according to their distance to the nearest *fpCGI*, and computed in each bin, the proportion of TSS with a strong H3K4me3 signal in their promoter region.

#### Conclusion

Both in rainbow trout and in coho salmon, we observed that 52% to 59% of promoter regions present the typical hallmarks of CGIs: low DNA methylation level, high signal of H3K4me3 marks, high $CpG_{oe}$. However, as in zebrafish, but contrary to mammals and birds, these GCIs are not G+C-rich. Hence, the criteria that are used classically to predict CGIs ($CpG_{oe}$>0.6 and G+C>50%) fail to detect a large majority of the CGIs present in salmonid genomes. We therefore propose that in these species, CGIs should be predicted solely based on their $CpG_{oe}$, not on their G+C-content. The very high number of *fpCGI*s identified in the whole genome (>500,000 *fpCGI*s) suggests that many of them are false positives. However, we showed that the presence or absence of *fpCGI*s in the vicinity of TSSs is a very good predictor of their epigenetic status (DNA methylation level, H3K4me3 marks) of promoter regions. Hence, the criteria that we used to annotate *fpCGI*s (DNA segment > 250 bp, with $CpG_{oe}$>0.6) appear appropriate to predict CGI-associated promoters in salmonids.

#### Material and methods

##### *Reference genome assembly, annotation of TSSs and promoter regions*

We retrieved genome sequences and annotations from NCBI:

- Coho salmon (*Oncorhynchus kisutch*): Okis_V1 (GCF_002021735.1)
- Rainbow trout (*Oncorhynchus mykiss*): USDA_OmykA_1.1 (GCF_013265735.2)

For each protein-coding gene, we selected the transcript encoding the longest CDS, and defined the TSS as the 5'end of this transcript. We excluded genes located on the mitochondrial genome, or on unmapped contigs.

We defined the 'promoter region' as the 500bp-long segment in 5' of the TSS. We computed the G+C content of sequences, and excluded promoter regions containing more than 100 undetermined bases (N's).

The CpGoe of promoter regions was calculated according to the formula :

```
CpGoe = Number of CpG * L / (Number of C * Number of G)
```

where L = length of sequence.

##### *Prediction of CGIs*

We used the *cpgplot* software (from the EMBOSS package) to identify genomic DNA segments matching the following criteria:

- length > 250 bp
- CpGoe > 0.6

We used the following command line:

```
cpgplot -sequence genome.fa -minlen 250 -minpc 0. -minoe 0.6 -window 500 -noplot
```

These DNA segments will hereafter be referred to as 'fish putative CGIs' (*fpCGIs*).

In addition, for the rainbow trout, we retrieved annotated *pCGIs* from the UCSC genome browser: http://genome.ucsc.edu/cgi-bin/hgTables?hgsid=1701022328_0XcuiZkramud5mBRGsZxZ4CT99HY&clade=hub_2243217&org=hub_2243217_USDA_OmykA_1.1+Sep.+2020&db=hub_2243217_GCF_013265735.2&hgta_group=allTracks&hgta_track=hub_2243217_cpgIslands&hgta_table=0&hgta_regionType=genome&position=NC_048566.1%3A34%2C602%2C292-34%2C612%2C292&hgta_outputType=sequence&hgta_outFileName=

These UCSC *pCGIs* have been predicted based on the 'classical' criteria (CpGoe > 0.6 and G+C content > 0.5).

##### *DNA methylation and H3K4me3 data*

We retrieved DNA methylation data from coho salmon, obtained by whole genome bisulfite sequencing (63)(NCBI project accession PRJNA678281). The BED file of a liver sample (NCBI biosample accession SAMN25653842) was kindly provided by Maeva Leitwein.

We retrieved H3K4me3 ChIPseq data from a rainbow trout brain sample (unpublished data, produced in the context of the Aqua-FAANG project, kindly provided by Lien Sigbjorn).

### 3. Supplementary Figures

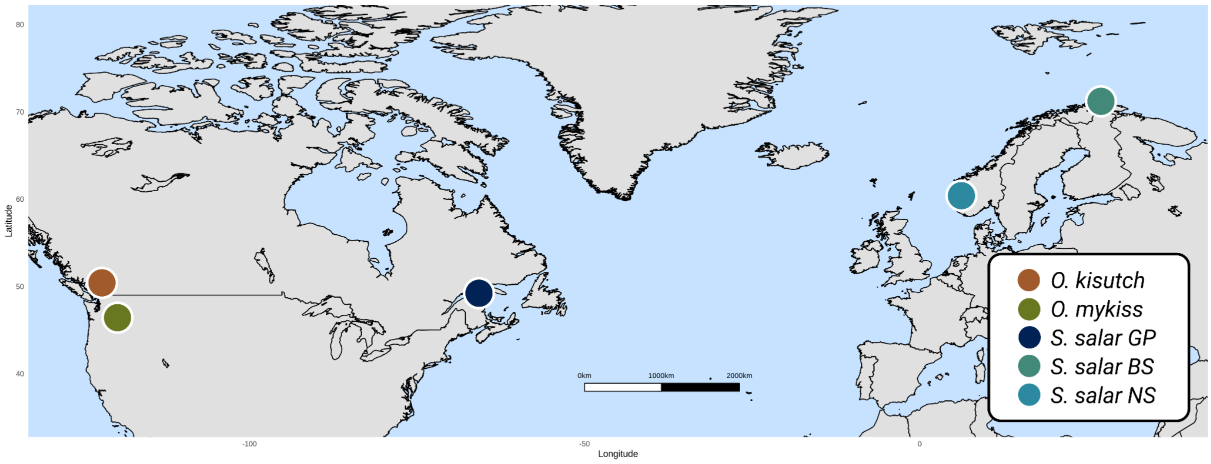


**Figure S1: Sample location.** The 20 individuals of *O. kisutch* were samples in the Columbia River (in orange) (27), the 22 samples of *O. mykiss* come from North America rivers (in green) (28), and the 60 individuals from Canada were sampled in Canada and Norway (29). Based on population structure analysis, we subdivided into three populations (in shades of blue): Gaspesie-Anticosti (GP) Barents sea (BS) and North sea (NS).

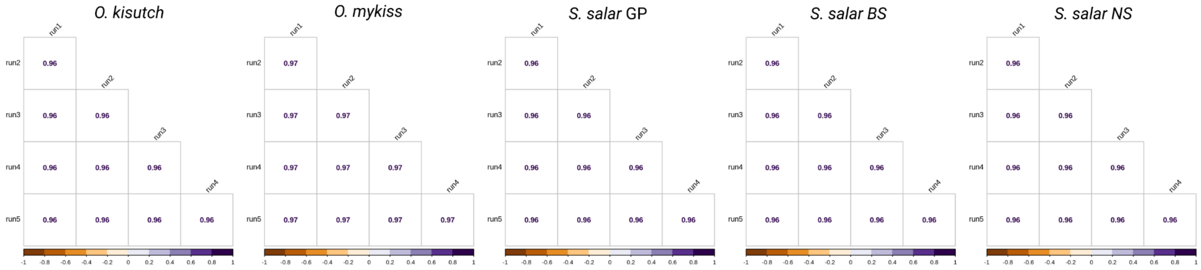


**Figure S2: Pairwise correlation between the five independent runs of LDhelmet.** Spearman's rank correlation matrix for the five populations, p-value < 0.05

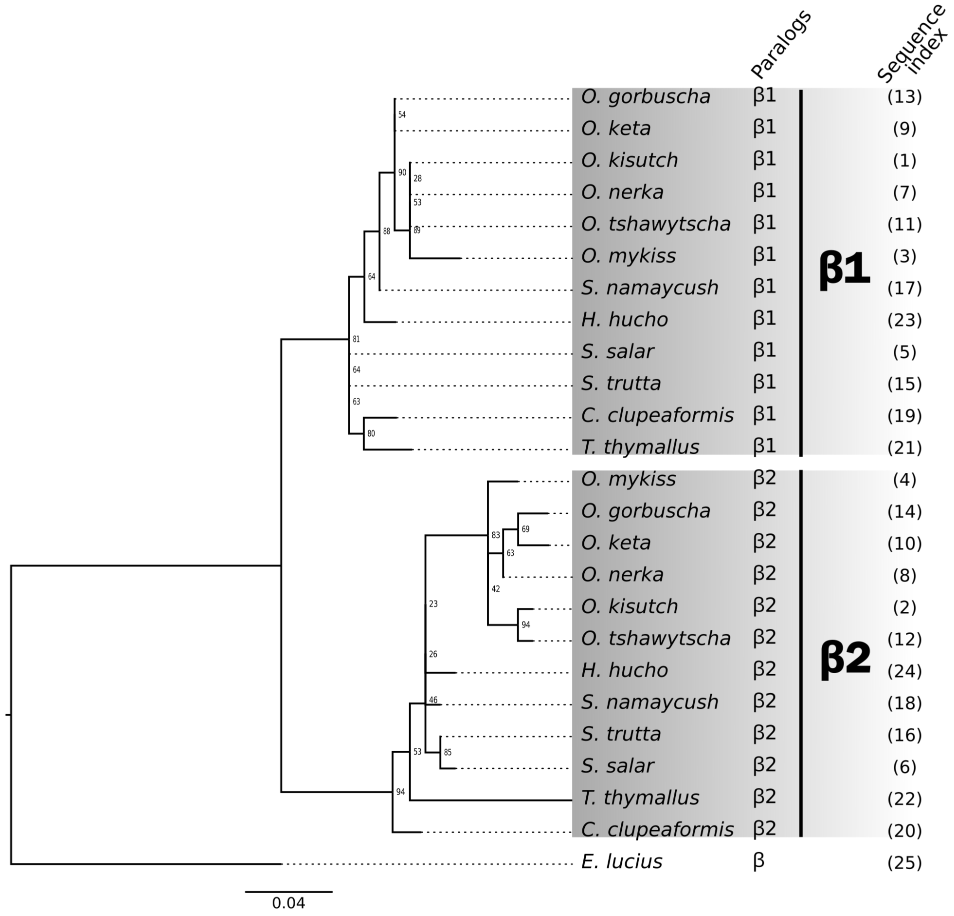


**Figure S3: Phylogenetic distribution of PRDM9β paralogs in twelve salmonids** (i.e. *Oncorhynchus kisutch*, *Oncorhynchus mykiss*, *Oncorhynchus nerka*, *Oncorhynchus keta*, *Oncorhynchus gorbuscha*, *Oncorhynchus tshawytscha*, *Salmo salar*, *Salmo trutta*, *Salvelinus namaycush*, *Hucho hucho*, *Coregonus clupeaformis*, *Thymallus thymallus*) and the northern pike (*Esox lucius*) as the outgroup species. The phylogenetic tree was realised with IQTREE on the concatenated 2 exons of the SET domain, with 1000 bootstrap replicates (values shown at nodes). All β copies have lost KRAB and SSXRD domains. In column from left to right, i) species; ii) annotated paralog copy; iii) the two major β clusters; iv) indexes referring to the **S7 Table** with additional information on the corresponding copy. The scale bar is in unit of substitution per site.

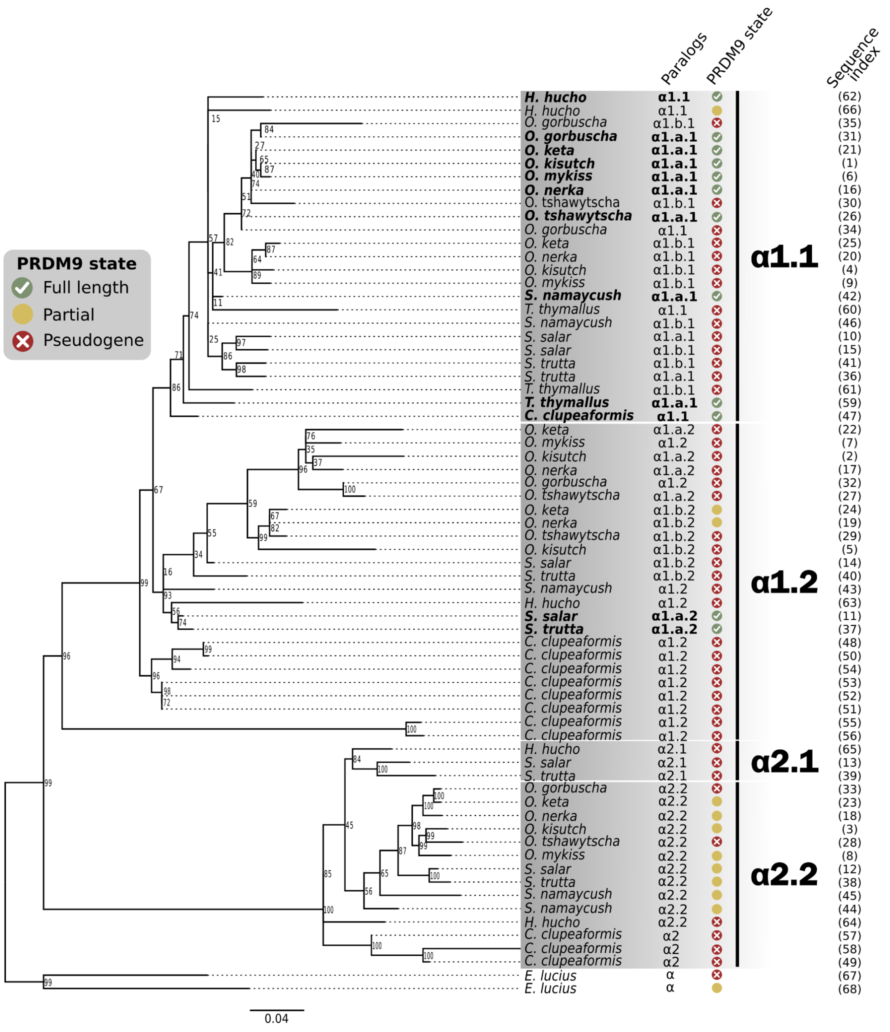


**Figure S4: Phylogenetic distribution of PRDM9α paralogs in twelve salmonids** (*i.e. Oncorhynchus kisutch*, *Oncorhynchus mykiss*, *Oncorhynchus nerka*, *Oncorhynchus keta*, *Oncorhynchus gorbuscha*, *Oncorhynchus tshawytscha*, *Salmo salar*, *Salmo trutta*, *Salvelinus namaycush*, *Hucho hucho*, *Coregonus clupeaformis*, *Thymallus thymallus*) and the northern pike (*Esox lucius*) as the outgroup species. The phylogenetic tree was realised with IQTREE on the concatenated 6 exons of the 3 canonical domains KRAB, SSXRD and SET, with 1000 bootstrap replicates (values shown at nodes). In column from left to right, i) species name; ii) annotated paralog copy, iii) PRDM9 copy status (*i.e.* full-length protein containing at least the three canonical domains, between 9 and 10 exons, with no evidence of pseudogeneisation, highlighted in bold; partial protein lacking exons but without evidence of pseudogeneisation; pseudogenes revealed by the presence of stop codon or frameshift); iv) the four major α clusters, and v) indexes referring to the **S7 Table** with additional information on the corresponding copy. The scale bar is in unit of substitution per site.

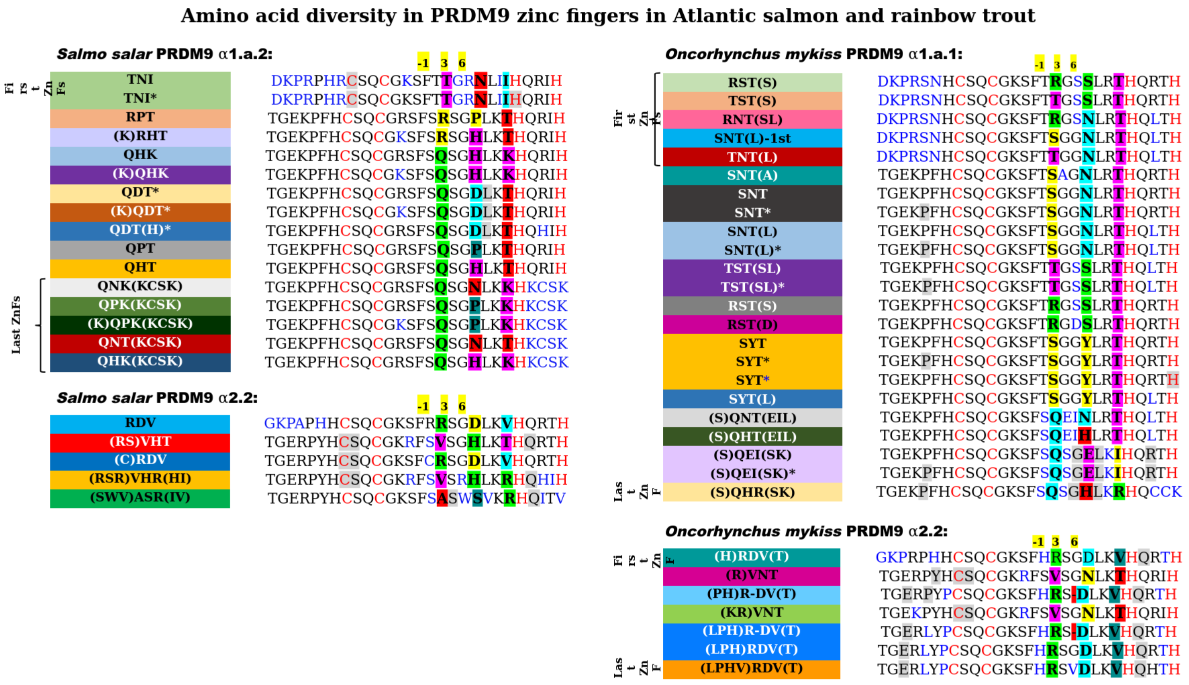


**Figure S5: Amino-acid diversity in full-length and partial PRDM9 zinc fingers in *S. salar* and *O. mykiss*.** Amino acid sequences of all unique zinc fingers found in alleles identified in *S. salar* PRDM9 α1.a.2 and α2.2, and in *O. mykiss* PRDM9 α1.a.1 and α2.2 (**Fig 2A**, **S7 Fig**). In bold colored boxes are indicated the 3 hypervariable DNA-binding residues. In red are reported the cysteine (C) and histidine (H) residues involved stabilizing the structure of the array. In blue are indicated the polymorphic residues compared to the consensus, outside the 3 amino-acids in contact with DNA. In shaded grey are reported the synonym variations in respect to the consensus. The complementary information about the DNA sequences of all alleles identified is available in the Supplementary Methods.

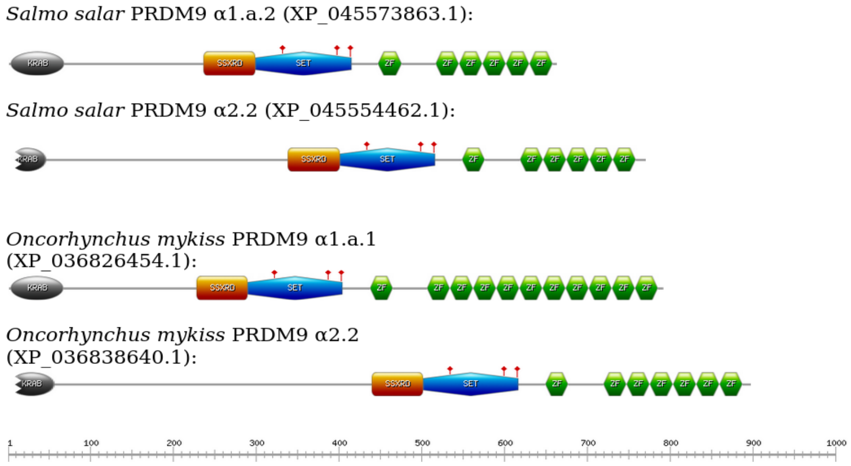


**Figure S6: Graphical view of PRDM9 paralogs.** Cartoon showing the functional domains of PRDM9 paralogs analyzed in this study. The amino acid sequences were obtained from the reference genome and analyzed using previously described methodology (64). Both α2.2 copies possess a partial KRAB domain, and we refer to these copies as partial PRDM9. All four copies present the three catalytic tyrosine residues in the SET domain, required for methyltransferase activity.

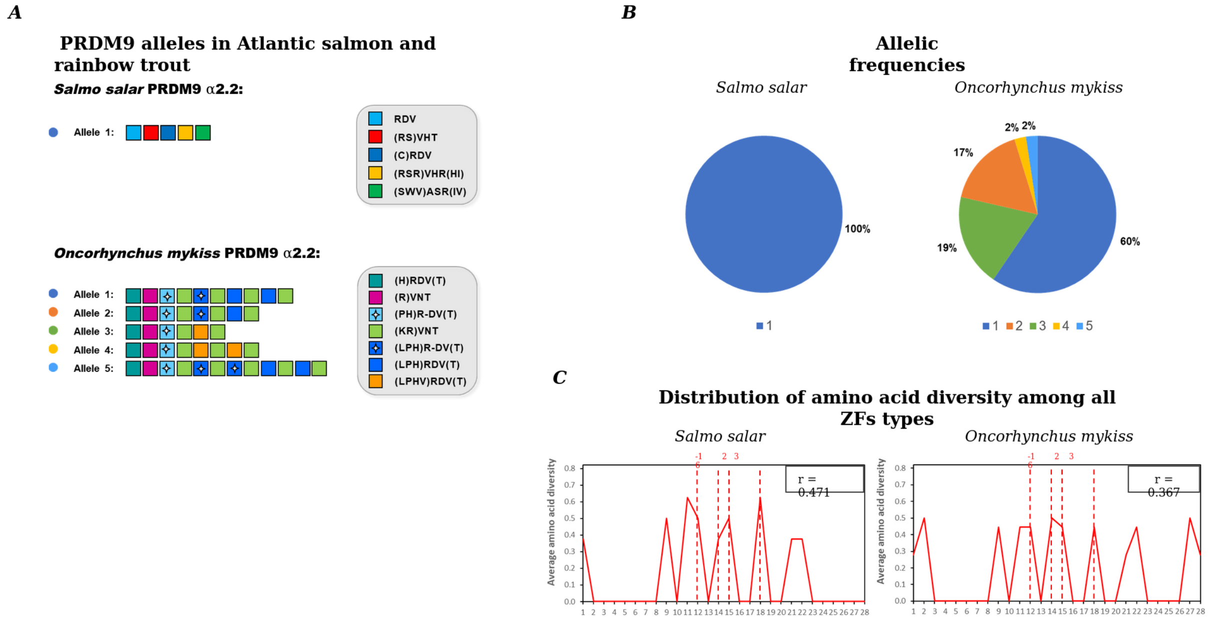


**Figure S7: Partial PRDM9 (KRAB-less) zinc finger allelic diversity in *S. salar* and *O. mykiss*. A)** Structure of PRDM9 zinc finger arrays of identified alleles in *S. salar* PRDM9 α2.2 and *O. mykiss* PRDM9 α2.2. Colored box represent unique zinc fingers, characterized by the 3 amino-acids in contact with DNA (3-letter code). Additional variation relative to a reference sequence is indicated between brackets. A white star indicates the zinc fingers missing one amino-acid residue (27 a.a. instead of 28). The complete zinc finger amino-acid sequences are shown in **S5 Fig**. **B)** Frequencies of the alleles displayed on panel A among the 20 *S. salar* and 20 *O. mykiss* individuals that were genotyped for PRDM9. **C)** Distribution of amino-acid diversity among all unique zinc fingers found in alleles displayed on panel A, following previously described methodology (10). The amino acid diversity is plotted as a function of amino acid position in the ZF alignment, ranging from position 1 to position 28 (first and last residues) of a ZF unit. The ratio of amino acid diversity at DNA-binding residues of the ZF array (-1, 2, 3 and 6), indicated as r, is shown in the upper box.

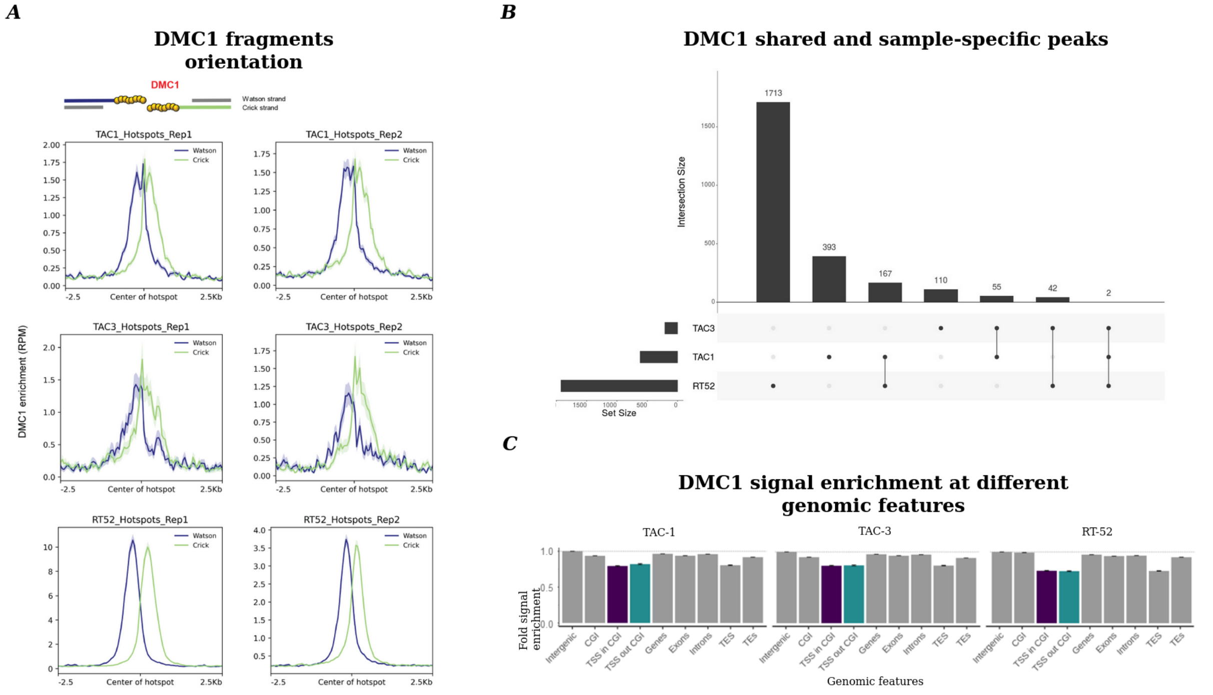


**Figure S8: Meiotic DSB hotspots features in *O. mykiss*. A)** Average profile of DMC1 ChIP-seq ssDNA fragments orientation in TAC-1, TAC-3 and RT-52 testes, at DSB hotspots detected in TAC-1, TAC-3 and RT-52. The profile from each experiment performed is shown (two replicates/sample). Signal mapped on the forward strand is depicted in blue, signal aligned to the reverse strand is shown in green, as shown in the cartoon on top of the panel. **B)** Upset plot showing intersections between DSB hotspots from TAC-1 (n=616), TAC-3 (n=209) and RT-52 (n=1924). **C)** DMC1 ChIP-seq signal fold enrichment (scaled by the average signal in intergenic regions) at multiple genomic features. TSS inside and outside CGIs are highlighted in purple and turquoise, respectively.

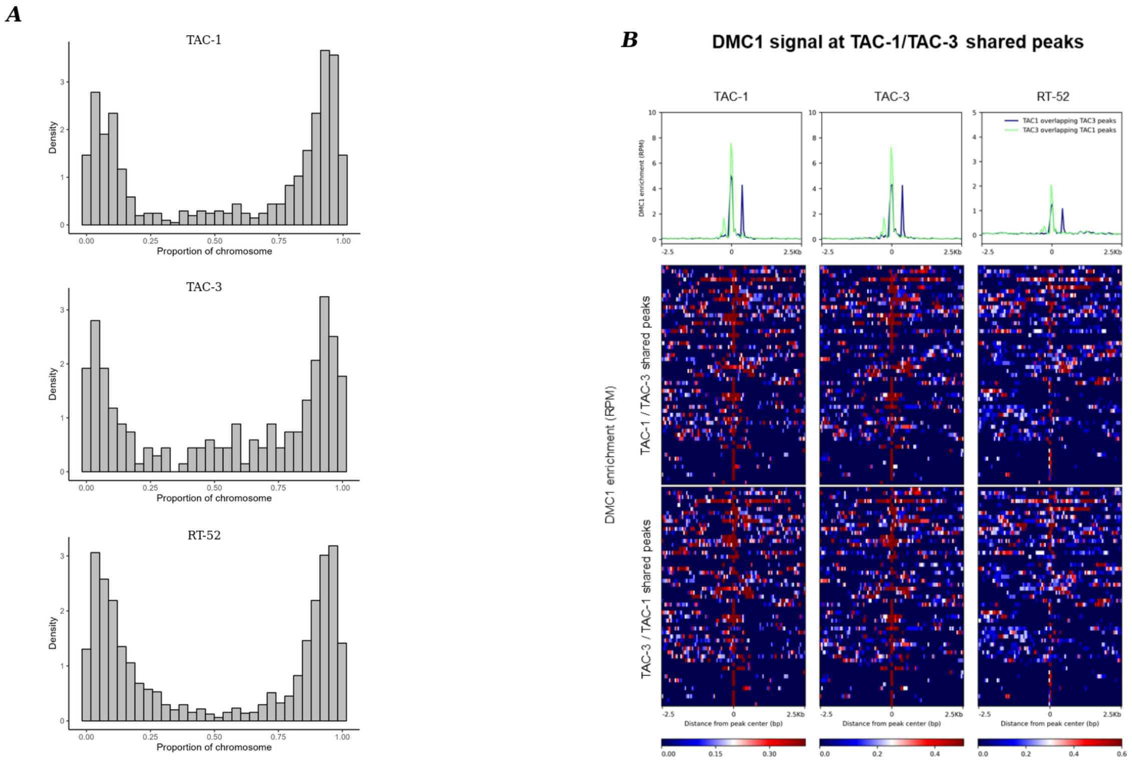


**Figure S9: Distribution of DSB hotspots along chromosomes in *O. mykiss*. A)** Distribution of DSB hotspots from TAC-1, TAC-3 and RT-52 along chromosomes (paces of 1/30 of chromosome length). **B)** DMC1 signal at TAC-1/TAC-3 shared peaks**.** Average profile and heatmap of DMC1 ChIP-seq ssDNA signal in TAC-1, TAC-3 and RT-52 testes, at shared DMC1 peaks: TAC-1 intersecting TAC-3 (blue, n=55), or TAC-3 intersecting TAC-1 (green, n=55).

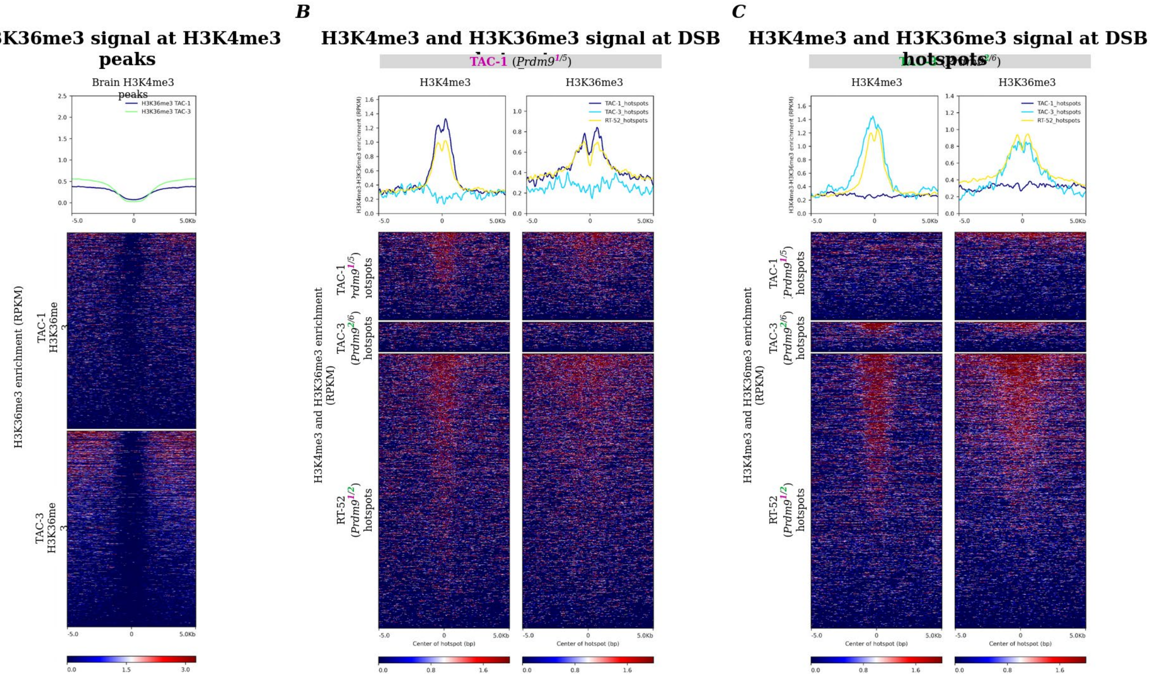


**Figure S10: Histone marks signal at H3K4me3 peaks and at DSB hotspots. A)** Average profile and heatmap of H3K36me3 ChIP-seq signal in TAC-1 (blue) and TAC-3 (green) testes, at H3K4me3 peaks detected in brain (Aqua-FAANG). **B)** Average profile and heatmap of H3K4me3 (left) and H3K36me3 (right) ChIP-seq signal in TAC-1 testes, at DSB hotspots detected in TAC-1 (blue), TAC-3 (cyan) and RT-52 (yellow). **C)** Average profile and heatmap of H3K4me3 (left) and H3K36me3 (right) ChIP-seq signal in TAC-3 testes, at DSB hotspots detected in TAC-1 (blue), TAC-3 (cyan) and RT-52 (yellow).

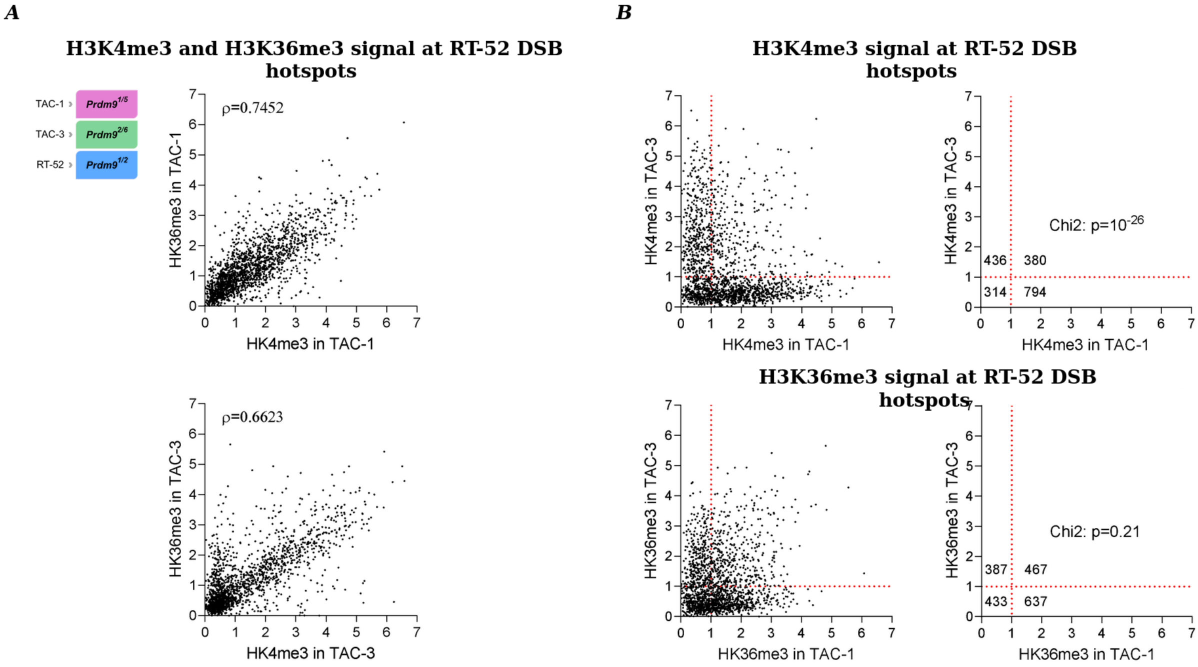


**Figure S11: Correlation of H3K4me3 and H3K36me3 signal at RT-52 hotspots. A)** Scatterplots showing H3K4me3 and H3K36me3 ChIP-seq signal in TAC-1 and TAC-3 testes, at RT-52 DSB hotspots. **B)** Left panels, scatterplots representing H3K4me3 (top) or H3K36me3 (bottom) ChIP-seq signal in TAC-1 and TAC-3 testes, at RT-52 DSB hotspots. Right panels, numbers of RT-52 hotspots with H3K4me3 (top) or H3K36me3 (bottom) ChIP-seq signal under or above 1 in TAC-1 and TAC-3. Chi-square test of homogeneity.

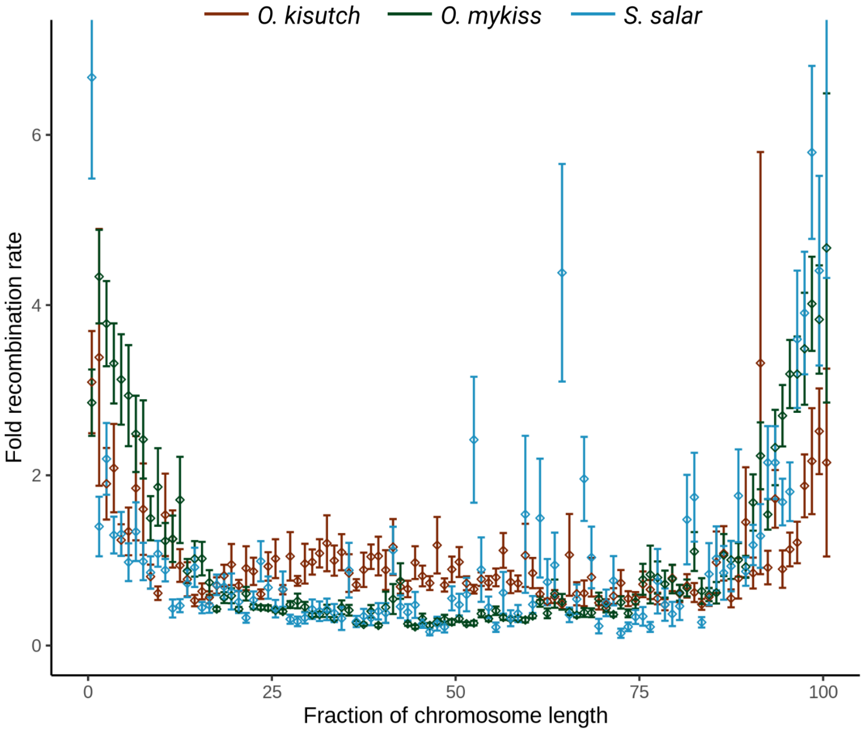


**Figure S12: Broad scale recombination rate variations along the genome.** Recombination rates were averaged into percentiles of chromosome length, and scaled by the genomic mean for *O. kisutch* (in orange), *O. mykiss* (in green) and *S. salar* (in blue, only the NS population is shown).

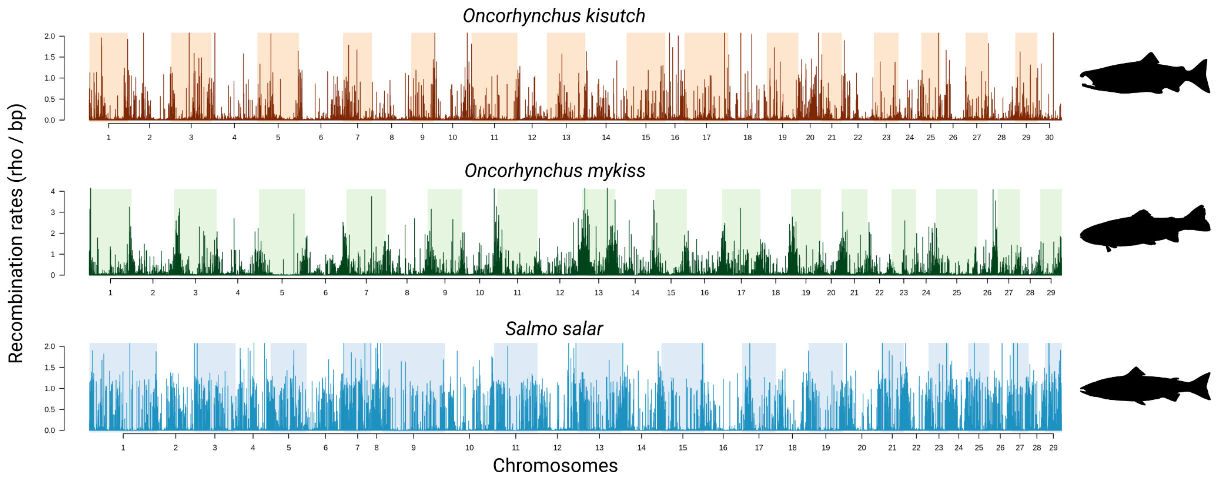


**Figure S13:** Fine-scale recombination landscapes of *O. kisutch* (in orange), *O. mykiss* (in green) and *S. salar* (in blue, only the NS population is shown), with recombination rates smoothed in 2 kb sliding windows.

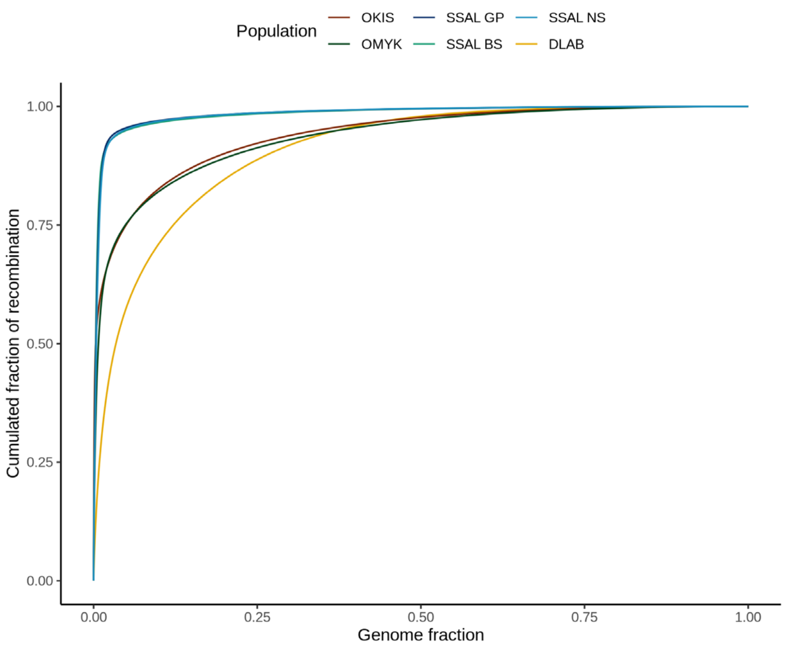


**Figure S14:** Proportion of recombination according to proportion of the genome for *O. kisutch* (orange), *O. mykiss* (green), *S. salar* (shades of blue) and *D. labrax* (gold). OKIS = *O. kisutch*, OMYK = *O. mykiss*, SSAL = *S. salar* (GP, BS and NS populations), DLAB = *D. labrax*.

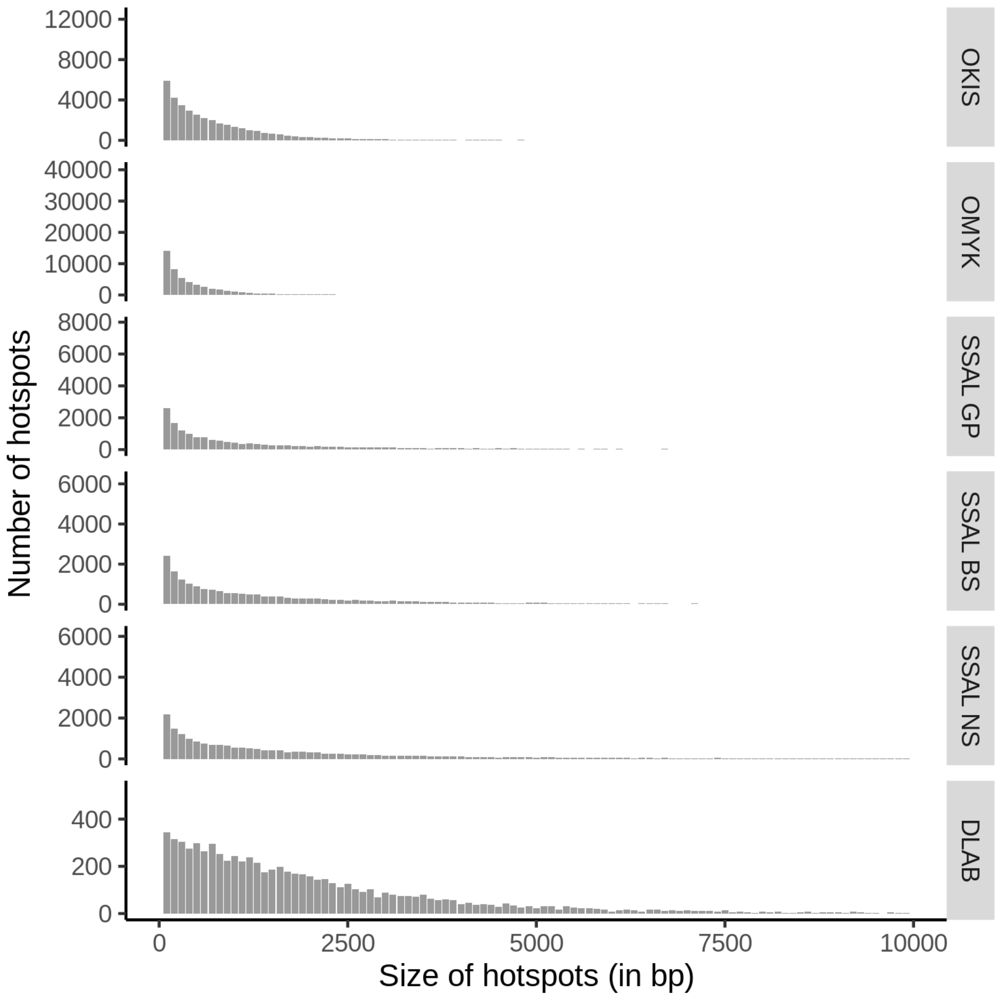


**Figure S15:** Size distribution of raw hotspots. Hotspots were defined as consecutives windows of two adjacent SNPs in which the recombination rate is at least 5-fold higher than the 50-kb flanking regions. OKIS = *O. kisutch*, OMYK = *O. mykiss*, SSAL = *S. salar* (GP, BS and NS populations), DLAB = *D. labrax*.

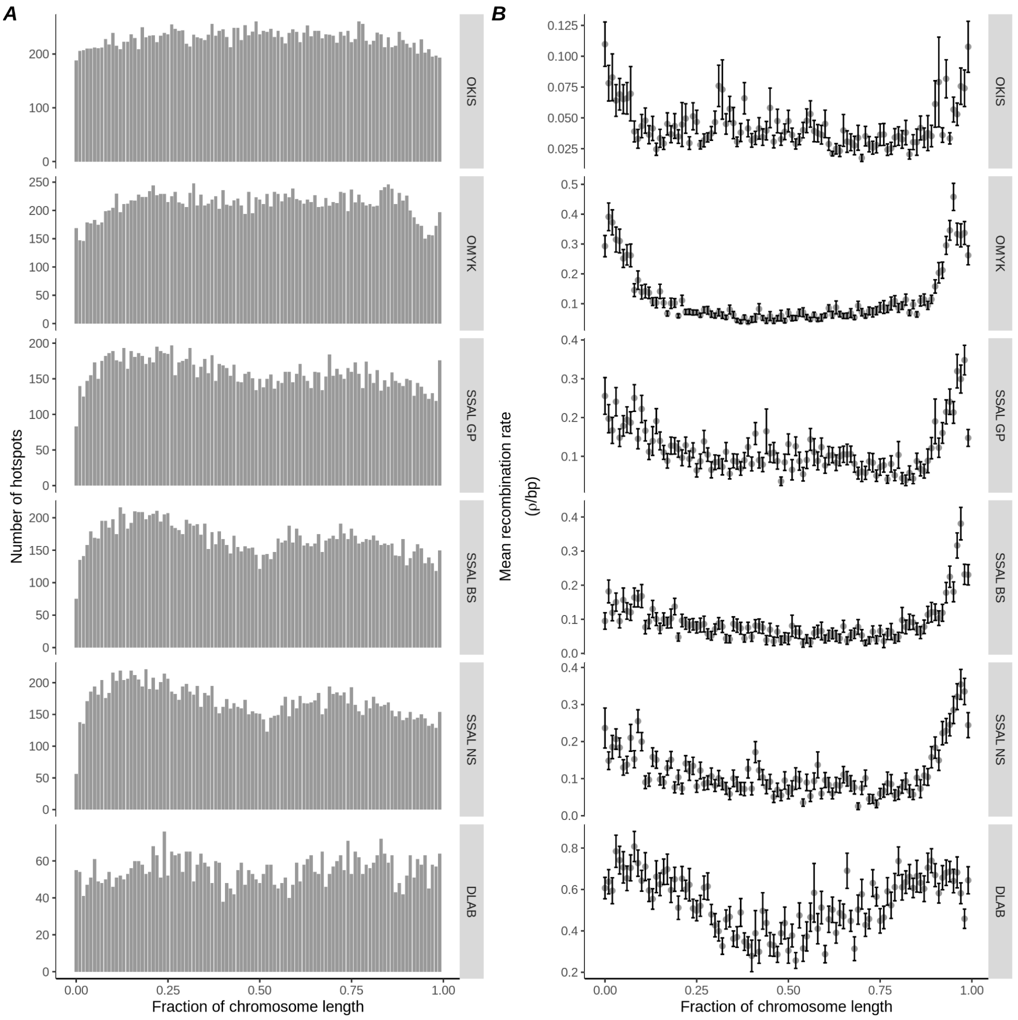


**Figure S16: Chromosome-wide distribution of 2-kb hotspots.** Hotspots were defined as 2-kb windows in which the recombination rate at least is 5-fold higher than the 50-kb flanking regions. If several consecutives windows passed the threshold, the heatest was retained. **A)** Distribution of hotspot counts along chromosomes. The number of hotspots has been averaged in percentiles of chromosome length. **C)** Recombination rates in hotspots averaged in percentiles of chromosome length. OKIS = *O. kisutch*, OMYK = *O. mykiss*, SSAL = *S. salar* (GP, BS and NS populations), DLAB = *D. labrax*.

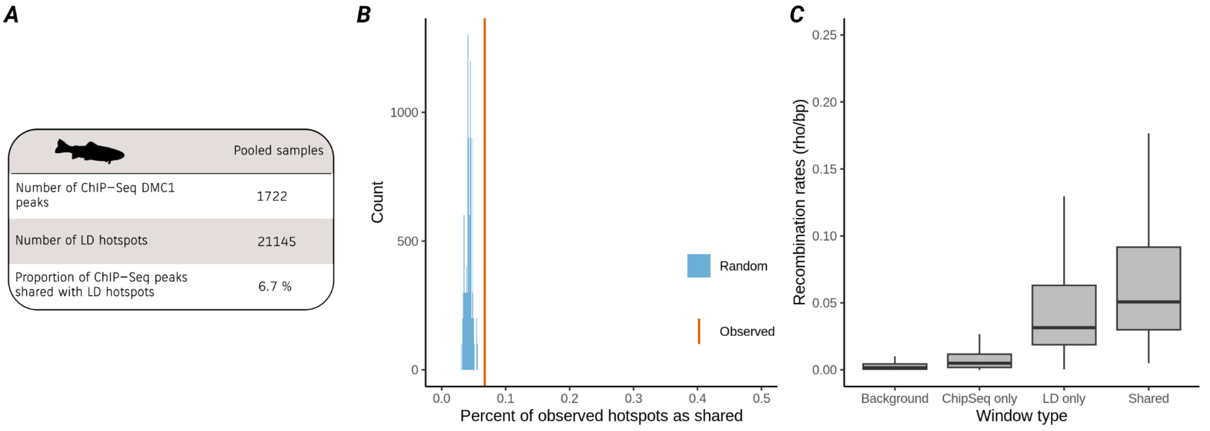


**Figure S17: Comparaison between the LD recombination landscape and the ChIP-Seq DMC1 map of the pooled samples. A)** Summary statistics. **B)** Random expectated (blue) and observed values (orange) of shared peaks between LD and ChIP-Seq maps. **C)** Recombination rates rho in the syntenic location of the ChIP-Seq peaks, in the LD-hotspots, in the shared ChIP-Seq and LD windows (*i.e.* 116 ChIP-Seq peaks shared with LD hotspots) and in the background landscapes (*i.e.* the genomic windows not containing neither a LD hotspots nor a ChIP-Seq peak).

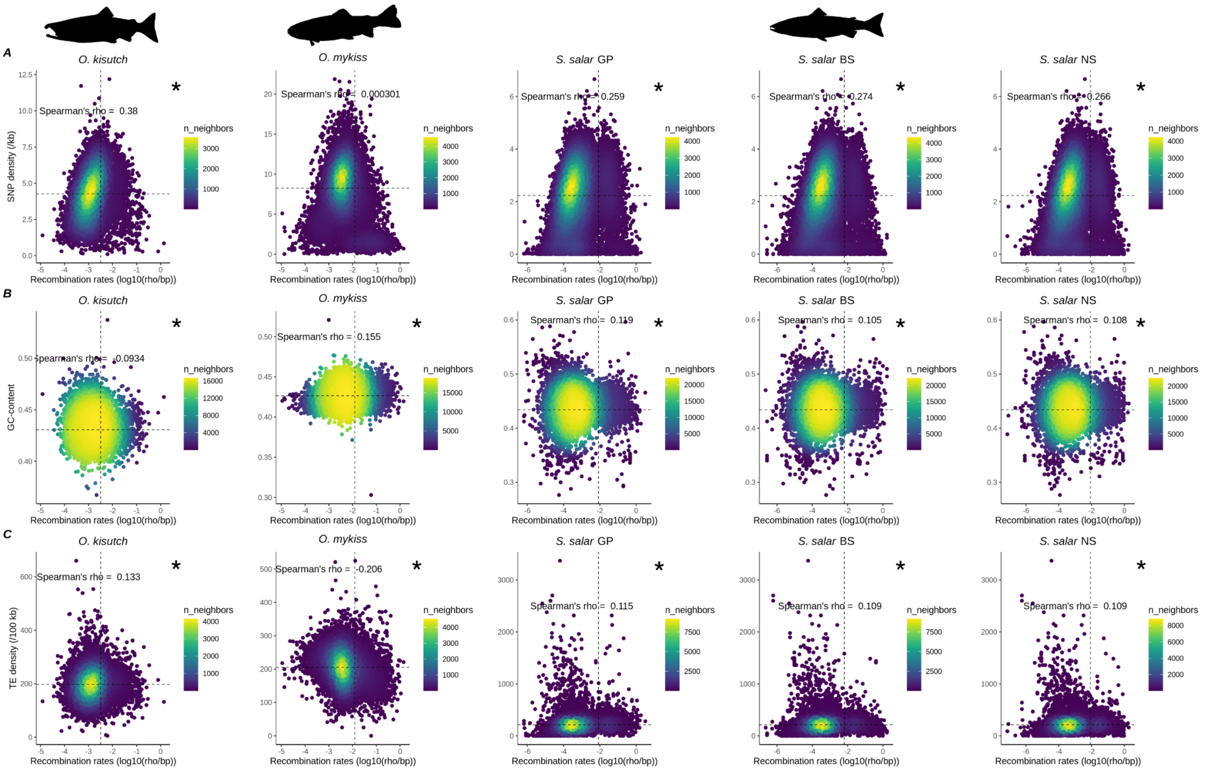


**Figure S18. Broad scale variation in genomic variables according to recombination rates. A)** SNP density (per kb). **B)** GC-content. **C)** TE density (per 100 kb). Recombination rates, SNP density GC-content and TE density were averaged in 100kb sliding windows. Significance p-value of Spearman's rank test < 0.05 are indicated by an asterisk in panels. The vertical dashed line is the mean recombination rates and the horizontal dashed line is the mean y variable.

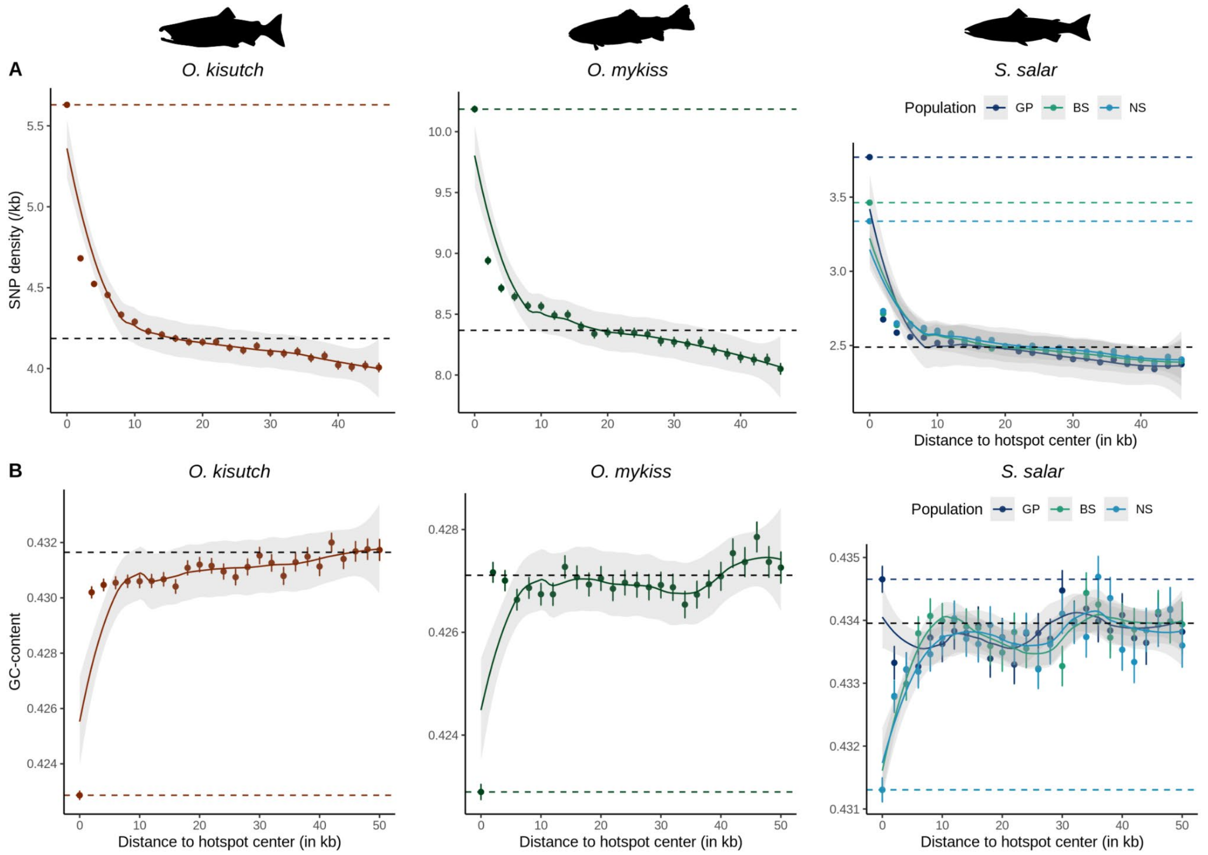


**Figure S19. Genetic diversity and base composition at recombination hotspots. A)** SNPs density (per kb), and **B)** GC content, according to distance to the nearest recombination hotspots. SNP density, GC-content and recombination rates were averaged in 2 kb windows. Colored (orange, green blue) dashed lines are mean of the y variable at hotspots of the corresponding populations, the black dashed line if the genomic mean (outside hotspots). Loess curves are shown for a span of 0.5.

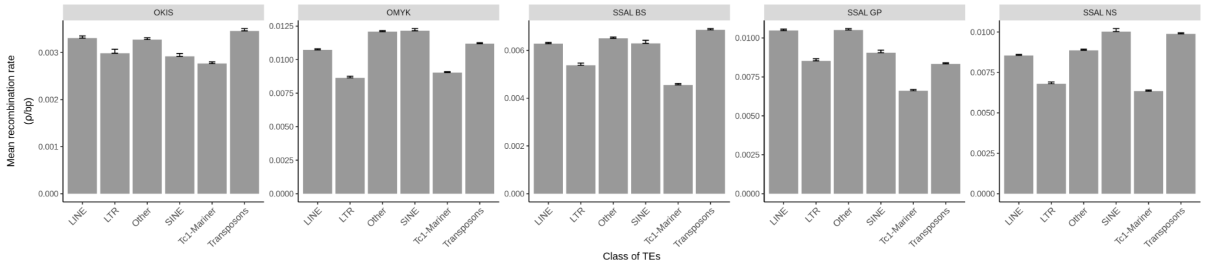


**Figure S20: Average recombination rates in TEs families.** Tc1-mariner, a family of LTR, are shown. OKIS = *O. kisutch*, OMYK = *O. mykiss*, SSAL = *S. salar* (GP, BS and NS populations), DLAB = *D. labrax*.

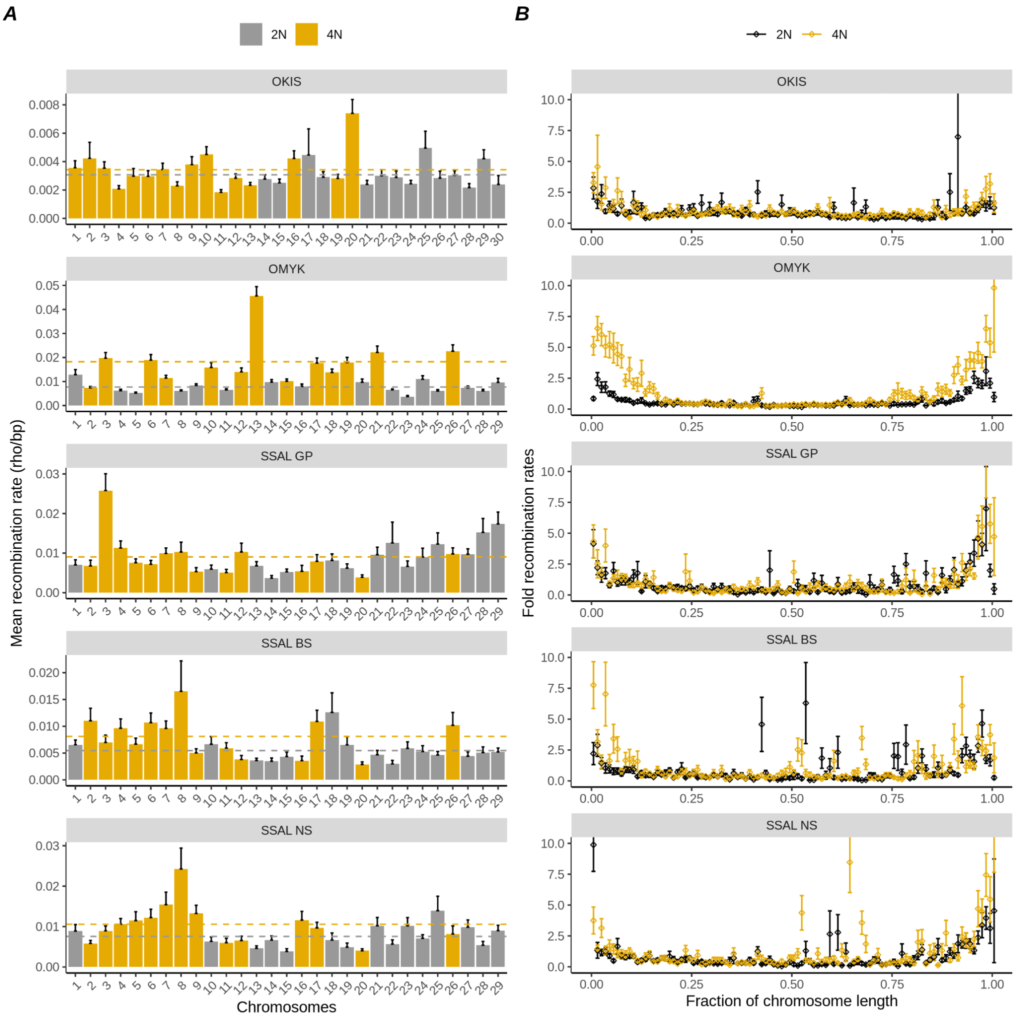


**Figure S21. Inter- and intra-chromosome variation in recombination rates in residual tetraploid chromosomes. A)** Averaged recombination rate per chromosome. Grey bars indicate chromosomes without residual tetrasomic regions (2N), and yellow bars indicate chromosomes with residual tetrasomic regions (4N) in the five Oncorhynchus and Salmo populations. Grey dashed lines represent the genome averaged recombination rates of the 2N chromosomes, and the yellow line in the 4N chromosomes. Recombination rates are significantly higher in 4N chromosomes compared to 2N chromosomes (Student test, t(13.258) =-3.9404, p < 0.05) in *O. mykiss* population, but not in *O. kisutch* (Student test, t(25.867) =-0.88786, p > 0.05) neither in *S. salar* populations (Student test, t(23.519) =-0.0026857, p > 0.05 for GP, t(20.99) = -2.2572, p < 0.05 for BS and t(19.677) =-1.9741, p = 0.06258 for NS). **B)** Recombination rates along the genome. Recombination rates were averaged into percentiles of chromosome length, and scaled by the genomic mean. Same color as panel A. OKIS = *O. kisutch*, OMYK = *O. mykiss*, SSAL = *S. salar* (GP, BS and NS populations).

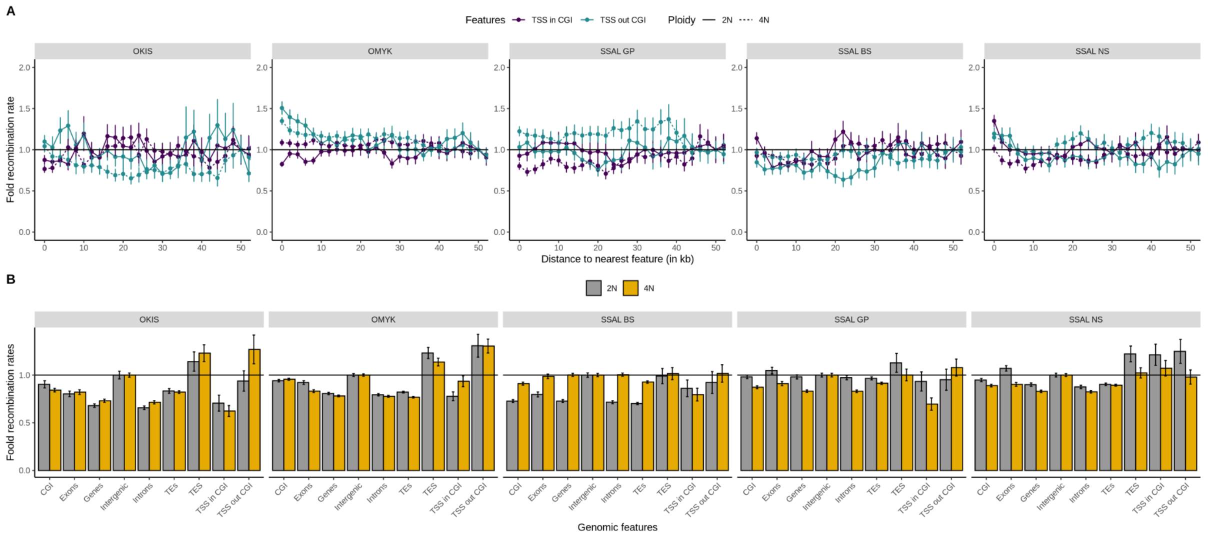


**Figure S22. Recombination rates at genomic features in residual tetraploid chromosomes. A)** Fold recombination rates (scaled by the average recombination rate at 50kb from the nearest feature) according to distance to the nearest promoter-like features (*i.e.* TSS overlapping or not a CGI). Recombination rates in chromosomes not containing residual tetraploid regions are shown by the continuous line, and by the dashed line for the 4N chromosomes. **B)** Fold recombination rates (scaled by the average recombination rates in intergenic regions) in genomic features, in 2N (grey) and 4N (yellow) chromosomes. The horizontal line shows the intergenic recombination level. TSS and TES were defined as first and last positions of genes. CGIs were mapped with EMBOSS using CpGoe > 0.6 and GC > 0. OKIS = *O. kisutch*, OMYK = *O. mykiss*, SSAL = *S. salar* (GP, BS and NS populations).

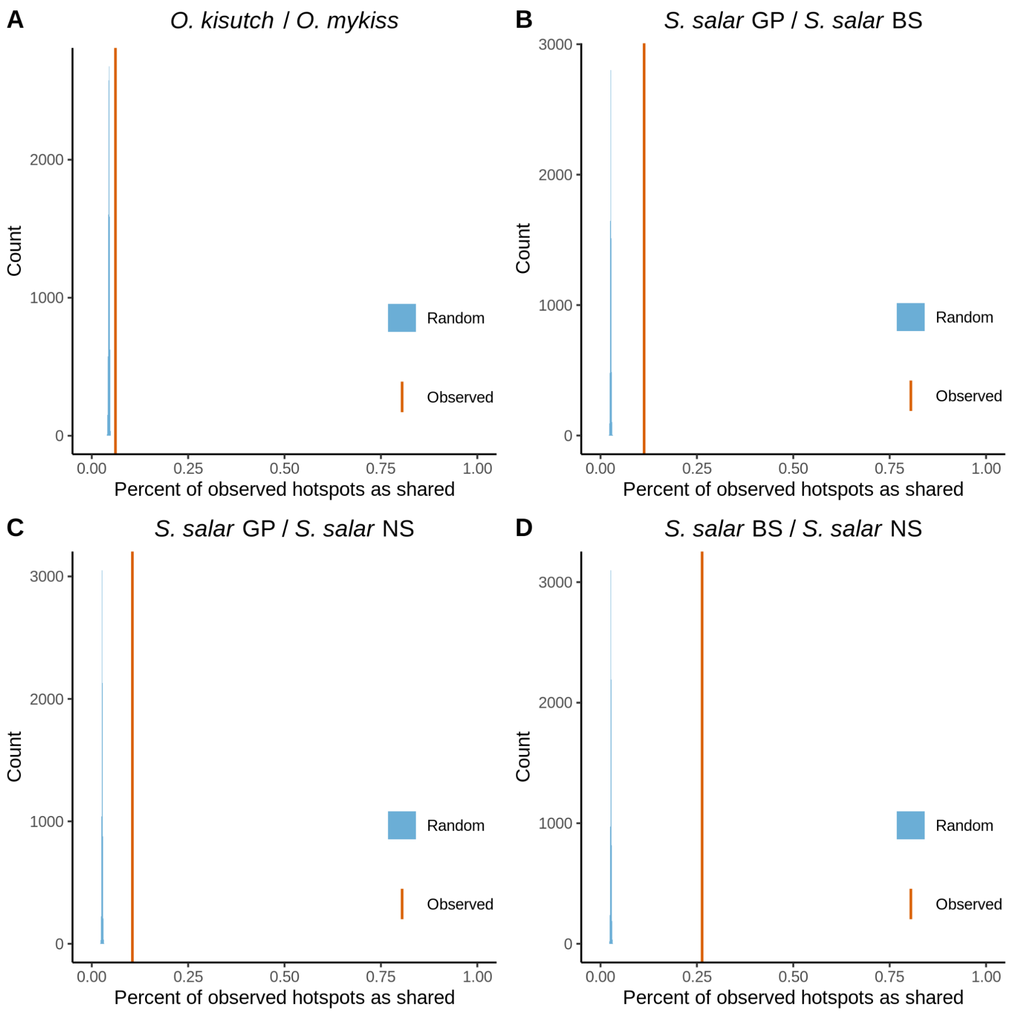


**Figure S23: Significance of hotspots sharing between closely related populations.** Random expectations (blue) and observed values (orange) of shared hotspots between **A)** *O. kisutch* and *O. mykiss*; between *S. salar* populations **B)** GP and BS; **C)** GP and NS; and between **D)** BS and NS. Shared hotspots were defined as 2-kb hotspots overlapping by at least 1 bp. Percent shared is calculated using the number of hotspots in the species/population with fewer hotspots as the denominator. The expected distribution of shared hotspots has been obtained from 1000 pairwise comparisons of random spot.

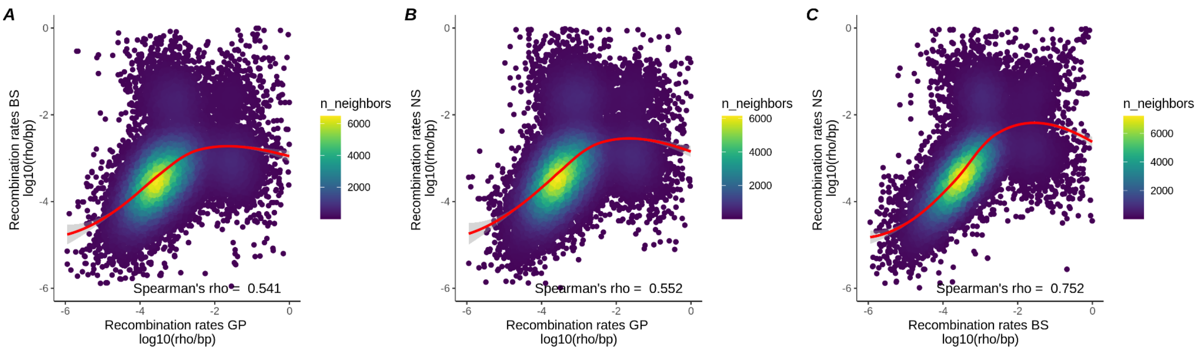


**Figure S24: Pairwise comparison of 100-kb smoothed recombination maps between *S. salar* populations. A)** Comparison between GP and BS populations. **B)** Comparison between GP and NS populations. **C)** Comparison between BS and NS populations. Spearman's rank test p-value < 0.05. Loess curves are shown for a span of 0.7.

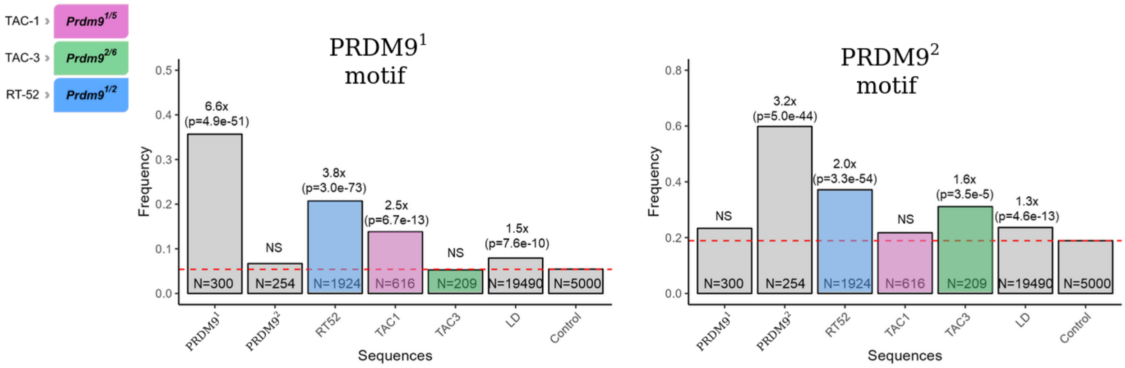


**Figure S25: DSB and LD hotspots are enriched in PRDM9 allele-specific motifs.** Frequency of sequences with at least one hit for PRDM9 allele 1 (left) and allele 2 (right) motifs at allele 1 and allele 2 sites, RT-52, TAC-1 and TAC-3 DSB hotspots, LD-hotspots and control sites. Fold enrichment relative to the control sites is shown on top of each column.

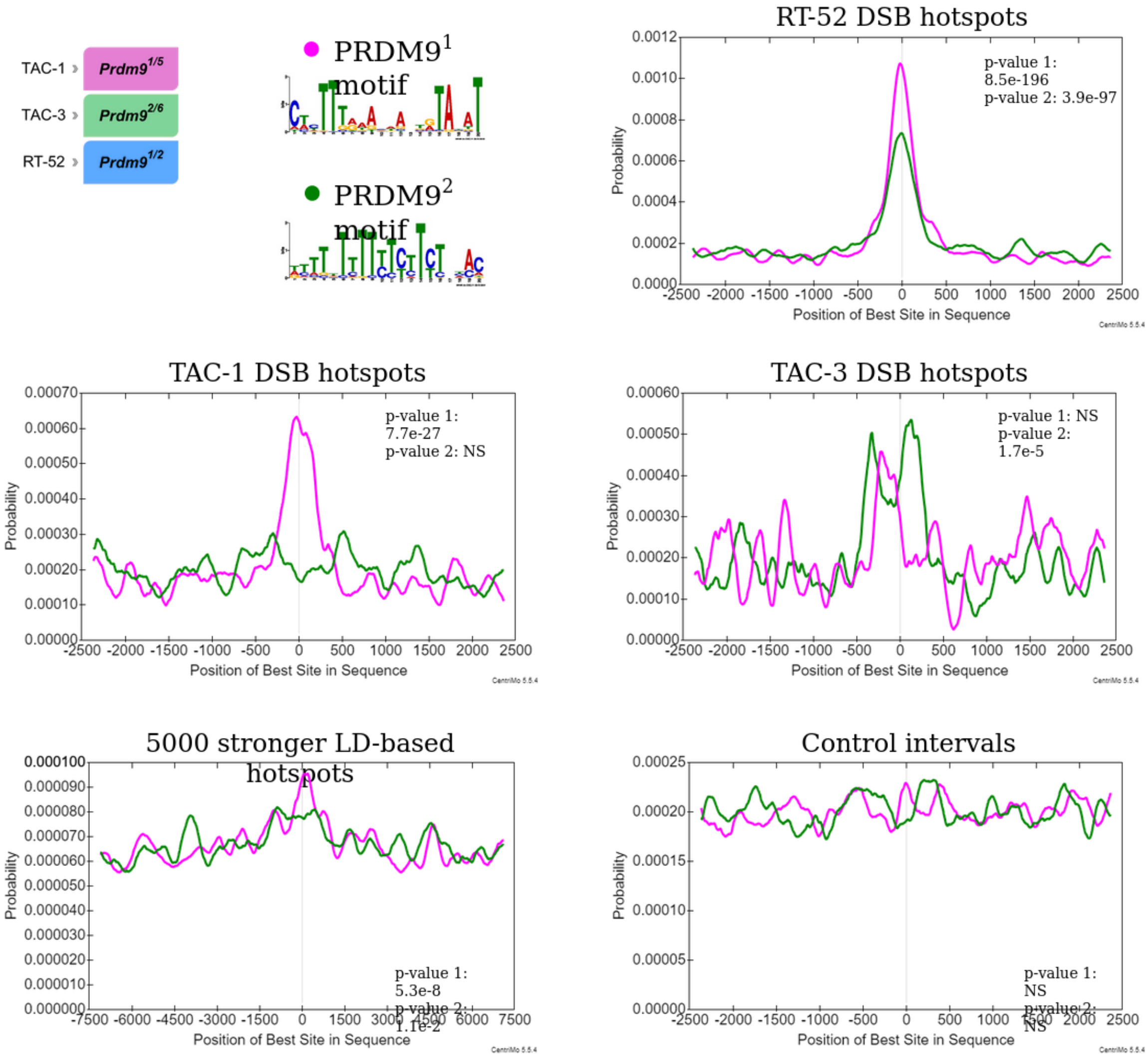


**Figure S26: DSB and LD-based hotspots are enriched in PRDM9 allele-specific motifs.** Positional distribution of hits for *Prdm9* allele 1 (pink) and allele 2 motifs (green) in RT-52, TAC-1, TAC-3 DSB hotspots, LD stronger hotspots (n=5000) and control sites (n=5000). The distribution is shown from the center of the sequence with a range of ± 2.5 kb for the DSB hotspots and the control sites. The LD hotspots were centered on the SNP interval showing the highest recombination rate (rho/bp) and the distribution extends up to 7.5 kb from the refined center. The signal is smoothed by weighted moving average and hits were calculated either in a 750 bp window for the LD hotpots and in a 250 bp window for all other sequences.

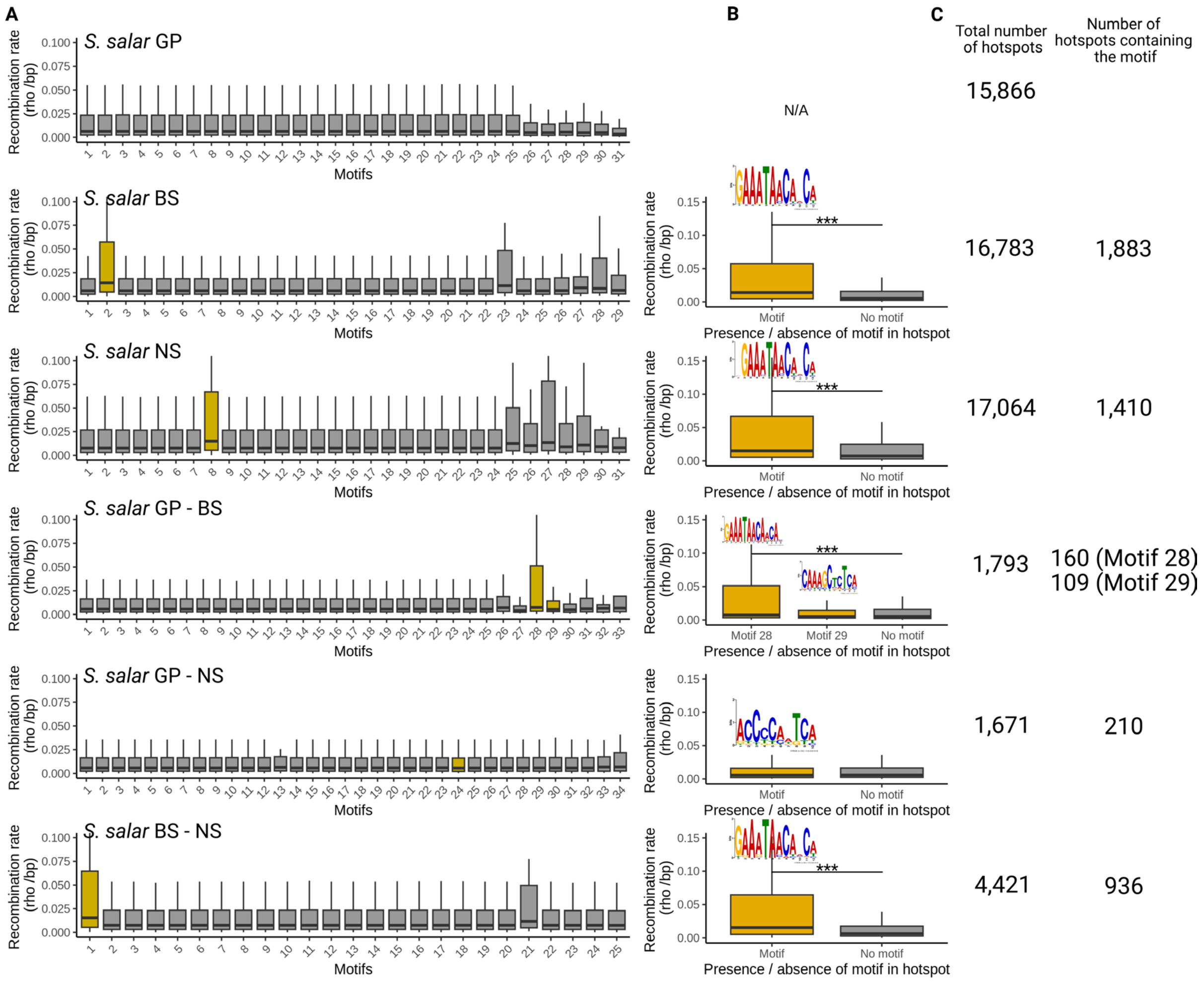


**Figure S27: Motifs enrichment at population-specific and shared recombination hotspots in *S. salar* populations. A)** Average recombination rate in motifs found enriched at hotspot. Yellow boxes show motifs found in at least 5% of hotspots showing two-fold enrichment compared to the control set of random spots. **B)** Average recombination rate in hotspots containing the retained motifs from panel A (with the corresponding motifs shown) compared to hotspots not containing the retained motifs. Significant Student's tests are indicated (***, p-value<0.05). **C)** Effective sizes (i.e. total number of population-specific and shared hotspots and number of hotspots containing the retained motifs shown in yellow).

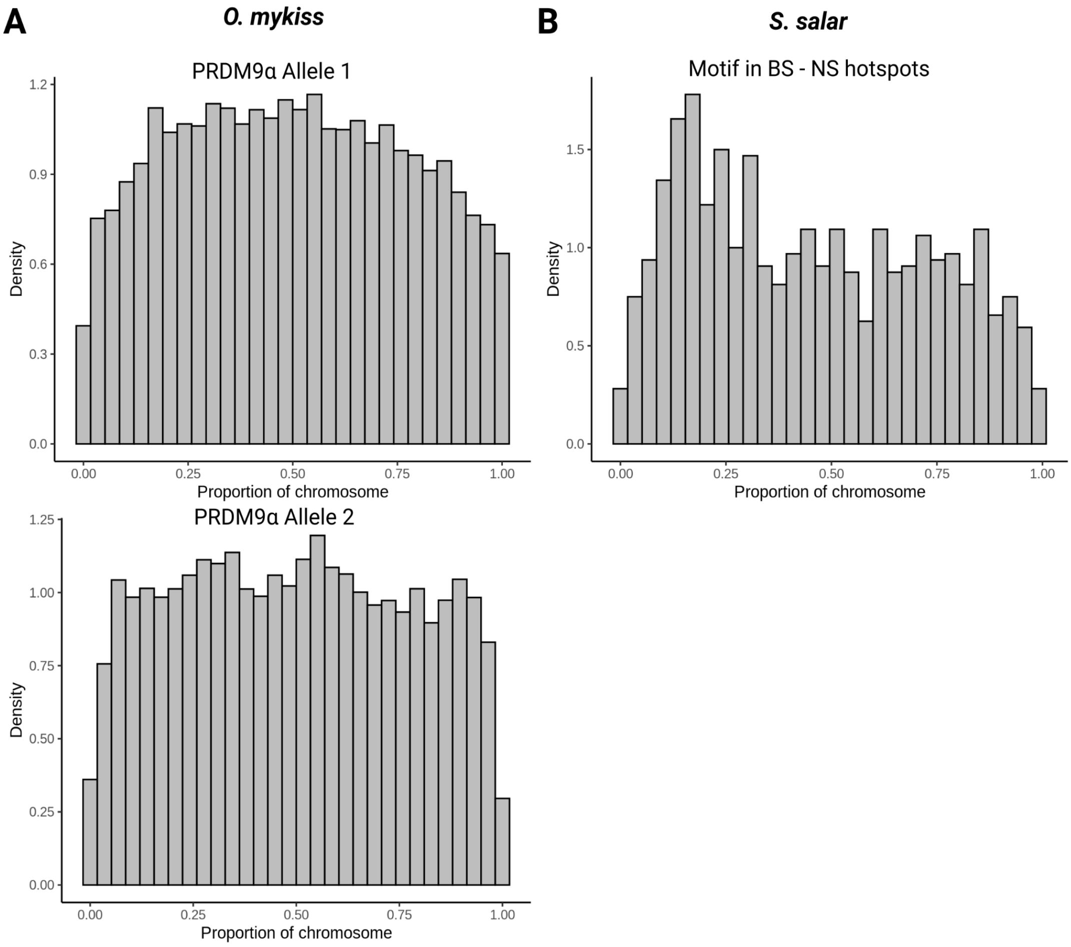


**Figure S28: Genome wide distribution of PRDM9α motifs along chromosomes in *O. mykiss*. A)** Distribution of PRDM9[1] (n=68047) and PRDM9[2] (n=59986) motifs in rainbow trout genome along chromosomes (paces of 1/30 of chromosome length). **B)** Distribution of motifs enriched in the shared hotspots between the BS and NS populations of the Atlantic salmon (n=936).

#### 4. Supplementary Tables

**Table S1: List of genotyped samples.** Details about the *Salmo salar* and *Oncorhynchus mykiss* samples used in this study and the corresponding *Prdm9α* genotypes identified.

| Sample | Sample location | PRDM9α genotypes | |
|---|---|---|---|
| ***Salmo salar*** | | ***α1.a.2*** | ***α2.2*** |
| ssa12816 | l'Oir / la Sélune river | allele 3 / allele 4 | allele 1 / allele 1 |
| ssa12805 | l'Oir / la Sélune river | allele 1 / allele 2 | allele 1 / allele 1 |
| ssa13740 | l'Oir / la Sélune river | allele 3 / allele 3 | allele 1 / allele 1 |
| ssa13649 | l'Oir / la Sélune river | allele 1 / allele 4 | allele 1 / allele 1 |
| ssa13672 | l'Oir / la Sélune river | allele 1 / allele 3 | allele 1 / allele 1 |
| ssa13651 | l'Oir / la Sélune river | allele 3 / allele 3 | allele 1 / allele 1 |
| ssa13652 | l'Oir / la Sélune river | allele 1 / allele 3 | allele 1 / allele 1 |
| ssa13653 | l'Oir / la Sélune river | allele 5 / allele 6 | allele 1 / allele 1 |
| ssa13654 | l'Oir / la Sélune river | allele 3 / allele 7 | allele 1 / allele 1 |
| ssa13655 | l'Oir / la Sélune river | allele 1 / allele 6 | allele 1 / allele 1 |
| ssa13656 | l'Oir / la Sélune river | allele 1 / allele 9 | allele 1 / allele 1 |
| ssa13657 | l'Oir / la Sélune river | allele 3 / allele 10 | allele 1 / allele 1 |
| ssa13658 | l'Oir / la Sélune river | allele 1 / allele 8 | allele 1 / allele 1 |
| ssa13659 | l'Oir / la Sélune river | allele 1 / allele 3 | allele 1 / allele 1 |
| ssa13661 | l'Oir / la Sélune river | allele 3 / allele 3 | allele 1 / allele 1 |
| ssa13662 | l'Oir / la Sélune river | allele 3 / allele 6 | allele 1 / allele 1 |
| ssa13663 | l'Oir / la Sélune river | allele 1 / allele 3 | allele 1 / allele 1 |
| ssa13664 | l'Oir / la Sélune river | allele 1 / allele 9 | allele 1 / allele 1 |
| ssa13668 | l'Oir / la Sélune river | allele 3 / allele 5 | allele 1 / allele 1 |
| ssa13669 | l'Oir / la Sélune river | allele 1 / allele 3 | allele 1 / allele 1 |
| PIT-475 | l'Oir / la Sélune river | allele 3 / allele 9 | |

| | | | |
|---|---|---|---|
| PIT-693 | l'Oir / la Sélune river | allele 3 / allele 3 | |
| PIT-948 | l'Oir / la Sélune river | allele 6 / allele 6 | |
| ssa-261 | l'Oir / la Sélune river | allele 1 / allele 11 | |
| ssa-673 | l'Oir / la Sélune river | allele 1 / allele 1 | |
| ssa-728 | l'Oir / la Sélune river | allele 3 / allele 3 | |
| ***Oncorhynchus mykiss*** | | ***α1.a.1*** | ***α2.2*** |
| rt601 | INRAE PEIMA (Sizun) | allele 1 / allele 6 | allele 1 / allele 1 |
| rt602 | INRAE PEIMA (Sizun) | allele 2 / allele 4 | allele 1 / allele 1 |
| rt603 | INRAE PEIMA (Sizun) | allele 2 / allele 6 | allele 2 / allele 2 |
| rt604 | INRAE PEIMA (Sizun) | allele 3 / allele 5 | allele 1 / allele 1 |
| rt605 | INRAE PEIMA (Sizun) | allele 1 / allele 3 | allele 1 / allele 4 |
| rt606 | INRAE PEIMA (Sizun) | allele 4 / allele 4 | allele 1 / allele 3 |
| rt607 | INRAE PEIMA (Sizun) | allele 4 / allele 6 | allele 1 / allele 5 |
| rt608 | INRAE PEIMA (Sizun) | allele 4 / allele 5 | allele 1 / allele 3 |
| rt609 | INRAE PEIMA (Sizun) | allele 4 / allele 4 | allele 1 / allele 1 |
| rt610 | INRAE PEIMA (Sizun) | allele 2 / allele 6 | allele 2 / allele 2 |
| rt611 | INRAE PEIMA (Sizun) | allele 5 / allele 6 | allele 1 / allele 3 |
| rt612 | INRAE PEIMA (Sizun) | allele 4 / allele 4 | allele 1 / allele 2 |
| rt613 | INRAE PEIMA (Sizun) | allele 4 / allele 7 | allele 1 / allele 2 |
| rt614 | INRAE PEIMA (Sizun) | allele 6 / allele 6 | allele 3 / allele 3 |
| rt615 | INRAE PEIMA (Sizun) | allele 2 / allele 6 | allele 1 / allele 1 |
| rt616 | INRAE PEIMA (Sizun) | allele 4 / allele 5 | allele 1 / allele 3 |
| rt617 | INRAE PEIMA (Sizun) | allele 4 / allele 6 | allele 1 / allele 1 |
| rt618 | INRAE PEIMA (Sizun) | allele 4 / allele 4 | allele 1 / allele 3 |
| rt619 | INRAE PEIMA (Sizun) | allele 4 / allele 4 | allele 1 / allele 1 |
| rt620 | INRAE PEIMA (Sizun) | allele 2 / allele 6 | allele 1 / allele 3 |

| | | |
|---|---|---|
| RT-52 | INRAE PEIMA (Sizun) | allele 1 / allele 2 |
| TAC-1 | INRAE PEIMA (Sizun) | allele 1 / allele 5 |
| TAC-3 | INRAE PEIMA (Sizun) | allele 2 / allele 6 |

**Table S2: List of primers.** The primers used in this study to genotype the zinc finger array of *Prdm9α* in *Salmo salar* and *Oncorhynchus mykiss*.

| Primer ID | Sequence (5'->3') | Target paralog | Use |
|---|---|---|---|
| ***Salmo salar*** | | | |
| ssa05Pa2F | TGTTGAGGAGTGGAGAGATCAGA | *α1.a.2* | PCR and sequencing |
| ssa05Pa2R | CCCGCCCAGTTAGGCTTCTA | *α1.a.2* | PCR |
| ssa05Pa3R | AACAGTTAGTTGAAAGTTCCACG | *α1.a.2* | Sequencing |
| ssa17Pa1F | TGTTCTCCTTCACGGCTCAG | *α2.2* | PCR |
| ssa17Pa2R | TGCACTGTCTTTGGGGCTGATTA | *α2.2* | PCR |
| ssa17Pa3F | GTGGCTCTCAACGATGTTCAA | *α2.2* | Sequencing |
| ssa17Pa3R | CCAGTAAATCAGTTGGTGCTA | *α2.2* | Sequencing |
| ***Oncorhynchus mykiss*** | | | |
| rt31Pa1F_new | CTCTGGCTGTCCGTTCTCCTTCACC | *α1.a.1* | PCR |
| rt31Pa2R_new | GACTGCAGTTGGTGTGGGTACTGG | *α1.a.1* | PCR |
| ssa05Pa2F | TGTTGAGGAGTGGAGAGATCAGA | *α1.a.1* | Sequencing |
| rt31Pa1R | ATACACAAGATGGACGCCAGAGT | *α1.a.1* | Sequencing |
| rt07Pa2F | GCTCTCAGCGGTGTTCAACAACT | *α2.2* | PCR and sequencing |
| rt07Pa1R | ACATGTCTCACTGCAGTCACC | *α2.2* | PCR and sequencing |

**Table S3: List of ChIP-seq experiments performed.** Details about the ChIP-seq experiment performed in *Oncorhynchus mykiss* testes. In the third column DMC1-GP or -R1 refer to the animal used to raise the antibody, respectively guinea pig and rabbit individual 1.

| Sample | Maturation stage | Antibody | Experiment | Replicate | Raw reads | Mapped reads | Accession number |
|---|---|---|---|---|---|---|---|
| RT-52 | IV | DMC1 - GP | ChIP-seq | 1 | 110M | 70M | |
| RT-52 | IV | DMC1 - R1 | ChIP-seq | 2 | 94M | 69M | |
| RT-52 | IV | None | Input | 1 | 105M | 90M | |
| TAC-1 | III b | DMC1 - R1 | ChIP-seq | 1 | 87M | 64M | |
| TAC-1 | III b | DMC1 - R1 | ChIP-seq | 2 | 100M | 78M | |
| TAC-1 | III b | H3K4me3 | ChIP-seq | 1 | 486M | 461M | |
| TAC-1 | III b | H3K4me3 | ChIP-seq | 2 | 400M | 371M | |
| TAC-1 | III b | H3K36me3 | ChIP-seq | 1 | 555M | 524M | |
| TAC-1 | III b | H3K36me3 | ChIP-seq | 2 | 394M | 370M | |
| TAC-1 | III b | None | Input | 1 | 518M | 509M | |
| TAC-3 | III a | DMC1 - R1 | ChIP-seq | 1 | 91M | 68M | |
| TAC-3 | III a | DMC1 - R1 | ChIP-seq | 2 | 67M | 51M | |
| TAC-3 | III a | H3K4me3 | ChIP-seq | 1 | 229M | 210M | |
| TAC-3 | III a | H3K4me3 | ChIP-seq | 2 | 472M | 456M | |
| TAC-3 | III a | H3K36me3 | ChIP-seq | 1 | 251M | 229M | |
| TAC-3 | III a | H3K36me3 | ChIP-seq | 2 | 449M | 433M | |
| TAC-3 | III a | None | Input | 1 | 327M | 316M | |
| TAC-3 | III a | None | Input | 2 | 536M | 528M | |

**Table S4: Sample accession number and location.** Population samples of *O. kisutch*, *O. mykiss* and *S. salar* used to build the linkage disequilibrium-based recombination landscapes were selected from (27-29).

| Sample | SRA accession number | Sample location |
|---|---|---|
| ***Oncorhynchus kisutch*** | | |
| Okis_ca_01 | SRX3161578 | Capilano River Hatchery |
| Okis_ca_02 | SRX3161579 | Capilano River Hatchery |
| Okis_ca_03 | SRX3161580 | Capilano River Hatchery |
| Okis_ca_04 | SRX3161581 | Capilano River Hatchery |
| Okis_ca_05 | SRX3161582 | Capilano River Hatchery |
| Okis_ic_01 | SRX3161545 | Inch Creek Hatchery |
| Okis_ic_02 | SRX3161544 | Inch Creek Hatchery |
| Okis_ic_03 | SRX3161543 | Inch Creek Hatchery |
| Okis_ic_04 | SRX3161547 | Inch Creek Hatchery |
| Okis_ic_05 | SRX3161546 | Inch Creek Hatchery |
| Okis_ro_01 | SRX3161552 | Robertson Creak Hatchery |
| Okis_ro_02 | SRX3161555 | Robertson Creak Hatchery |
| Okis_ro_03 | SRX3161554 | Robertson Creak Hatchery |
| Okis_ro_04 | SRX3161557 | Robertson Creak Hatchery |
| Okis_ro_05 | SRX3161556 | Robertson Creak Hatchery |
| Okis_sa_01 | SRX3161583 | Salmon River |
| Okis_sa_02 | SRX3161584 | Salmon River |
| Okis_sa_03 | SRX3161585 | Salmon River |
| Okis_sa_04 | SRX3161575 | Salmon River |
| Okis_sa_06 | SRX3161576 | Salmon River |
| **Oncorhynchus mykiss** | | |
| Omyk_dw_01 | SRX2820108 | Dworshak |
| Omyk_dw_02 | SRX2820111 | Dworshak |
| Omyk_dw_03 | SRX2820105 | Dworshak |
| Omyk_dw_04 | SRX2820113 | Dworshak |
| Omyk_el_01 | SRX2820112 | Elwha |
| Omyk_el_02 | SRX2820103 | Elwha |
| Omyk_el_03 | SRX2820102 | Elwha |
| Omyk_el_04 | SRX2820071 | Elwha |

| | | |
|---|---|---|
| Omyk_id_01 | SRX2820064 | Idaho (Big Bear River) |
| Omyk_id_02 | SRX2820079 | Idaho (Big Bear River) |
| Omyk_lq_01 | SRX2820073 | L. Quinault |
| Omyk_lq_02 | SRX2820107 | L. Quinault |
| Omyk_lq_03 | SRX2820106 | L. Quinault |
| Omyk_lq_04 | SRX2820084 | L. Quinault |
| Omyk_qu_01 | SRX2820093 | Quinault |
| Omyk_qu_02 | SRX2820085 | Quinault |
| Omyk_qu_03 | SRX2820087 | Quinault |
| Omyk_qu_04 | SRX2820104 | Quinault |
| Omyk_sk_01 | SRX2820109 | Skamania |
| Omyk_sk_02 | SRX2820098 | Skamania |
| Omyk_sk_03 | SRX2820101 | Skamania |
| Omyk_sk_04 | SRX2820072 | Skamania |
| ***Salmo Salar*** | | |
| Ssal_gp_01 | ERS4601709 | Bonaventure |
| Ssal_gp_02 | ERS4601710 | Bonaventure |
| Ssal_gp_03 | ERS4601714 | Bonaventure |
| Ssal_gp_04 | ERS4601715 | Bonaventure |
| Ssal_gp_05 | ERS4601716 | Bonaventure |
| Ssal_gp_06 | ERS4601717 | Bonaventure |
| Ssal_gp_07 | ERS4601718 | Bonaventure |
| Ssal_gp_08 | ERS4601977 | Petite riviere Cascapedia |
| Ssal_gp_09 | ERS4601978 | Petite riviere Cascapedia |
| Ssal_gp_10 | ERS4601979 | Petite riviere Cascapedia |
| Ssal_gp_11 | ERS4601980 | Petite riviere Cascapedia |
| Ssal_gp_12 | ERS4601981 | Petite riviere Cascapedia |
| Ssal_gp_13 | ERS4601983 | Petite riviere Cascapedia |
| Ssal_gp_14 | ERS4601985 | Petite riviere Cascapedia |
| Ssal_gp_15 | ERS4601986 | Petite riviere Cascapedia |
| Ssal_gp_16 | ERS4601721 | De la Chaloupe |
| Ssal_gp_17 | ERS4601723 | De la Chaloupe |
| Ssal_gp_18 | ERS4601724 | De la Chaloupe |

| | | |
|---|---|---|
| Ssal_gp_19 | ERS4601725 | De la Chaloupe |
| Ssal_gp_20 | ERS4601726 | De la Chaloupe |
| Ssal_bs_01 | ERS4601833 | Komagelva |
| Ssal_bs_02 | ERS4601834 | Komagelva |
| Ssal_bs_03 | ERS4601835 | Komagelva |
| Ssal_bs_04 | ERS4601836 | Komagelva |
| Ssal_bs_05 | ERS4601837 | Komagelva |
| Ssal_bs_06 | ERS4601839 | Komagelva |
| Ssal_bs_07 | ERS4601842 | Komagelva |
| Ssal_bs_08 | ERS4601935 | Neiden |
| Ssal_bs_09 | ERS4601936 | Neiden |
| Ssal_bs_10 | ERS4601938 | Neiden |
| Ssal_bs_11 | ERS4601939 | Neiden |
| Ssal_bs_12 | ERS4601941 | Neiden |
| Ssal_bs_13 | ERS4601942 | Neiden |
| Ssal_bs_14 | ERS4601943 | Neiden |
| Ssal_bs_15 | ERS4601944 | Neiden |
| Ssal_bs_16 | ERS4601862 | Langfjordvassdraget |
| Ssal_bs_17 | ERS4601865 | Langfjordvassdraget |
| Ssal_bs_18 | ERS4601866 | Langfjordvassdraget |
| Ssal_bs_19 | ERS4601867 | Langfjordvassdraget |
| Ssal_bs_20 | ERS4601869 | Langfjordvassdraget |
| Ssal_ns_01 | ERS4602054 | Suldalslaagen |
| Ssal_ns_02 | ERS4602056 | Suldalslaagen |
| Ssal_ns_03 | ERS4602057 | Suldalslaagen |
| Ssal_ns_04 | ERS4602058 | Suldalslaagen |
| Ssal_ns_05 | ERS4602059 | Suldalslaagen |
| Ssal_ns_06 | ERS4602060 | Suldalslaagen |
| Ssal_ns_07 | ERS4602061 | Suldalslaagen |
| Ssal_ns_08 | ERS4602062 | Suldalslaagen |
| Ssal_ns_09 | ERS4602063 | Suldalslaagen |
| Ssal_ns_10 | ERS4602090 | Vikedalselva i Vindafjord |
| Ssal_ns_11 | ERS4602091 | Vikedalselva i Vindafjord |
| Ssal_ns_12 | ERS4602092 | Vikedalselva i Vindafjord |

| | | |
|---|---|---|
| Ssal_ns_13 | ERS4602093 | Vikedalselva i Vindafjord |
| Ssal_ns_14 | ERS4602094 | Vikedalselva i Vindafjord |
| Ssal_ns_15 | ERS4602097 | Vikedalselva i Vindafjord |
| Ssal_ns_16 | ERS4602098 | Vikedalselva i Vindafjord |
| Ssal_ns_17 | ERS4602099 | Vikedalselva i Vindafjord |
| Ssal_ns_18 | ERS4601778 | Flaamselva |
| Ssal_ns_19 | ERS4601780 | Flaamselva |
| Ssal_ns_20 | ERS4601784 | Flaamselva |

**Table S5: Programme versions.** Details of the programme versions used at each step of the reconstruction of LD-based recombination landscapes in the five salmonid populations. * The *D. labrax* dataset was taken from Duranton et al. 2020, who used a reference panel of 22 genomes fully phased-by-transmission using trio-sequencing as a learning reference for the statistical phasing of 46 additional genomes with Eagle2 v2.4. Variants were oriented using whole-genome resequencing data (>20x) from the closely related species *Dicentrarchus punctatus*, which was used as an outgroup.

| Steps | Read mapping | BAM formating | Variant calling | Variant Filtering | Variant Filtering | Pre-phasing | Phasing | Variant orientation | Recombination rates estimation | TEs de novo annotation | TEs mapping | CGIs annotation |
|---|---|---|---|---|---|---|---|---|---|---|---|---|
| Programmes | bwa mem | Picard | GATK | Bcftools | VCFtools | WhatsHap | Shapeit | Est-sfs | LDhelmet | Repeat Modeler | Repeat Masker | EMBOSS |
| *O. kisutch* | v0.7.17 | v2.25.6 | v3.8-0 (Rondeau et al., 2023) | 1.9 | v 0.1.16 | 0.18 | 4.2.1 | 2.03 | 1.9 | 2.03 | 4.1.3 | 6.6.0 |
| *O. mykiss* | v0.7.17 | v2.25.6 | v4.2.2.0 | 1.9 | v 0.1.16 | 0.18 | 4.2.1 | 2.03 | 1.9 | 2.03 | 4.1.3 | 6.6.0 |
| *S. salar* GP | v0.7.17 | v2.18.29 | 4.1.8.1 | 1.9 | 0.1.17 | 1.3 | 4.2.2 | 2.04 | 1.9 | 2.03 | 4.1.3 | 6.6.0 |
| *S. salar* BS | v0.7.17 | v2.18.29 | 4.1.8.1 | 1.9 | 0.1.17 | 1.3 | 4.2.2 | 2.04 | 1.9 | 2.03 | 4.1.3 | 6.6.0 |
| *S. salar* NS | v0.7.17 | v2.18.29 | 4.1.8.1 | 1.9 | 0.1.17 | 1.3 | 4.2.2 | 2.04 | 1.9 | 2.03 | 4.1.3 | 6.6.0 |
| *D. labrax* | v0.7.5a | v1.112 | v3.3-0 (Duranton et al., 2020) | - | v0.1.11 | - | * | - | v1.10 | - | - | 6.6.0 |

**Table S6: Summary statistics of the LD-landscape reconstruction pipeline.** Information is retrieved on the reference genome sizes, population sample sizes, mapping depth statistics, variant calling statistics and effective sizes of genomic features for each population.

| | | | *S. salar* | | | |
|---|---|---|---|---|---|---|
| | ***O. kisutch*** | ***O. mykiss*** | **GP** | **BS** | **NS** | **D. labrax** |
| **Reference genome** | | | | | | |
| Genome size | 1,686,580,692 | 1,949,962,539 | | 2,499,322,922 | | 578,963,054 |
| Number of chromosomes | 30 | 29 | | 29 | | 24 |
| **Sample collection** | | | | | | |
| Number of samples | 20 | 22 | 20 | 20 | 20 | 14 |
| **Mapping statistics** | | | | | | |
| Mean depth per individual | 29.54X | 24.87X | | 9.97X | | >20X |
| Range of mean depth per individual | [23.47-32.95] | [10.97-31.27] | | [7.43-13.07] | | ~[15-40] |
| **Variant calling and filtering statistics** | | | | | | |
| Number of SNPs after variant calling | 9,590,270 | 38,601,311 | | 27,061,466 | | 14,579,961 |
| Number of SNPs after filtering | 5,133,567 | 10,797,232 | 2,829,055 | 2,700,533 | 2,521,890 | 5,074,249 |
| SNP density (per bp) after filtering | 0.0031 | 0.0055 | 0.0011 | 0.0011 | 0.0010 | 0.009 |
| **Genomic features** | | | | | | |
| Number of genes | 31,150 | 44,615 | | 52,016 | | 18,536 |
| Number of exons | 600,584 | 836,394 | | 1,405,313 | | 192,583 |
| Number of introns | 289,493 | 370,185 | | 432,062 | | 175,218 |
| Number of TSS | 31,137 | 44,615 | | 52,015 | | 18,191 |
| Number of TES | 31,132 | 44,588 | | 51,988 | | 18,141 |
| Number of CGIs | 466,523 | 684,137 | | 648,505 | | 131,512 |
| Percent of TSS in CGIs | 60% | 58% | | 57% | | 38% |
| Percent of TEs elements | 47.37% | 48.56% | | 52.26% | | *NA* |
| Total length of interspersed repeats (in bp) | 1,122,660,520 | 1,058,078,392 | | 1,440,595,994 | | *NA* |
| % Tc1-mariner | 13.16% | 14.7% | | 12.48% | | *NA* |
| Number of Tc1 mariner | 799,417 | 748,980 | | 782,170 | | *NA* |

**Table S7: Chromosome location of the retained PRDM9 paralog copies.** The index allow to identify the corresponding copy in the phylogeny of the α paralog copies in **S4 Fig** and the β copies in **S3 Fig**. We retrieved the location of the regions covering the three domains KRAB, SSXRD and SET obtained from the blast analysis. The start and end positions correspond to the start position of the first exon blasted and the end position of the last exon blasted.

| Species | Index | Reference genome RefSeq accession number | Chromosome RefSeq accession number | Start position | End position |
|---|---|---|---|---|---|
| **Prdm9 α paralogs** | | | | | |
| *O. kisutch* | 1 | GCF_002021735.2 | NC_034188.2 | 12887053 | 12894097 |
| | 2 | | NC_034192.2 | 56290129 | 56303034 |
| | 3 | | NC_034178.2 | 5473664 | 5478378 |
| | 4 | | NC_034188.2 | 13227731 | 13233628 |
| | 5 | | NC_034192.2 | 56235978 | 56236786 |
| *O. mykiss* | 6 | GCF_002163495.1 | NC_035101.1 | 8415083 | 8423079 |
| | 7 | | NC_035090.1 | 72512843 | 72525023 |
| | 8 | | NC_035083.1 | 9752825 | 9756862 |
| | 9 | | NC_035101.1 | 8732180 | 8735814 |
| *S. salar* | 10 | GCF_905237065.1 | NC_059450.1 | 87343648 | 87355556 |
| | 11 | | NC_059446.1 | 12774207 | 12794545 |
| | 12 | | NC_059458.1 | 19507805 | 19530082 |
| | 13 | | NC_059457.1 | 72831017 | 72843243 |
| | 14 | | NC_059446.1 | 12827488 | 12840869 |
| | 15 | | NC_059450.1 | 87687617 | 87688156 |
| *O. nerka* | 16 | GCF_006149115.2 | NW_021798387.1 | 5360 | 13922 |
| | 17 | | NW_021812881.1 | 27422 | 38545 |
| | 18 | | NW_021787950.1 | 49862 | 52085 |
| | 19 | | NW_021810464.1 | 37685 | 38512 |
| | 20 | | NW_021792348.1 | 114852 | 115615 |
| *O. keta* | 21 | GCF_023373465.1 | NC_068450.1 | 14332662 | 14340913 |
| | 22 | | NC_068424.1 | 74859328 | 74871640 |
| | 23 | | NC_068457.1 | 17803818 | 17834627 |
| | 24 | | NW_026283388.1 | 13799 | 14605 |
| | 25 | | NC_068450.1 | 14795938 | 14799988 |

| | | | | | |
|---|---|---|---|---|---|
| O. tshawytscha | 26 | GCF_018296145.1 | NC_056436.1 | 78654218 | 78662610 |
| | 27 | | NC_056449.1 | 33170582 | 33187287 |
| | 28 | | NW_024609832.1 | 379135 | 382678 |
| | 29 | | NC_056449.1 | 33127116 | 33138001 |
| | 30 | | NC_056436.1 | 78152338 | 78156301 |
| O. gorbuscha | 31 | GCF_021184085.1 | NC_060195.1 | 13395381 | 13404061 |
| | 32 | | NC_060185.1 | 88927636 | 88981659 |
| | 33 | | NC_060193.1 | 53131373 | 53143551 |
| | 34 | | NC_060195.1 | 13858179 | 13860498 |
| | 35 | | NC_060184.1 | 84369277 | 84370834 |
| S. trutta | 36 | GCA_901001165.2 | LR584428.1 | 31835604 | 31845878 |
| | 37 | | LR584416.1 | 9894917 | 9917525 |
| | 38 | | LR584413.1 | 12304219 | 12315348 |
| | 39 | | LR584406.1 | 51081287 | 51092978 |
| | 40 | | LR584416.1 | 9946574 | 9958847 |
| | 41 | | CAAJIE020000684.1 | 38776 | 41222 |
| S. namaycush | 42 | GCF_016432855.1 | NC_052323.1 | 34467596 | 34477759 |
| | 43 | | NC_052348.1 | 18528766 | 18529803 |
| | 44 | | NC_052313.1 | 49372386 | 49380629 |
| | 45 | | NC_052346.1 | 12719793 | 13806471 |
| | 46 | | NC_052323.1 | 20296993 | 20315388 |
| C. clupeaformis | 47 | GCF_020615455.1 | NC_059212.1 | 8991317 | 9007109 |
| | 48 | | NC_059226.1 | 40101271 | 40117907 |
| | 49 | | NW_025534320.1 | 170337 | 176421 |
| | 50 | | NW_025535102.1 | 51554 | 67751 |
| | 51 | | NW_025537026.1 | 17152 | 33482 |
| | 52 | | NC_059194.1 | 2849538 | 2871608 |
| | 53 | | NC_059194.1 | 2824237 | 2832607 |
| | 54 | | NW_025537642.1 | 2376 | 21348 |
| | 55 | | NW_025533470.1 | 712759 | 713888 |
| | 56 | | NW_025533470.1 | 810220 | 811146 |

| | | | | | |
|---|---|---|---|---|---|
| | 57 | | NW_025533877.1 | 108337 | 114502 |
| | 58 | | NW_025534294.1 | 39024 | 42742 |
| *T. thymallus* | 59 | GCA_023634145.1 | CM042384.1 | 27241336 | 27251246 |
| | 60 | | CM042385.1 | 6555373 | 6788464 |
| | 61 | | CM042384.1 | 27035181 | 27035496 |
| *H. hucho* | 62 | GCA_003317085.1 | QNTS01002334.1 | 29710 | 39792 |
| | 63 | | QNTS01002761.1 | 75215 | 83155 |
| | 64 | | QNTS01003609.1 | 55551 | 56973 |
| | 65 | | QNTS01000290.1 | 78088 | 82204 |
| | 66 | | QNTS01015822.1 | 8019 | 8541 |
| *E. lucius* | 67 | GCF_011004845.1 | NC_047593.1 | 21477662 | 21488947 |
| | 68 | | NC_047587.1 | 2136731 | 2399395 |
| **Prdm9 β paralogs** | | | | | |
| *O. kisutch* | 1 | GCF_002021735.2 | NC_034198.2 | 25039623 | 25040671 |
| | 2 | | NC_034184.2 | 7538438 | 7540180 |
| *O. mykiss* | 3 | GCF_002163495.1 | NC_035096.1 | 23688475 | 23689502 |
| | 4 | | NC_035099.1 | 42290320 | 42293196 |
| *S. salar* | 5 | GCF_905237065.1 | NC_059469.1 | 18949696 | 18950780 |
| | 6 | | NC_059442.1 | 94679893 | 94684082 |
| *O. nerka* | 7 | GCF_006149115.2 | NC_042550.1 | 22686397 | 22687439 |
| | 8 | | NW_021789518.1 | 42389 | 44985 |
| *O. keta* | 9 | GCF_023373465.1 | NC_068444.1 | 25615799 | 25616838 |
| | 10 | | NC_068422.1 | 7947206 | 7950309 |
| *O. tshawytscha* | 11 | GCF_018296145.1 | NC_056453.1 | 25860865 | 25861913 |
| | 12 | | NC_056429.1 | 88507466 | 88510273 |
| *O. gorbuscha* | 13 | GCF_021184085.1 | NC_060179.1 | 66056844 | 66057889 |
| | 14 | | NC_060178.1 | 98157196 | 98161663 |
| *S. trutta* | 15 | GCA_901001165.2 | LR584419.1 | 25732352 | 25733424 |
| | 16 | | LR584435.1 | 14382999 | 14387213 |
| *S. namaycush* | 17 | GCF_016432855.1 | NC_052341.1 | 15666464 | 15667511 |
| | 18 | | NC_052310.1 | 78263958 | 78266635 |
| *C. clupeaformis* | 19 | | NC_059222.1 | 13680015 | 13681061 |

| | | | | | |
|---|---|---|---|---|---|
| | 20 | GCF_020615455.1 | NC_059192.1 | 95051878 | 95052654 |
| *T. thymallus* | 21 | GCA_023634145.1 | CM042387.1 | 22808132 | 22809233 |
| | 22 | | CM042388.1 | 11440396 | 11441186 |
| *H. hucho* | 23 | GCA_003317085.1 | QNTS01001395.1 | 107566 | 108622 |
| | 24 | | QNTS01001493.1 | 122445 | 127178 |
| *E. lucius* | 25 | GCF_011004845.1 | NC_047573.1 | 20413747 | 20414822 |

**Table S8: Fine scale variations in recombination rates and raw recombination hotspots.** Summary statistics of the variations in recombination rates smoothed in 2-kb sliding windows, and of recombination hotspots retrieved from the inter-SNP recombination landscapes. The raw hotspots were defined as the consecutives inter-SNP windows with a recombination rate five-fold higher than the 50-kb flanking regions.

| | | | *S. salar* | | | |
|---|---|---|---|---|---|---|
| | ***O. kisutch*** | ***O. mykiss*** | **GP** | **BS** | **NS** | ***D. labrax*** |
| **Genome-wide recombination rates** | | | | | | |
| Genome-wide recombination rate (ρ/bp) | 0,0032 | 0.0123 | 0.0084 | 0.0065 | 0.0085 | 0.039 |
| Variation range of recombination rates (smoothed at 2 kb) | [4.49x10-7; 7.6] | [3.83x10-7; 5.07] | [2.48x10-7; 7.86] | [6.47x10-8; 5.82] | [7.33x10-8; 4.23] | [3.26x10-7; 5.68] |
| Percent of recombination in 20% of the genome | 90.1 % | 89.1 % | 98.3 % | 98 % | 98.1 % | 84.6 % |
| **Raw recombination hotspots** | | | | | | |
| Number of hotspots | 49742 | 90696 | 25139 | 25973 | 26550 | 7897 |
| Mean fold recombination rate in hotspots | 19 | 19.2 | 19.6 | 19.8 | 20.5 | 18.9 |
| Mean hotspots size (in bp) | 418 | 309 | 927 | 980 | 1073 | 1995 |
| Fraction of recombination in hotspots | 24% | 17% | 34% | 37% | 39% | 3% |
| Proportion of hotspots shorter than 2 kb | 93,5 % | 97,5 % | 80,1 % | 76 % | 73,1 % | 63.7% |

**Table S9: Overlaps between DSB hotspots and TSS/TES regions.** Overlaps were assessed for 400bp-wide windows centered on DSB hotspot centers and TSS/TES regions, defined as sequences found within 1 kb of distance from the transcription start/end site. The expected overlaps were estimated as the chance for 2 kb windows of overlapping 400bp DSB hotspots genome-wide.

| Sample | TSS category | # DSB HS | # TSS | Overlap. DSB HS | % of overlap. DSB HS | Exp. Overlap. DSB HS | Exp. % of overlap. DSB HS | p-value (chi2) |
|---|---|---|---|---|---|---|---|---|
| **DSB hotspots and TSS** | | | | | | | | |
| RT-52 | all genes | 1924 | 72603 | 96 | **5.0%** | 146 | 7.6% | 1.7E-05 |
| TAC-1 | all genes | 616 | 72603 | 28 | **4.5%** | 47 | 7.6% | 0.0039 |
| TAC-3 | all genes | 209 | 72603 | 11 | **5.3%** | 16 | 7.6% | 0.19 |
| RT-52 | protein coding genes | 1924 | 41890 | 89 | **4.6%** | 84 | 4.4% | 0.58 |
| TAC-1 | protein coding genes | 616 | 41890 | 24 | **3.9%** | 27 | 4.4% | 0.55 |
| TAC-3 | protein coding genes | 209 | 41890 | 7 | **3.3%** | 9 | 4.4% | 0.50 |
| RT-52 | non-coding genes | 1924 | 30713 | 7 | **0.4%** | 62 | 3.2% | 1.2E-12 |
| TAC-1 | non-coding genes | 616 | 30714 | 4 | **0.6%** | 20 | 3.2% | 0.00028 |
| TAC-3 | non-coding genes | 209 | 30715 | 4 | **1.9%** | 7 | 3.2% | 0.25 |
| **DSB hotspots and TES** | | | | | | | | |
| RT-52 | all genes | 1924 | 72589 | 164 | **8.5%** | 146 | 7.6% | 0.12 |
| TAC-1 | all genes | 616 | 72589 | 45 | **7.3%** | 47 | 7.6% | 0.76 |
| TAC-3 | all genes | 209 | 72589 | 22 | **10.5%** | 16 | 7.6% | 0.12 |
| RT-52 | protein coding genes | 1924 | 41886 | 150 | **7.8%** | 84 | 4.4% | 1.79E-13 |
| TAC-1 | protein coding genes | 616 | 41886 | 39 | **6.3%** | 27 | 4.4% | 0.018 |
| TAC-3 | protein coding genes | 209 | 41886 | 15 | **7.2%** | 9 | 4.4% | 0.041 |
| RT-52 | non-coding genes | 1924 | 30703 | 17 | **0.9%** | 62 | 3.2% | 6.27E-09 |
| TAC-1 | non-coding genes | 616 | 30703 | 7 | **1.1%** | 20 | 3.2% | 0.0031 |
| TAC-3 | non-coding genes | 209 | 30703 | 7 | **3.3%** | 7 | 3.2% | 1 |

**Table S10: Mean sequence identity score obtained from the blast search of the 100-kb flanking sequences of the variants in the ingroup species against the reference genome of each outgroup.** Outgroups 1, 2 and 3 for *O. kisutch* were *O. tshawytscha*, *O. nerka* and *O. mykiss* respectively; *O. tschawytscha*, *O. nerka* and *O. kisutch* for *O. mykiss*; and *Salmo trutta*, *Salvelinus alpinus* and *O. mykiss* for *S. salar*.

| | Outgroup 1 | Outgroup 2 | Outgroup 3 |
|---|---|---|---|
| *O. kisutch* | 96.53 | 95.71 | 95.48 |
| *O. mykiss* | 95.76 | 95.64 | 95.62 |
| *S. salar* | 96.95 | 93.95 | 93.31 |

#### 5. Supplementary references


1. Altschul SF, Gish W, Miller W, Myers EW, Lipman DJ. Basic local alignment search tool. J Mol Biol. 1990;215(3):403-10.
2. Ranwez V, Douzery EJP, Cambon C, Chantret N, Delsuc F. MACSE v2: Toolkit for the Alignment of Coding Sequences Accounting for Frameshifts and Stop Codons. Mol Biol Evol. 2018;35(10):2582-4.
3. Borowiec ML. AMAS: a fast tool for alignment manipulation and computing of summary statistics. PeerJ. 2016;4:e1660.
4. Christoffels A, Koh EG, Chia JM, Brenner S, Aparicio S, Venkatesh B. Fugu genome analysis provides evidence for a whole-genome duplication early during the evolution of ray-finned fishes. Mol Biol Evol. 2004;21(6):1146-51.
5. Macqueen DJ, Johnston IA. A well-constrained estimate for the timing of the salmonid whole genome duplication reveals major decoupling from species diversification. Proc Biol Sci. 2014;281(1778):20132881.
6. Vandepoele K, De Vos W, Taylor JS, Meyer A, Van de Peer Y. Major events in the genome evolution of vertebrates: paranome age and size differ considerably between ray-finned fishes and land vertebrates. Proc Natl Acad Sci U S A. 2004;101(6):1638-43.
7. Nguyen LT, Schmidt HA, von Haeseler A, Minh BQ. IQ-TREE: a fast and effective stochastic algorithm for estimating maximum-likelihood phylogenies. Mol Biol Evol. 2015;32(1):268-74.
8. Hayashi K, Yoshida K, Matsui Y. A histone H3 methyltransferase controls epigenetic events required for meiotic prophase. Nature. 2005;438(7066):374-8.
9. Birney E, Clamp M, Durbin R. GeneWise and Genomewise. Genome Res. 2004;14(5):988-95.
10. Baker Z, Schumer M, Haba Y, Bashkirova L, Holland C, Rosenthal GG, et al. Repeated losses of PRDM9-directed recombination despite the conservation of PRDM9 across vertebrates. eLife. 2017;6.
11. Cavassim MIA, Baker Z, Hoge C, Schierup MH, Schumer M, Przeworski M. PRDM9 losses in vertebrates are coupled to those of paralogs ZCWPW1 and ZCWPW2. Proc Natl Acad Sci U S A. 2022;119(9).
12. Persikov AV, Osada R, Singh M. Predicting DNA recognition by Cys2His2 zinc finger proteins. Bioinformatics. 2009;25(1):22-9.
13. Persikov AV, Singh M. De novo prediction of DNA-binding specificities for Cys2His2 zinc finger proteins. Nucleic Acids Res. 2014;42(1):97-108.
14. Billard R, Solari A, Escaffre AM. [Method for the quantitative analysis of spermatogenesis in teleost fish]. Ann Biol Anim Biochim Biophys. 1974;14(1):87-104.
15. Diagouraga B, Clement JAJ, Duret L, Kadlec J, de Massy B, Baudat F. PRDM9 Methyltransferase Activity Is Essential for Meiotic DNA Double-Strand Break Formation at Its Binding Sites. Mol Cell. 2018;69(5):853-65 e6.
16. Tardat M, Brustel J, Kirsh O, Lefevbre C, Callanan M, Sardet C, et al. The histone H4 Lys 20 methyltransferase PR-Set7 regulates replication origins in mammalian cells. Nat Cell Biol. 2010;12(11):1086-93.
17. Brick K, Pratto F, Sun CY, Camerini-Otero RD, Petukhova G. Analysis of Meiotic Double-Strand Break Initiation in Mammals. Methods Enzymol. 2018;601:391-418.
18. Khil PP, Smagulova F, Brick KM, Camerini-Otero RD, Petukhova GV. Sensitive mapping of recombination hotspots using sequencing-based detection of ssDNA. Genome Res. 2012;22(5):957-65.
19. Ewels PA, Peltzer A, Fillinger S, Patel H, Alneberg J, Wilm A, et al. The nf-core framework for community-curated bioinformatics pipelines. Nat Biotechnol. 2020;38(3):276-8.
20. Ramírez F, Ryan DP, Grüning B, Bhardwaj V, Kilpert F, Richter AS, et al. deepTools2: a next generation web server for deep-sequencing data analysis. Nucleic Acids Res. 2016;44(W1):W160-5.

21. Quinlan AR, Hall IM. BEDTools: a flexible suite of utilities for comparing genomic features. Bioinformatics. 2010;26(6):841-2.
22. Auffret P, de Massy B, Clement JAJ. Mapping Meiotic DNA Breaks: Two Fully-Automated Pipelines to Analyze Single-Strand DNA Sequencing Data, hotSSDS and hotSSDS-extra. Methods Mol Biol. 2024;2770:227-61.
23. Li Q, Brown JB, Huang H, Bickel PJ. Measuring reproducibility of high-throughput experiments. The Annals of Applied Statistics. 2011;5(3):1752-79, 28.
24. Lawrence M, Huber W, Pagès H, Aboyoun P, Carlson M, Gentleman R, et al. Software for computing and annotating genomic ranges. PLoS Comput Biol. 2013;9(8):e1003118.
25. Brick K, Smagulova F, Khil P, Camerini-Otero RD, Petukhova GV. Genetic recombination is directed away from functional genomic elements in mice. Nature. 2012;485(7400):642-5.
26. Raynaud M, Gagnaire P-A, Galtier N. Performance and limitations of linkage-disequilibrium-based methods for inferring the genomic landscape of recombination and detecting hotspots: a simulation study. Peer Community Journal. 2023;3.
27. Rondeau EB, Christensen KA, Minkley DR, Leong JS, Chan MTT, Despins CA, et al. Population-size history inferences from the coho salmon (Oncorhynchus kisutch) genome. G3 (Bethesda). 2023;13(4).
28. Gao G, Nome T, Pearse DE, Moen T, Naish KA, Thorgaard GH, et al. A New Single Nucleotide Polymorphism Database for Rainbow Trout Generated Through Whole Genome Resequencing. Front Genet. 2018;9:147.
29. Bertolotti AC, Layer RM, Gundappa MK, Gallagher MD, Pehlivanoglu E, Nome T, et al. The structural variation landscape in 492 Atlantic salmon genomes. Nat Commun. 2020;11(1):5176.
30. McKenna A, Hanna M, Banks E, Sivachenko A, Cibulskis K, Kernytsky A, et al. The Genome Analysis Toolkit: a MapReduce framework for analyzing next-generation DNA sequencing data. Genome Res. 2010;20(9):1297-303.
31. Van der Auwera GA, Carneiro MO, Hartl C, Poplin R, Del Angel G, Levy-Moonshine A, et al. From FastQ data to high confidence variant calls: the Genome Analysis Toolkit best practices pipeline. Current protocols in bioinformatics. 2013;43(1110):11.0.1-.0.33.
32. Li H, Durbin R. Fast and accurate short read alignment with Burrows-Wheeler transform. Bioinformatics. 2009;25(14):1754-60.
33. Danecek P, Auton A, Abecasis G, Albers CA, Banks E, DePristo MA, et al. The variant call format and VCFtools. Bioinformatics. 2011;27(15):2156-8.
34. Martin M, Patterson M, Garg S, Fischer SO, Pisanti N, Klau GW, et al. WhatsHap: fast and accurate read-based phasing. bioRxiv. 2016:085050.
35. Delaneau O, Zagury JF, Robinson MR, Marchini JL, Dermitzakis ET. Accurate, scalable and integrative haplotype estimation. Nat Commun. 2019;10(1):5436.
36. Stapley J, Feulner PGD, Johnston SE, Santure AW, Smadja CM. Variation in recombination frequency and distribution across eukaryotes: patterns and processes. Philos Trans R Soc Lond B Biol Sci. 2017;372(1736).
37. Keightley PD, Jackson BC. Inferring the Probability of the Derived vs. the Ancestral Allelic State at a Polymorphic Site. Genetics. 2018;209(3):897-906.
38. Crespi BJ, Teo R. Comparative phylogenetic analysis of the evolution of semelparity and life history in salmonid fishes. Evolution. 2002;56(5):1008-20.
39. Chan AH, Jenkins PA, Song YS. Genome-Wide Fine-Scale Recombination Rate Variation in Drosophila melanogaster. PLoS Genet. 2012;8(12):e1003090.
40. Duranton M, Allal F, Valière S, Bouchez O, Bonhomme F, Gagnaire PA. The contribution of ancient admixture to reproductive isolation between European sea bass lineages. Evol Lett. 2020;4(3):226-42.
41. Flynn JM, Hubley R, Goubert C, Rosen J, Clark AG, Feschotte C, et al. RepeatModeler2 for automated genomic discovery of transposable element families. Proc Natl Acad Sci U S A. 2020;117(17):9451-7.
42. Larsen F, Gundersen G, Lopez R, Prydz H. CpG islands as gene markers in the human genome. Genomics. 1992;13(4):1095-107.

43. Cross S, Kovarik P, Schmidtke J, Bird A. Non-methylated islands in fish genomes are GC-poor. Nucleic Acids Res. 1991;19(7):1469-74.
44. Long HK, Sims D, Heger A, Blackledge NP, Kutter C, Wright ML, et al. Epigenetic conservation at gene regulatory elements revealed by non-methylated DNA profiling in seven vertebrates. eLife. 2013;2:e00348.
45. Bailey TL, Johnson J, Grant CE, Noble WS. The MEME Suite. Nucleic Acids Res. 2015;43(W1):W39-49.
46. Grant CE, Bailey TL, Noble WS. FIMO: scanning for occurrences of a given motif. Bioinformatics. 2011;27(7):1017-8.
47. Bailey TL. STREME: accurate and versatile sequence motif discovery. Bioinformatics. 2021;37(18):2834-40.
48. Minkin I, Medvedev P. Scalable multiple whole-genome alignment and locally collinear block construction with SibeliaZ. Nat Commun. 2020;11(1):6327.
49. de Mendoza A, Lister R, Bogdanovic O. Evolution of DNA Methylome Diversity in Eukaryotes. J Mol Biol. 2020;432(6):1687-705.
50. Tweedie S, Charlton J, Clark V, Bird A. Methylation of genomes and genes at the invertebrate-vertebrate boundary. Mol Cell Biol. 1997;17(3):1469-75.
51. Bird AP. CpG-rich islands and the function of DNA methylation. Nature. 1986;321(6067):209-13.
52. Deaton AM, Bird A. CpG islands and the regulation of transcription. Genes Dev. 2011;25(10):1010-22.
53. Bird AP. DNA methylation and the frequency of CpG in animal DNA. Nucleic Acids Res. 1980;8(7):1499-504.
54. Auton A, McVean G. Estimating recombination rates from genetic variation in humans. Methods Mol Biol. 2012;856:217-37.
55. Auton A, Rui Li Y, Kidd J, Oliveira K, Nadel J, Holloway JK, et al. Genetic Recombination Is Targeted towards Gene Promoter Regions in Dogs. PLoS Genet. 2013;9(12):e1003984.
56. Cohen NM, Kenigsberg E, Tanay A. Primate CpG islands are maintained by heterogeneous evolutionary regimes involving minimal selection. Cell. 2011;145(5):773-86.
57. Hoge C, de Manuel M, Mahgoub M, Okami N, Fuller Z, Banerjee S, et al. Patterns of recombination in snakes reveal a tug-of-war between PRDM9 and promoter-like features. Science. 2024;383(6685):eadj7026.
58. Joseph J, Prentout D, Laverré A, Tricou T, Duret L. High prevalence of Prdm9-independent recombination hotspots in placental mammals. bioRxiv. 2023:2023.11.17.567540.
59. Kawakami T, Mugal CF, Suh A, Nater A, Burri R, Smeds L, et al. Whole-genome patterns of linkage disequilibrium across flycatcher populations clarify the causes and consequences of fine-scale recombination rate variation in birds. Mol Ecol. 2017;26(16):4158-72.
60. Schield DR, Pasquesi GIM, Perry BW, Adams RH, Nikolakis ZL, Westfall AK, et al. Snake Recombination Landscapes Are Concentrated in Functional Regions despite PRDM9. Mol Biol Evol. 2020;37(5):1272-94.
61. Singhal S, Leffler EM, Sannareddy K, Turner I, Venn O, Hooper DM, et al. Stable recombination hotspots in birds. Science. 2015;350(6263):928-32.
62. Gardiner-Garden M, Frommer M. CpG islands in vertebrate genomes. J Mol Biol. 1987;196(2):261-82.
63. Leitwein M, Wellband K, Cayuela H, Le Luyer J, Mohns K, Withler R, et al. Strong Parallel Differential Gene Expression Induced by Hatchery Rearing Weakly Associated with Methylation Signals in Adult Coho Salmon (O. kisutch). Genome Biol Evol. 2022;14(4).
64. Sigrist CJ, Cerutti L, Hulo N, Gattiker A, Falquet L, Pagni M, et al. PROSITE: a documented database using patterns and profiles as motif descriptors. Brief Bioinform. 2002;3(3):265-74.
